# Identification of radical SAM enzymes responsible for the methylation and desaturation of archaeal lipids and an AttH hydratase mediating hydroxy-GDGT biosynthesis

**DOI:** 10.1101/2025.10.20.683597

**Authors:** Andy A. Garcia, Grayson L. Chadwick, Dipti D. Nayak, Paula V. Welander

## Abstract

Lipid biosynthesis in archaea is highly dependent upon radical (*S*)-adenosyl methionine (rSAM) enzymes. The formation of their membrane-spanning lipids, known as glycerol dibiphytanyl glycerol tetraethers (GDGTs), is catalyzed by the rSAM tetraether synthase and the subsequent cyclization of their lipid tails is performed by the B12-binding rSAM (B12-rSAM) GDGT ring synthase. Further, GDGT tails are cross-linked by the rSAM GMGT synthase and then methylated by the B12-rSAM GMGT methylase. Here, utilizing three archaeal model organisms – *Sulfolobus acidocaldarius*, *Thermococcus kodakarensis*, and *Methanosarcina acetivorans* – we expand this repertoire of rSAM enzymes further, identifying two B12-rSAM GDGT methylases and a novel B12-rSAM GDGT desaturase. We also extend beyond the rSAM enzyme superfamily, identifying an AttH-like hydratase mediating hydroxy-GDGT (OH-GDGT) biosynthesis. Specifically, we identify two GDGT methylases in *S. acidocaldarius* and *Thermococcus aggregans* which possess different substrate specificities. While GDGT methylation is generally low in archaea, we observe high levels of lipid methylation in response to penicillin G and hexanoic acid amphiphile exposure, suggesting a role for lipid methylation in response to membrane-destabilizing chemical agents. Utilizing heterologous expression in *M. acetivorans* and *T. kodakarensis*, we uncover a novel B12-rSAM enzyme from *Candidatus* Bathyarchaeota B1_G15 with unusual tetraether desaturase (Ted) activity that seemingly reverses previous saturation by geranylgeranyl reductase, anaerobically forming double bonds in the inert hydrocarbon tails of a GDGT. Finally, we show that a non-rSAM AttH-like hydratase, genomically associated with Ted, is a hydroxy-GDGT synthase (Hgs), putatively hydrating the double bond(s) introduced by Ted to form OH-GDGTs. These results highlight the value of exploring the rSAM enzyme landscape of archaea – revealing diverse and novel rSAM enzyme activities amongst homologous proteins that invoke new paradigms such as reversibility in the archaeal lipid biosynthesis pathway.

## Introduction

Lipids are a class of chemically diverse amphipathic compounds that comprise the cell membranes of life on Earth. Archaea, in particular, utilize a unique set of isoprenoid-based lipids that differ considerably from the fatty acid-based membranes of bacteria and eukaryotes^1,2^. These archaeal lipids are constructed from highly branched hydrocarbon “tails” that are ether-bonded to glycerol-1-phosphate to form diether lipids known as archaeols^1^. This contrasts with the linear hydrocarbon tails, ester bonds, and glycerol-3-phosphate commonly found in bacterial and eukaryotic membrane lipids. Notably though, both archaea and bacteria/eukaryotes are known to chemically modify their lipids as an adaptive response to environmental challenges (e.g. high temperature)^3^. Such modifications alter the structure of the lipid and work as a mechanism to adjust the biophysical properties of the membrane to function optimally under different environmental conditions.

These lipid modifications and the enzymes that perform them commonly differ between archaea and bacteria/eukaryotes. In response to elevated growth temperatures^4–6^, archaea are known to employ the radical (*S*)-adenosyl methionine (rSAM) enzyme tetraether synthase (Tes)^7,8^ to fuse their bilayer/diether lipids into a monolayer of membrane-spanning lipids known as glycerol dibiphytanyl glycerol tetraethers (GDGTs)^9^. GDGTs can be further modified by introducing cyclopentane rings and/or crosslinks into the lipid tails by the rSAMs GDGT ring synthase (Grs)^10^ and GMGT synthase (Gms)^11,12^, respectively. These three modifications increase the packing efficiency of the lipids, resulting in a denser membrane that is more rigid and less permeable, thereby preserving the membrane’s integrity under harsher conditions^13,14^. While some bacteria are known to make analogous membrane-spanning lipids (e.g. branched GDGTs^15^), this appears to be restricted to certain phylogenetic groups (e.g. Acidobacteria and Thermotogales)^16–20^, albeit with a potentially wider distribution than previously appreciated^21^. In contrast, bacteria and eukaryotes typically respond to elevated growth temperatures by increasing the chain length of their lipid tails through the regulation of acyltransferases, ketosynthases, and thioesterases that control chain elongation and termination^22,23^ or by adjusting ratios of *iso* and *anteiso* branched chain fatty acids^24^. For cold stress, bacteria/eukaryotes commonly produce unsaturated fatty acids by either desaturation or dehydration of their lipid tails^3,25^. Desaturation is oxygen-dependent and is performed by fatty acid desaturases which utilize a diiron cofactor and O_2_ to abstract hydrogen atoms from the saturated lipid tails, forming double bonds^25^. In contrast, oxygen-independent dehydratase enzymes such as FabA and FabZ dehydrate the hydroxy-functionalized intermediates of fatty acid biosynthesis to form double bonds^26^. These unsaturated fatty acids disrupt lipid packing and prevent membrane over-rigidification/gel-phase transitions^3,24,25,27^. Archaeal responses to lower growth temperatures are less well characterized but may involve lipid methylation^28^ and hydroxylation^29,30^ in addition to regulating the saturation state of their lipids^31^.

These modifications result in a large chemical diversity of lipids. In archaea, GDGTs are known to possess a number of modifications such as cyclization, cross-linking to form glycerol monoalkyl glycerol tetraethers (GMGTs), methylation, hydroxylation, and unsaturation, giving rise to a variety of structurally distinct tetraether lipids^32^. While the biosynthesis and physiological roles of some of these modifications have been well characterized (e.g. cyclization and cross-linking)^10,11,33^, the formation and roles of a large portion of the chemical diversity of GDGTs remain enigmatic. Of particular interest are GDGT modifications associated with mesophilic environments. While GDGTs and several of their modifications are traditionally found in extreme localities and have been shown to be important for survival under these regimes^33–35^, archaea are known to inhabit a plethora of more modest habitats including the deep ocean^36^, soils^37,38^, sediments^39^, and the human microbiome^40^. It is therefore of interest to understand why archaea synthesize GDGTs under less extreme conditions and how they may modify them to function optimally under these different conditions.

One of the most notable classes of GDGTs associated with these mesophilic environments are the hydroxy-GDGTs (OH-GDGTs) which possess 1-2 tertiary alcohol groups at the C-3 position(s) in their biphytanyl lipid tails^41^. These lipids were first identified in environmental samples from marine sediments^41^ and have since been shown to occur in widespread mesophilic marine and lacustrine habitats^29^. The abundance of OH-GDGTs increases with decreasing temperature and increasing latitude in marine settings^41,42^, and they form the basis of the %OH and TEX_86_^OH^ temperature proxies^29,43^ used to calculate sea-surface temperatures based upon the hydroxylation state of GDGTs alone or in combination with their non-hydroxylated and cyclized counterparts, respectively. Several archaeal clades involved in important biogeochemical cycles have been found to produce OH-GDGTs including mesophilic members of the ammonia-oxidizing Thaumarchaeota (now Nitrososphaerota) including *Nitrosopumilus maritimus*^44^, mesophilic anaerobic methanotrophic (ANME) archaea such as *Ca. Methanoperedens* sp. strain BLZ2^45^, the thermophilic methanogen *Methanothermococcus thermolithautotrophicus*^41,46^, and putatively in mesophilic methylotrophic methanogens^47^. While the enzyme(s) responsible for the formation of OH-GDGTs are unknown, monooxygenases such as P450 hydroxylase enzymes are known to install hydroxyl groups in some eukaryotic and bacterial lipids including sterols and acylceramides^48–50^. While these oxygen-requiring enzymes could be responsible for OH-GDGT production in the aerobic Nitrososphaerota, they cannot explain OH-GDGT biosynthesis in the anaerobic ANME and methanogenic archaea.

Recently, a phytoene desaturase family protein, here termed hydroxy-archaeol synthase (Has), from *Methanosarcina acetivorans* was shown to use water to form analogous hydroxylated diether lipids^51^ known as hydroxy-archaeols (OH-archaeols) which are widespread among methanogenic and methanotrophic archaea^45,52–54^. This enzyme hydrates the C2-C3 double bond in unsaturated archaeol, forming OH-archaeols during heterologous expression in an *Escherichia coli* strain that produces fully unsaturated archaeol, known as digeranylgeranylglyceryl phosphate (DGGGP)^51^. However, it is still unclear if archaeol hydroxylation is the native activity of the Has protein when expressed in *M. acetivorans*, if this protein can also hydroxylate unsaturated GDGTs to form OH-GDGTs or, alternatively, if OH-archaeols are fused by the Tes enzyme directly to yield OH-GDGTs. Further, a number of observations complicate the association of OH-GDGTs with low temperatures such as: 1) the production of OH-GDGTs by thermophilic archaea^41,46^, 2) no increase in OH-GDGT levels in response to lowering growth temperatures in pure cultures of archaea^55^, and 3) variables other than temperature (e.g. nutrient concentrations and water stratification) influencing OH-GDGT abundance in the environment^56^. Taken together, these observations call for a better understanding of the biosynthesis and physiological roles of OH-GDGT lipids.

Another GDGT lipid class commonly found in mesophilic environments are the unsaturated GDGTs (uns-GDGTs) which possess one or more double bonds within their lipid tails^57^. Uns-GDGTs with 1-6 double bonds are found in mesophilic marine and lacustrine environments and are particularly abundant in euxinic settings^57–59^. Few cultured archaea have been shown to produce uns-GDGTs, but GDGTs with 1-4 double bonds have been tentatively identified at trace levels in the hyperthermophilic archaeon *Thermococcus kodakarensis*^60^. The biosynthesis of these lipids could theoretically take place via two routes: 1) fusion of partially unsaturated archaeol intermediates by Tes or 2) desaturation of GDGTs. The archaeal lipid biosynthesis pathway naturally produces unsaturated intermediates which arise from the unsaturated 5-carbon isoprene building blocks isopentyl pyrophosphate (IPP) and dimethylallyl pyrophosphate (DMAPP) used to construct their lipid tails^61^. The unsaturated intermediate DGGGP possesses 8 double bonds (4 on each tail) and is saturated by the enzyme geranylgeranyl reductase (GGR)^62^. Partially unsaturated archaeols with 1-6 double bonds are commonly detected in lipid extracts of cultured archaea^63,64^, suggesting GGR does not always fully saturate archaeol. It is therefore possible for unsaturated intermediates to be fused by Tes to form uns-GDGTs. Alternatively, saturation by GGR could be reversed by desaturation of a fully saturated GDGT in a strategy analogous to fatty acid desaturation in bacteria and eukaryotes. However, fatty-acid desaturases are oxygen-dependent and uns-GDGTs are common in anaerobic environments, suggesting that if archaeal GDGT desaturation exists, it may occur by a novel O_2_-independent mechanism.

Finally, one last class of understudied GDGTs that may be involved in archaeal adaptation to low/suboptimal growth temperatures are the extra-methylated GDGTS. While GDGTs natively possess eight methyl groups per tail, arising from their isoprene building blocks, 1-3 additional methyl groups can be added at the C-13 position(s) within their biphytanyl tails to form Me-GDGTs^65^ or to the glycerol backbone to form butanetriol dibiphytanyl glycerol tetraethers (BDGTs)^66^. Me-GDGTs have been identified in several environmental samples^67–69^ and in a few cultured archaea at generally low levels, including on acyclic GDGTs in both mesophilic methanogens such as *Methanobrevibacter smithii*^60^ and thermophilic methanogens like *Methanothermobacter thermautotrophicus*^65^ and on cyclic GDGTs in the thermoacidophiles *Sulfolobus acidocaldarius*^65^ and *Thermoplasma acidophilum*^70^. Recent molecular dynamics simulations suggest that GDGT methylation disrupts the packing of archaeal lipids, increasing membrane fluidity, and thus may be an adaptation to growth at lower/suboptimal temperatures^28^. However, few culture studies investigating the methylation patterns of archaeal lipids exist to validate this hypothesis. One study found that GDGT methylation does increase approximately 5-fold from 2% to 10% of tetraethers in response to decreasing growth temperature from 60 °C to 45 °C in *M. thermautotrophicus*, albeit in a single replicate^65^. However, in other studies additional factors such as exposure to the membrane-destabilizing detergent Tween-20^71^ or lower nutrient conditions^72^, appeared to trigger GDGT methylation in the related archaeon *Methanothermobacter marburgensis* and in *M. thermautorophicus*, respectively. Detergents disrupt lipid packing and solubilize/permeabilize membranes^73^, suggesting catastrophic increases in membrane fluidity and complicating the idea that Me-GDGTs would be used to further decrease fluidity. In our previous work, we demonstrate that methylation of a related class of lipids, GMGTs, increases in response to increased growth temperature in *Vulcanisaeta distributa*^11^ and that an rSAM enzyme, GMGT methylase (Gmm), is responsible for this modification. However, Gmm does not robustly methylate regular (un-crosslinked) GDGTs, suggesting other enzymes may be responsible for Me-GDGT production. These scant and seemingly confounding results implore deeper investigations into GDGT methylation in archaea.

Our lack of information on the biosynthetic route of OH-GDGTs, uns-GDGTs, and Me-GDGTs is a major barrier to determining the physiological roles, archaeal sources, and environmental significance of these lipids. While GDGT biosynthesis utilizes radical SAM enzymology in all steps characterized to date, these reactions all involve the formation of new carbon-carbon bonds at inert sp^3^ hybridized centers (dimerization^7,8^, cyclization^10^, crosslinking^11,12^, and methylation^11^ of the hydrocarbon tails), and it is thus unclear if they would be responsible for the introduction of additional modifications which do not involve carbon-carbon bond formation (OH-GDGTs and uns-GDGTs). The rSAM enzyme superfamily, however, is known to possess incredible functional diversity^74,75^. For example, although the B12-binding rSAMs (B12-rSAMs) Grs and Gmm are homologous, they carry out disparate reactions, with Grs catalyzing the cyclization of lipid tails and Gmm catalyzing their methylation. Such diverse activity is typical of the B12-rSAM enzyme subfamily^74^, and members of this subclass carry out several distinct reactions ranging from complex ring formations^76^ and contractions^77^ to various forms of methylation^78,79^ and, interestingly, to seemingly off-target Csp^3^-Csp^3^ desaturation reactions at inert carbon centers during in vitro studies (albeit with no biological example of such activity)^80^. We thus hypothesized that homologs of the B12-rSAMs Gmm and Grs might be responsible for the remaining unknown steps of GDGT biosynthesis.

In this study, we both confirm and refute this hypothesis, identifying B12-rSAM enzymes which methylate and desaturate GDGTs to form Me-GDGTs and uns-GDGTs, respectively, and a non-rSAM AttH-like hydratase that forms OH-GDGTs. More specifically, we identify Gmm homologs that function as 1) cyclized GDGT methylases (Cgm), preferentially methylating the highly cyclized GDGTs-4, 5, and 6 over GDGT-0, or as 2) promiscuous GDGT methylases (Pgm), robustly methylating GDGT-0 as well as bilayer lipids to a lesser extent. Furthermore, we glean insights into a potential physiological role for Me-GDGTs in archaea, identifying a unique amphiphile-induced lipid methylation response in *S. acidocaldarius* and *T. aggregans* that is triggered during exposure to penicillin G and hexanoic acid. We also demonstrate that a distant Gmm homolog from a *Candidatus* Bathyarchaeon possesses tetraether desaturase (Ted) activity rather than methylase activity, forming double bonds within inert GDGT hydrocarbon tails – the first example of a lipid desaturase in archaea and the first biological example of Csp^3^-Csp^3^ desaturase activity at inert carbon centers in the radical SAM superfamily. Finally, we identify a hydroxy-GDGT synthase (Hgs) that co-occurs with Ted in archaeal genomes and putatively hydrates the double bonds introduced by Ted to form OH-GDGTs. Altogether, this study validates the importance of rSAM enzymes to archaeal GDGT biosynthesis, uncovers novel rSAM activities, and introduces new enzyme families involved in GDGT modification.

## Results and Discussion

### Identification of a cyclized GDGT methylase in *S. acidocaldarius*

GDGTs that possess an extra methyl group in their biphytanyl lipid tails (Me-GDGTs) have been identified in a number of (thermo)acidophilic archaea including *Sulfolobus acidocaldarius*^65^*, Thermoplasma acidophilum*^81^, *Ferroplasma acidarmanus*^70^, and *Ferroplasma acidiphilum*^70^. Of these archaea, Me-GDGTs have been most well characterized in *S. acidocaldarius*, where the additional methyl group was tentatively found to occur at C-13, deep within the hydrocarbon tail^65^. In the same study, GDGTs containing 4, 5, and 6 rings (GDGT-4, 5, and 6) were found to be enriched in this extra methylation compared to the lower cyclized GDGTs and acyclic GDGT-0. However, these Me-GDGTs comprised <2% of total core lipids and are thus often overlooked in studies of the *S. acidocaldarius* lipidome with the 2015 Knappy et al study^65^ being the first and only such investigation to our knowledge.

To better understand GDGT methylation in *S. acidocaldarius*, we first sought to identify and characterize Me-GDGT production in *S. acidocaldarius* throughout its growth phases. We grew *S. acidocaldarius* in triplicate for 6 days, harvesting samples every 24 hours and analyzing core lipid extracts of these cultures via liquid chromatography-mass spectrometry (LC-MS) in reverse phase. We detected generally low abundances of Me-GDGTs in all samples, primarily observing mono-methylated derivatives of the highly cyclized GDGTs-4, 5, and 6: Me-GDGT-4, Me-GDGT-5, and Me-GDGT-6 (Fig. 1A-B, S1-2). We calculated the methylation index of GDGTs-4, 5, and 6 (MI_GDGT-4,5,6_) as the weighted average number of extra methyl groups on GDGTs-4, 5, and 6 together and found that GDGT methylation gradually increases during growth. Specifically, the methylation index increased from an MI_GDGT-4,5,6_ = 0.01 ± 0.00 on day 1 to an MI_GDGT-4,5,6_ = 0.09 ± 0.01 on day 3 to an MI_GDGT-4,5,6_ = 0.19 ± 0.01 on day 6 (Fig. S2).

**Figure 1.**
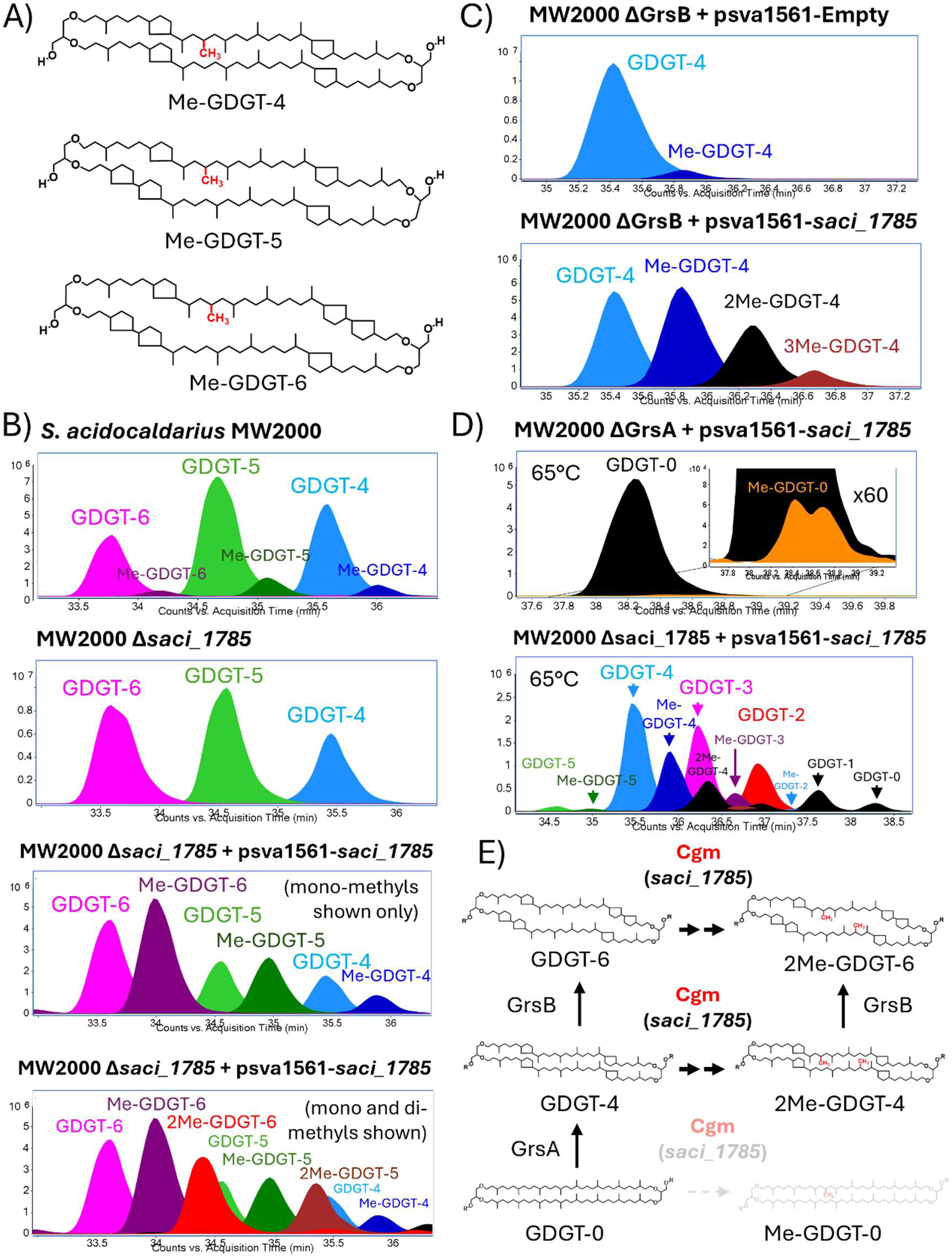
*Saci_1785* encodes a cyclized GDGT methylase (Cgm) in *S. acidocaldarius*. A) Structures of the most abundant methylated GDGTs (Me-GDGTs) found in *S. acidocaldarius* under standard growth conditions. B) Representative, overlayed, extracted ion chromatograms (EIC) (*m/z* values discussed in the supplement*) of core lipids extracted from acid hydrolyzed biomass of the parental *S. acidocaldarius* strain MW2000, the *saci_1785* deletion strain (Δ*saci_1785*), and the *saci_1785* complementation strain (Δ*saci_*1785 + psva1561-*saci_1785*). The parental strain produces abundant GDGTs with 4, 5, and 6 cyclopentane rings (GDGT-4, GDGT-5, and GDGT-6) and only produces small amounts of their methylated derivatives, Me-GDGT-4, Me-GDGT-5, and Me-GDGT-6. In *Δsaci_1785,* deletion of *saci_1785* results in the complete loss of methylated lipids, demonstrating that this locus is required for GDGT methylation. Complementation of Δ*saci_1785* with psva1561-*saci_1785* restores GDGT methylation, producing both mono- and di-methylated GDGTs-4, 5, and 6. C) Representative, overlayed EICs showing that overexpression of *saci_1785* in the *grsB* deletion mutant (Δ*grsB* + psva1561-*saci_1785*) results in robust methylation of GDGT-4 producing mono-, di-, and tri-methylated derivatives. The empty plasmid control strain (Δ*grsB* + psva1561) only produces trace amounts of Me-GDGT-4. D) Representative, overlayed EICs showing *saci_1785* overexpression in the *grsA* deletion mutant (Δ*grsA* + psva1561-*saci_1785*) or in the *saci_1785* deletion mutant *(Δsaci_1785* + psva1561-*saci_1785)* at 65°C. At this lower growth temperature, only trace GDGT-0 methylation is observed in the *ΔgrsA* background. However, overexpression in the Δ*saci_1785* complementation strain at 65°C results in robust GDGT-4 methylation suggesting that GDGT-0 is not the native substrate of Saci_1785. E) Saci_1785 is a cyclized GDGT methylase (Cgm) that preferentially methylates highly cyclized GDGTs over GDGT-0. GrsA cyclizes GDGT-0, forming GDGT-4 in high amounts. GDGT-4 may then be mono- and di-methylated by Cgm (Saci_1785). Mono- and di-methylated derivatives of GDGTs-5 and 6 can theoretically be formed by two routes: 1) direct methylation of GDGT-5/6 by Cgm or 2) cyclization of Me-GDGT-4 by GrsB.

To identify a candidate protein responsible for catalyzing cyclized GDGT methylation, we searched the *S. acidocaldarius* genome for homologs of the B12-binding radical SAM (B12-rSAM) GMGT methylase (Gmm) which performs a similar reaction on the monoalkyl tail of cross-linked GDGTs, known as GMGTs^11^. A BLASTP search revealed that the locus *saci_1785* encodes a Gmm homolog with an e-value of 7e^-72^ and 34% identity to the *Thermococcus guaymasensis* Gmm. Further, the Position of Proteins in Membranes Webserver (PPM 3.0)^82,83^ predicts that Saci_1785 is a C-terminally bound peripheral membrane protein with a robust ΔG_transfer_ value of -14.8 ± 0.5 kcal/mol^82,83^ (Fig. S3, Table S1). As *S. acidocaldarius* does not produce GMGT lipids, we hypothesized that *saci_1785* instead encodes a methylase that specifically methylates cyclized GDGTs. To test this hypothesis, we constructed an in-frame, markerless deletion mutant of *saci_1785* (Δ*saci_1785*) in *S. acidocaldarius* strain MW2000. Lipid analysis of Δ*saci_1785* revealed the complete loss of Me-GDGTs, demonstrating *saci_1785* is required for GDGT methylation in *S. acidocaldarius* (Fig. 1B, S4-5). Furthermore, complementation of the strain via re-introduction of *saci_1785* on the autonomously replicating plasmid psva1561^84^, restored Me-GDGT production at levels 10-fold higher than the parental strain (MI_GDGT-4,5,6_ = 0.92 ± 0.07 vs. MI_GDGT-4,5,6_ = 0.09 ± 0.01), resulting in the production of both mono- and di-methylated GDGTs-4, 5, and 6 (Fig. 1B, S4-5). Interestingly, even under these high methylation conditions, we did not detect any methylation on GDGT-0 or GDGT-1 and only detected low levels of methylation on GDGT-2 (MI_GDGT-2_ = 0.07 ± 0.00) and GDGT-3 (MI_GDGT-3_ = 0.24 ± 0.01), suggesting that the enzyme may prefer to work on cyclized GDGTs of increasing ring number. However, GDGTs-0, 1, 2, and 3 together only comprised 3.3% ± 0.7% of total core lipids, which could suggest that the methylated derivatives of GDGTs-0, 1, 2, and 3 may be present but below detection limits. It might also be possible that the more abundant highly cyclized GDGTs simply outcompete the lower abundance lipids for the *saci_1785* methylase.

To further interrogate the substrate preference of the methylase, we overexpressed *saci_1785* on the psva1561 plasmid in a GDGT ring synthase B (GrsB) deletion mutant (Δ*grsB*) which only makes GDGTs-0, 1, 2, 3, and 4^10^. Overexpression of *saci_1785* in Δ*grsB* resulted in the robust production of mono, di, and tri-methylated GDGT-4 with an MI_GDGT-4_ = 1.04 ± 0.01 compared to an MI_GDGT-4_ = 0.06 ± 0.0 in the empty plasmid control strain (Fig. 1C, S4-5). GDGTs-0, 1, 2, and 3 together comprised a slightly larger percentage of total core lipids at 5.3% ± 0.7% but we again did not detect any Me-GDGT-0 or Me-GDGT-1 and only observed low GDGT-2 methylation (MI_GDGT-2_ = 0.08 ± 0.03). GDGT-3 methylation increased to an MI_GDGT-3_ = 0.42 ± 0.01 but was still considerably less than that for GDGT-4.

To better determine if the methylase can work on acyclic GDGT-0, we overexpressed *saci_1785* in a GDGT ring synthase A (GrsA) deletion mutant (Δ*grsA*) which makes GDGT-0 as its primary core lipid^10^. The *grsA* deletion mutant is temperature sensitive so expression was carried out at 65°C rather than the optimal 75°C. Overexpression of *saci_1785* in the Δ*grsA* background resulted in only trace methylation of GDGT-0 with an MI_GDGT-0_ = 0.02 ± 0.00 (Fig. 1D, S4-5). In comparison, when the complementation strain, Δ*saci_1785* + psva1561-*saci_1785*, is grown at the same temperature, the cyclized GDGT methylation index is over 30-fold higher with an MI_GDGT-4,5,6_ = 0.63 ± 0.05 (Fig. 1D, S4-5). Additionally, at 65°C we observed lower levels of GDGT cyclization. At this lower temperature, GDGTs-0, 1, 2, and 3 together composed 44.2% ± 1.7% of core lipids in the complementation strain while GDGT-4 comprised 53.9% ± 1.7% of core lipids, allowing us to more reliably compare their methylation levels (Fig. S4). Under these conditions, we observed a stepwise decrease in methylation with decreasing ring number, from an MI_GDGT-4_ = 0.64 ± 0.05 to an MI_GDGT-3_ = 0.22 ± 0.00 to an MI_GDGT-2_ = 0.05 ± 0.00 and no methylation of GDGT-1 and GDGT-0, consistent with the observations at 75°C (Fig. S5). Taken together, these data demonstrate that *saci_1785* encodes a cyclized GDGT methylase (Cgm) that prefers to methylate highly cyclized GDGTs over lower cyclized and acyclic GDGT-0 (Fig. 1E).

### Identification of a promiscuous GDGT methylase from *T. aggregans*

To better understand the distribution of cyclized GDGT methylation in archaea, we searched the National Center for Biotechnology Information (NCBI) protein database for Cgm homologs using Saci_1785 as our BLASTP search query (e-value ≤ 1e^-99^, >495 amino acids). We found that Cgm homologs are phylogenetically restricted to a few archaeal clades within the TACK and Euryarchaeota, including Thermoplasmata (31% of hits), Sulfolobaceae (14%), Fervidicoccaceae (13%), non-ammonia-oxidizing members of the Nitrososphaerota (11%), Thermococci (8%), Desulfurococcales (5%), and Marsarchaeota (4%) (Fig. S6). Construction of a phylogenetic tree of these Cgm homologs revealed the presence of two distinct Cgm clades (Fig. 2A). Clade 1 possesses Cgm proteins from *S. acidocaldarius* and other (thermo)acidophilic archaea within Sulfolobaceae, Thermoplasmata, and Marsarchaeota. In contrast, clade 2 consists of Cgm homologs from the more neutrophilic Thermococci and Fervidicoccaceae. While archaea within the acidophilic Thermoplasmata and Sulfolobaceae have been shown to produce abundant cyclized GDGTs^32^, the neutrophilic Thermococci primarily produce GDGT-0 and the bilayer lipid archaeol as their major membrane lipids, while cyclized GDGTs are typically absent or only present at trace levels^85^. This suggests that the Cgm homologs found in Thermococci might have greater substrate promiscuity than Saci_1785 and may be able to more robustly methylate acyclic GDGT-0. In support of this, previous lipid analysis of the only three cultured *Thermococcus* species with a Cgm homolog (*Thermococcus aggregans*, *Thermococcus litoralis*, and *Thermococcus sibiricus*) found either no or trace cyclized GDGTs^85^. Further, while *T. aggregans* and *T. litoralis* have the genomic potential to cyclize GDGTs, possessing GDGT ring synthase genes (*F865_RS03120* and *OCC_RS06675*, respectively), *T. sibiricus* does not, suggesting Thermococci Cgm homologs can function in the absence of any cyclized GDGTs.

**Figure 2.**
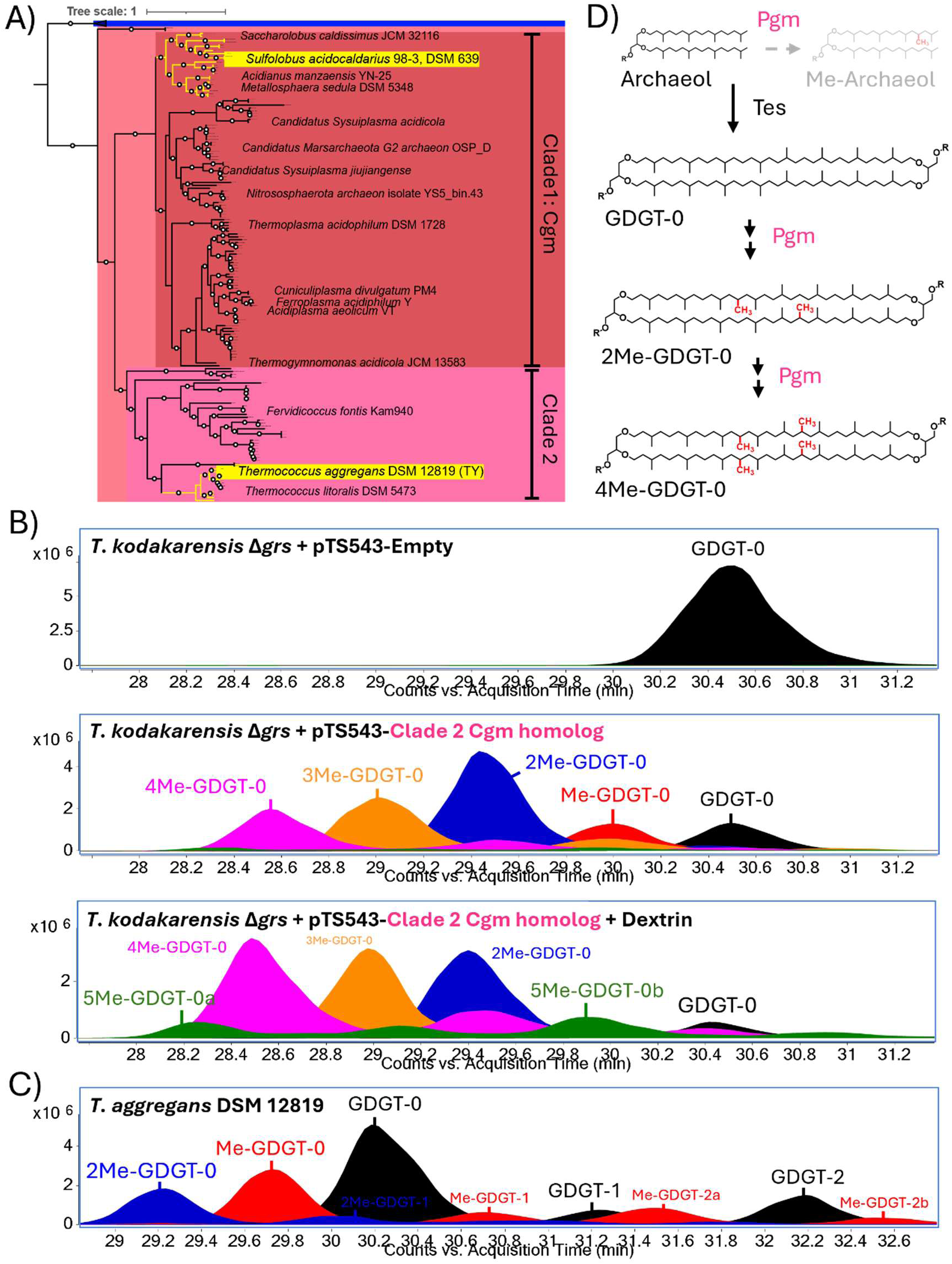
A Cgm homolog from *Thermococcus aggregans* encodes a promiscuous GDGT methylase (Pgm). A) A phylogenetic tree of Cgm proteins. Gmm is used as an outgroup (blue). Cgm homologs are shown in red/pink and can be divided into two major clades with strong bootstrap support (≥0.9). Clade 1 possesses “canonical” Cgm proteins, including Saci_1785 from *S. acidocaldarius*, and Cgm proteins from other (thermo)acidophilic archaea within Sulfolobaceae (branches highlighted in yellow at the top of clade 1), Thermoplasmata, Nitrososphaerota, and Marsarchaeota. Clade 2 possesses divergent Cgm homologs from the more neutrophilic archaea within Fervidococcacaea and Thermococci (branches highlighted in yellow at the bottom of clade 2), including *T. aggregans*. Bootstrap values ≥ 0.9 are shown with open circles. The tree scale represents one change per amino acid site. B) Representative, overlayed extracted ion chromatograms (EICs) of core lipids extracted from acid-hydrolyzed biomass of the *T. kodakarensis* control strain (Δ*grs* + pTS543-Empty) and *T. kodakarensis* heterologously expressing the *T. aggregans* clade 2 Cgm homolog under basal (no dextrin) and inducing conditions. The control strain only makes GDGT-0 and no detectable Me-GDGTs. In contrast, the strain expressing the clade 2 Cgm homolog produces mono-, di-, tri-, and tetra-methylated GDGT-0 as its major core lipids under basal and inducing conditions. This contrasts with the Cgm enzyme from *S. acidocaldarius* which can only produce trace levels of Me-GDGT-0. Increasing expression of the clade 2 Cgm homolog by growing the strain in the presence of the inducer, dextrin, results in even higher levels of Me-GDGTs, including the production of unusual penta-methylated GDGT-0 (5Me-GDGT-0). C) Representative, overlayed EIC of core lipids extracted from acid -hydrolyzed biomass of *T. aggregans* DSM 12819 grown under standard conditions. *T. aggregans* shows robust native production of mono- and di-methylated GDGT-0, demonstrating that the heterologous activity of the clade 2 Cgm homolog in *T. kodakarensis* is consistent with that activity seen in the native host *T. aggregans*. D) Clade 2 Cgm proteins are promiscuous GDGT methylases (Pgm) capable of methylating both bilayer and monolayer lipids and producing a range of poly-methylated products.

To determine if Cgm homologs from Thermococci can methylate GDGT-0, we expressed the *T. aggregans* homolog (7e^-123^ e-value and 40% identity to Cgm in *S. acidocaldarius*) in a *Thermococcus kodakarensis grs* deletion strain (Δ*grs*/AL010) that produces GDGT-0 and archaeol as its major core lipids^35^. Specifically, we placed the *T. aggregans* Cgm homolog under the control of the dextrin-semi-inducible promoter pfba^86^ on the self-replicating plasmid pTS543^87^ and then introduced this plasmid into *T. kodakarensis* for heterologous expression in triplicate and analyzed the core lipids of this strain via LC-MS in reverse phase. This strain produced abundant Me-GDGTs even under basal expression conditions (no dextrin), converting >90% of GDGT-0 into its mono- (9.9% ± 0.4% of GDGTs), di- (32.8% ± 0.7%), tri- (25.0% ± 0.5%), and even tetra-methylated (18.8% ± 0.2%) derivatives (Me-GDGT-0, 2Me-GDGT-0, 3Me-GDGT-0, and 4Me-GDGT-0). The empty plasmid control strain had no detectable GDGT methylation (Fig. 2B, Fig. S7&10). Upon increasing expression of the gene during growth on dextrin, we found an increase in the MI_GDGT-0_ from 2.48 ± 0.02 (basal) to 3.37 ± 0.03 (induced) with 4Me-GDGT-0 being the dominant GDGT lipid under the inducing conditions (32.4% ± 0.9%). These data demonstrate that the *T. aggregans* Cgm homolog is a GDGT methylase protein that is functionally distinct from Saci_1785, robustly methylating acyclic GDGT-0 (Fig. 2B, S8&10).

Under inducing conditions, we also observed the production of a variety of penta-methylated GDGT-0 isomers (5Me-GDGT-0), together comprising 13.2% ± 0.6% of core lipids (Fig. 2B, S8, S9A, S10A). GDGTs possess four equivalent C-13 positions on their biphytanyl lipid tails and as such, each sequential methylation is presumed to occur on a “free” unmethylated C-13 atom, theoretically yielding GDGTs with up to four additional methyl groups. However, the detection of penta-methylated GDGTs (as well as trace hexa- and hepta-methylated GDGTs) suggests that the methylase can 1) doubly methylate the C-13 carbon atom, forming a quaternary carbon center (such as by the CysS B12-rSAM^88^), 2) methylate the previously introduced methyl group to form an ethyl group (such as by the TokK^89^ and TmoD^90^ B12-rSAMs), and/or 3) methylate at a position other than C-13 (Fig. S9A). More in-depth chemical analyses are needed to distinguish between these scenarios, but they demonstrate the promiscuous activity of the GDGT methylase from *T. aggregans* compared to the *S. acidocaldarius* Cgm. Interestingly, analysis of the bilayer lipids of this strain revealed moderate levels of mono- and di-methylated archaeols as well (∼25% bilayer lipids together), compared to no archaeol methylation in the empty plasmid control strain, further highlighting the promiscuous activity of the enzyme (Fig. S9B, S10B&D).

To investigate the potential for cyclized GDGT methylation by the *T. aggregans* methylase, we co-expressed this protein with *gdgt ring synthase A* from *T. kodakarensis* (TK0167) which synthesizes the cyclized GDGTs-1 to 4. Lipid analysis of this strain revealed high levels of methylation on both GDGT-0 and its cyclized derivatives, with an MI_GDGT-0_ = 2.29 ± 0.01 compared to a similar MI_GDGT-4_ = 1.93 ± 0.01 (Fig. S9C, S10C-D). This suggests that the *T. aggregans* methylase works robustly on both acyclic and cyclic GDGTs and/or that the methylated GDGT-0 derivatives are cyclized by Grs, also yielding cyclized, methylated GDGTs. Given that heterologous expression of the *T. aggregans* methylase results in the production of diverse products (Me-archaeols, mono-, di-, tri-, and tetra-Me-GDGT-0, multiple unusual penta-methylated GDGT-0 isomers, and methylated cyclized GDGTs), we propose to call the Cgm homologs from Thermococci archaea promiscuous GDGT methylases (Pgm) (Fig. 2D).

Finally, to ensure that GDGT-0 methylation by Pgm reflects its native activity, we analyzed core lipids from the native host *T. aggregans* at four timepoints representing exponential, early stationary, late stationary, and death phases in triplicate to search for natural Me-GDGT-0 production. We identified robust production of mono- and di-methylated GDGT-0 in *T. aggregans* which increased gradually throughout the growth phases, as in *S. acidocaldarius*, from an MI_GDGT-0_ = 0.44 ± 0.01 in exponential phase to an MI_GDGT-0_ = 0.68 ± 0.02 in late stationary and death phase (Fig. 2C, S11-12). These data demonstrate that Pgm proteins from Thermococci archaea robustly methylate GDGT-0, leading to diverse, multi-methylated acyclic and cyclic GDGT lipids.

### Characterization of an amphiphile-induced GDGT methylation response in *S. acidocaldarius and T. aggregans*

While methylation is a structurally simpler modification compared to those typically found in archaeal GDGTs (e.g. rings and crosslinks), the methylation of lipids can significantly alter membrane properties and has been shown to play important physiological roles in a variety of microorganisms^91–95^. GDGT methylation is generally low in archaea, including in *S. acidocaldarius*, suggesting Me-GDGTs are not as important for growth under standard laboratory conditions and that they may instead have a more specialized role in the cell. To gain insight into the potential roles of Me-GDGTs in archaea, we investigated the genomic context of *cgm* and *pgm* genes using the Enzyme Function Initiative Genome Neighborhood Tool (EFI-GNT) to generate a genome neighborhood network (GNN) of GDGT methylase genes (see methods). This GNN revealed that *cgm*/*pgm* commonly co-occur (≥25%) with genes coding for the protein families PF07690 (56%), PF00005 (27%), and PF00501 (25%) which consist of major facilitator superfamily (MFS) transporters, ABC transporters, and AMP-binding enzymes, respectively.

A closer inspection of these MFS transporters and AMP-binding enzymes revealed that they are putatively involved in the export and degradation of aromatic compounds and fatty acids. In Sulfolobaceae, *cgm* often occurs next to 1) genes encoding the medium-chain-fatty-acid-CoA synthetase AlkK (PF00501) which activates fatty acids for degradation via beta-oxidation^96–100^ and 2) a variety of gene clusters encoding proteins responsible for the degradation of diverse aromatic compounds such as phenylacetic acid, ethylbenzene, benzylformate, and caffeic acid (Fig. S13). In *S. acidocaldarius*, *cgm* lies next to both *alkK* and a gene cluster of phenylacetic acid (paa) degradation genes including 1) *paaK*^101^ (PF00501) which activates paa for aerobic degradation and 2) penicillin G amidase which is responsible for freeing paa from a variety of naturally occurring esters and amides (e.g. penicillin)^102^. In other members of Sulfolobaceae, *cgm* occurs next to genes encoding other aromatic compound degradation pathways, including oxygen-independent degradation by ethylbenezene dehydrogenase^103^ in *Saccharolobus caldissimus* or oxygen-independent, electron bifurcating degradation by caffeyl-CoA reductase^104^ in *Metallosphaera sedula* (Fig. S13). In the more distantly related Thermoplasmata, *cgm* also commonly occurs next to fatty acid and aromatic compound degradation genes. For example, in *Ferroplasma acidiphilum*, *cgm* occurs near a putative long-chain-alcohol dehydrogenase gene and a gene cluster encoding proteins (e.g. 4-hydroxybenzoyl-acetyl-CoA synthetase (PF00501)) that are responsible for the degradation of tyrosine and a variety of other hydroxy-benzoate compounds^105^. Cgm often co-occurs with a putative aromatic compound MFS exporter^106,107^ (PF07690) in both Sulfolobacaea and Thermoplasmata, including *M. sedula* and in *Candidatus* Sysuiplasma superficiale where the MFS exporter gene also clusters with *bldR2*, a gene encoding an aromatic aldehyde responsive MarR transcription factor from the archaeon *Saccharolobus solfataricus*^106,108^. Similarly, *pgm* also clusters with genes involved in the degradation of aromatic compounds such as 4-oxalocrotonate tautomerase^109^ (4-OT) whose coding sequence overlaps with *pgm* in *T. aggregans* (Fig. S13). In addition to 4-OT, a hydrophobe-amphiphile transporter (HAE3 family) and an aromatic amino acid responsive MarR transcription factor (TK0169-related)^110^ also occur next to *pgm* in *T. aggregans*.

Fatty acids and functionalized aromatic compounds are amphipathic, possessing hydrophobic and hydrophilic chemical moieties. Such compounds are known to intercalate into the cell membrane, disrupting lipid packing and destabilizing the membrane^111,112^. The localization of *cgm* and *pgm* genes next to genes involved in the efflux and degradation of fatty acids and aromatic compounds suggests that Me-GDGTs may play a role in cellular responses to these membrane-destabilizing chemical agents. To interrogate this, we grew *S. acidocaldarius* and *T. aggregans* in the presence of penicillin G or hexanoic acid (a medium chain fatty acid) in triplicate for 6 days. Growth was monitored and samples were taken every 24 hours for core lipid analysis via LC-MS chromatography in reverse phase. The two archaea exhibited different sensitivities to these compounds - *S. acidocaldarius* growth was impaired by the presence of either 2mM penicillin G or 3.5mM hexanoic acid while the growth of *T. aggregans* was not affected by even higher levels of either compound (5mM) (Fig. 3A-B).

**Figure 3.**
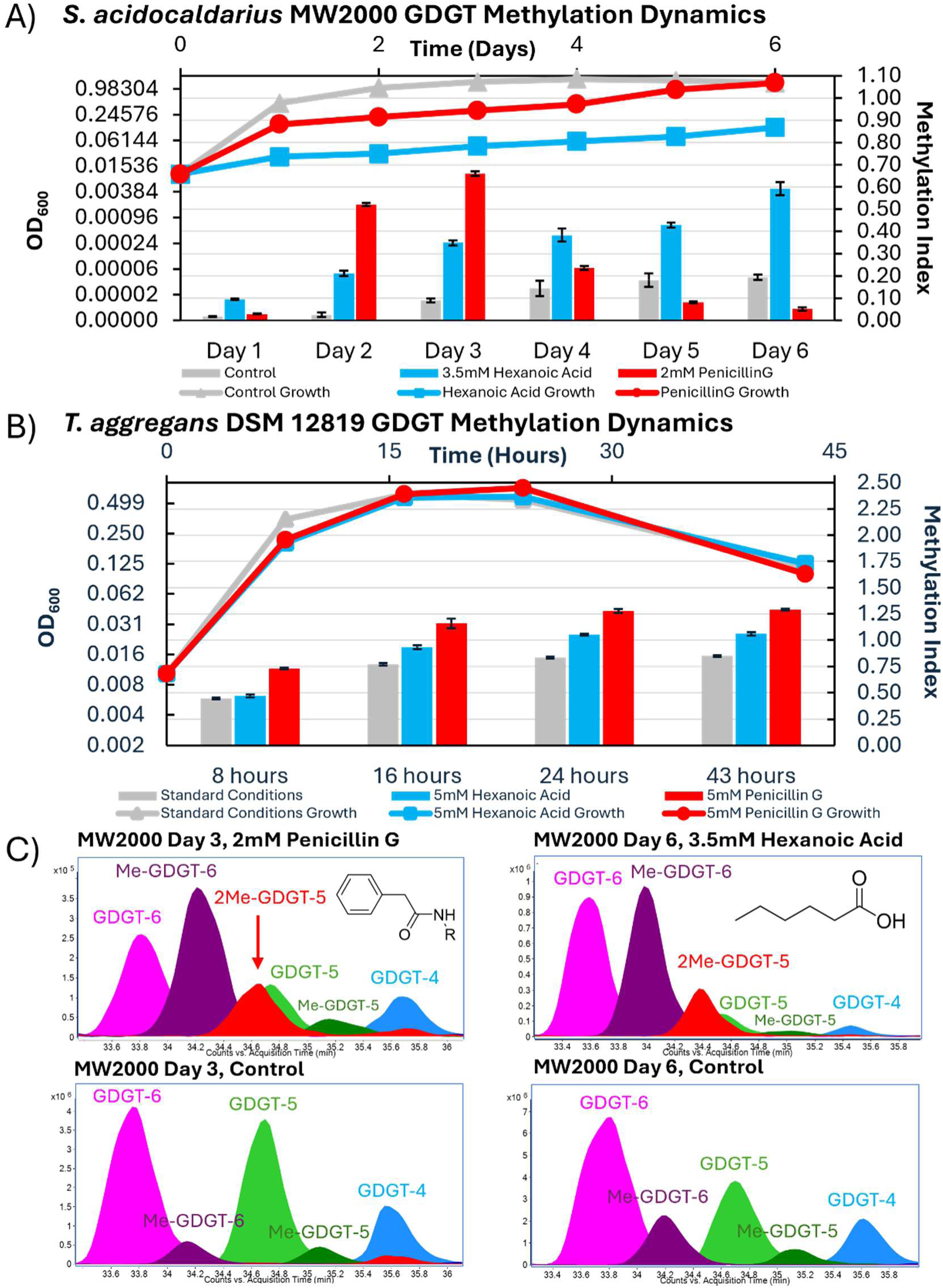
Exposure to the amphiphiles penicillin G and hexanoic acid increases GDGT lipid methylation in *S. acidocaldarius* and *T. aggregans*. A) Growth curves of *S. acidocaldarius* MW2000 under standard conditions (gray triangles), in the presence of 3.5 mM hexanoic acid (blue squares), and in the presence of 2.0 mM penicillin G (red circles) overlayed with the corresponding methylation index (MI_GDGT-4,5,6_), a measure of the weighted average number of extra methyl groups on GDGTs-4, 5, and 6, from core lipids of cells harvested the same day. Growth is impaired by both hexanoic acid and penicillin G, but *S. acidocaldarius* eventually recovers from penicillin G exposure during days 4-6. Similarly, GDGT methylation levels increase in response to both amphiphiles. GDGT methylation remains high in hexanoic acid treated cells but is only transiently upregulated in penicillin G exposed cells during days 2 and 3 and decreases when the growth recovers from days 4-6. B) Growth curve of *T. aggregans* under standard conditions (grey triangles) or in the presence of 5.0 mM hexanoic acid (blue squares) or 5.0 mM penicillin G (red circles) overlayed with the corresponding methylation index (MI_GDGT-0,1,2,3,4_). While growth is mostly unaffected by exposure to hexanoic acid and penicillin G at these levels, both compounds significantly increase GDGT methylation, albeit less so than in *S. acidocaldarius*. C) Representative, overlayed EIC of lipids extracted from acid-hydrolyzed biomass of control and amphiphile exposed *S. acidocaldarius* cultures on days 3 and 6, when methylation is highest in penicillin G and hexanoic acid treated cells, respectively.

Despite the difference in growth defects, we found that both archaea significantly increased Me-GDGT production in response to amphiphile exposure (Fig. 3A-C, S14). In the presence of hexanoic acid, *S. acidocaldarius* shows increased GDGT methylation levels across the 6-day time course, ranging from an MI_GDGT-4,5,6_ = 0.09 ± 0.00 at day 1 (vs. 0.02 ± 0.00 in the control) to an MI_GDGT-4,5,6_ = 0.59 ± 0.03 at day 6 (vs. 0.19 ± 0.01 in the control) (Fig. 3A,C). In contrast, penicillin G triggers a transient increase in GDGT methylation in *S. acidocaldarius.* We observed high levels of GDGT methylation on day 2 (MI_GDGT-4,5,6_ = 0.52 ± 0.01 vs. 0.03 ± 0.01 in the control) and day 3 (MI_GDGT-4,5,6_ = 0.66 ± 0.01 vs. 0.09 ± 0.01 in the control) which then decreases to near or below wildtype levels over the next 3 days (Fig. 3A,C). Also, the days of elevated GDGT methylation correspond to the most impaired growth period (days 2-4) and the subsequent decrease in methylation tracks with the recovery of *S. acidocaldarius* growth to near wild-type levels between days 4-6 (Fig. 3A). This recovery suggests that *S. acidocaldarius* can degrade penicillin G over the 6-day growth period and that the GDGT-methylation response is correspondingly down-regulated when this occurs.

In turn, the lack of recovery of *S. acidocaldarius* growth during exposure to 3.5mM hexanoic acid suggests that the archaeon is unable to completely degrade this compound over the 6-day time-period and suggests that the GDGT-methylation response remains “on”, resulting in the constitutively high Me-GDGT levels seen during the time course.

In *T. aggregans*, the increase in GDGT methylation in response to amphiphile exposure is apparent but less profound than in *S. acidocaldarius*, potentially due to the greater resistance of *T. aggregans* to these compounds or its higher basal levels of GDGT methylation. At equimolar concentrations, penicillin G triggers a greater level of GDGT methylation than hexanoic acid in *T. aggregans* (Fig. 3B). For penicillin G, the MI_GDGT-0,1,2,3,4_ (i.e. MI_GDGT_) is elevated across the time course, ranging from an exponential phase MI_GDGT_ = 0.73 ± 0.01 (vs. 0.45 ± 0.01 in the control) to a death phase MI_GDGT_ = 1.29 ± 0.01 (vs. 0.85 ± 0.01 in the control) (Fig. 3B, S14). In contrast, hexanoic acid exposure results in an elevated MI only during the stationary and death phases, with more modest increases ranging from an early stationary MI_GDGT_ = 0.93 ± 0.02 (vs. 0.77 ± 0.01 in the control) to a death phase MI_GDGT_ = 1.06 ± 0.02 (vs. 0.85 ± 0.01 in the control) (Fig. 3B). While hexanoic acid only induces moderate increases in GDGT methylation at 5mM levels, we hypothesized that higher concentrations may trigger greater Me-GDGT production. To test this, we attempted to grow *T. aggregans* in the presence of 10mM hexanoic acid; however, such levels were toxic, seemingly inhibiting growth. However, after 9 days a single replicate showed a small amount of growth. Analysis of the lipids of this culture revealed extremely high levels of GDGT methylation with an MI_GDGT_ = 2.32 (Fig. S14). We also observed low level production of penta-methylated GDGT-0 under these conditions, demonstrating that 5Me-GDGT-0 production is not simply an artefact during heterologous expression of Pgm in *T. kodakarensis*.

### Interplay of GDGT methylation and cyclization upon exposure to amphiphile stressors

Here, we demonstrated a clear and conserved increase in GDGT methylation in response to penicillin G and hexanoic acid exposure. However, the exact physiological role that this extra methylation may play remains unclear. Interestingly, we observe a concomitant increase in GDGT cyclization with the increase of GDGT methylation in *S. acidocaldarius* during exposure to the amphiphiles (Fig. S15). The highly-cyclized ring index (H-RI) or average number of cyclopentane rings amongst the dominant core lipids GDGT-4, 5, and 6 (essentially a measure of GrsB activity), increases from an H-RI = 4.28 ± 0.08 in the control on day 1 to an H-RI of 5.57 ± 0.02 and 5.05 ± 0.10 in hexanoic acid and penicillin G exposed cells, respectively. The H-RI continuously increases and remains high during hexanoic acid exposure, mirroring the trend of GDGT methylation under these conditions. Likewise, the H-RI increases up to day 3 in penicillin G exposed cells and then drops dramatically with GDGT methylation over the next two days. These data demonstrate that cyclization is also involved in the amphiphile-stress response of *S. acidocaldarius* and suggests that there may be an interplay amongst the two modifications.

Recent molecular dynamics simulations have indicated that the additional methyl group at C-13 in Me-GDGTs disrupts the packing of lipids and increases the fluidity of the membrane, at least when this methylation occurs on GDGT-0^28^. However, no specific investigation into the effect of this modification on cyclized GDGT structures has been carried out. If methylation generates a similar disruption in the packing of cyclized GDGTs and increases fluidity, it is curious why archaea would increase GDGT methylation in response to membrane destabilizing amphiphiles as these compounds also disrupt lipid packing and solubilize membranes, increasing their permeability and making them “leaky”. In contrast, the increase in cyclization in response to amphiphiles is consistent with observations that cyclopentane rings decrease fluidity and permeability and thus would counteract the effects of the amphiphiles. Perhaps the introduction of additional methyl groups on the highly cyclized GDGT-6 (the dominant membrane lipid in *S. acidocaldarius* amphiphile exposed cells) does moderately disrupt lipid packing to generate a membrane with properties in between that of an over-rigidified and impermeable GDGT-6 membrane and an over-fluid, leaky GDGT-4/5 membrane. In this way, the extra methylation(s) “fine-tune” membrane properties, allowing the cell to achieve a more optimal “goldilocks zone” of biophysical parameters than can be achieved by adjusting the ring number alone.

However, we did not observe any clear increases in GDGT cyclization in *T. aggregans* during penicillin G or hexanoic acid exposure, suggesting that the methylation itself could play a protective role against amphiphiles, perhaps sterically hindering the intercalation of these compounds into the membrane. Alternatively, GDGT methylation could have a more indirect role, such as being involved in a downstream signaling pathway as has been observed with cholesterol methylation and the regulation of dauer formation in nematodes^113^. Alternatively, the levels of amphiphiles used against *T. aggregans* may not have been high enough to induce the cyclization response. This is supported by observations that the growth of *T. aggregans* was not impaired by 5 mM hexanoic acid and 5 mM penicillin G and that when the growth was impaired by 10 mM hexanoic acid, we did observe a higher average number of rings per GDGT molecule with the RI = 1.51 (albeit in the singular sample that grew) as compared to the RI’s = 0.08 ± 0.01, 0.88 ± 0.06, 1.13 ± 0.01, and 1.07 ± 0.04 in the control during exponential, early stationary, late stationary, and death phase, respectively. Further, GDGT methylation and cyclization do concomitantly increase throughout the growth phases of *T. aggregans*, suggesting the two modifications may indeed be linked (Fig. S11&15). Additionally, Thermococci are more well known to respond to environmental stressors by altering the ratio of monolayer to bilayer lipids instead of employing cyclization (e.g. increasing the relative abundance of GDGTs to archaeol with increasing temperature to increase membrane rigidity)^6,114^. Thus, methylation could be used to “fine-tune” this strategy as well, increasing with higher levels of GDGTs to generate a membrane with properties intermediate that of a bilayer and monolayer. Altogether, these results reveal a novel and conserved connection between GDGT methylation and membrane-destabilizing amphiphiles amongst diverse archaea and call for a more in-depth characterization of the mechanistic role that Me-GDGTs play in response to these compounds.

### Identification of a tetraether desaturase from *Candidatus* Bathyarchaeota B1_G15

Having demonstrated that homologs of the previously characterized GMGT methylase (Gmm) can perform distinct and diverse lipid methylation reactions, we next sought to determine if more divergent archaeal Gmm homologs still possess methylase activity or if they might perform novel reactions. To do this we searched the NCBI database for lower homology Gmm homologs possessing e-values > 1e^-70^. Doing so, we uncovered a promising metagenomic B12-rSAM candidate (e-value = 1e^-60^, 31% identity to Gmm) from *Candidatus* Bathyarchaeota B1_G15. Involvement of this protein in archaeal lipid biosynthesis was suggested by its genomic co-localization with a GDGT ring synthase gene and a PPM webserver prediction that it is a peripheral membrane protein like Saci_1785 (ΔG_transfer_ = -10.5 ± 0.5 kcal/mol, PPM 3.0) (Fig. S16, Table S2). Further, the protein is modeled (Alphafold^115^ v.3) to possess a hydrophobic pocket directly beneath the membrane-associated portion of the protein, suggesting that it may bind overlying lipids (Fig. S16).

The Bathyarchaeon metagenome-assembled genome (MAG) in which this B12-rSAM enzyme gene was detected was assembled from deep-sea hydrothermal vent sediments of only moderate temperature (41°C)^116^. As such, we chose to characterize the activity of this B12-rSAM enzyme in the mesophilic archaeon *Methanosarcina acetivorans* C2A. However, we found that *M. acetivorans* only produces trace amounts of GDGTs (the presumed substrate) under standard growth conditions. Instead, the major membrane lipids are the bilayer lipid archaeol and its hydroxylated derivative, hydroxy-archaeol (OH-archaeol), that is formed by the phytoene desaturase family protein hydroxy-archaeol synthase (Has) via hydration of unsaturated intermediates (Fig. S17)^51^. Because we hypothesize that the Bathyarchaeon B12-rSAM would use GDGTs as a substrate, we first sought to increase GDGT production by overexpressing the native *tes* gene (*MA_1486*) in *M. acetivorans*. To do so, the gene was placed under the control of a tetracycline inducible promoter on the self-replicating plasmid pGLC157^117^ and we introduced this plasmid into *M. acetivorans*. Lipid analysis via LC-MS in reverse phase revealed the robust production of GDGT lipids, including GDGT-0 and its glycerol trialkyl glycerol tetraether (GTGT) intermediate (Fig. S17). We also detected the presence of abundant hydroxy-GDGTs (OH-GDGT-0 and 2OH-GDGT-0) in similar proportions to the OH-archaeols, suggesting Tes can covalently join OH-archaeols to produce OH-GDGTs (Fig. S17).

To investigate the activity of the Bathyarchaeon B12-rSAM enzyme, we co-expressed the protein with Tes in *M. acetivorans*. Strikingly, expression of this B12-rSAM enzyme resulted in the presence of a series of four new, earlier eluting GDGT peaks with a corresponding homologous series of OH-GDGTs and 2OH-GDGTs (and GTGTs) (Fig. 4A). The peak closest to GDGT-0 displayed an *m/z* value = 1300, two mass units less than that for GDGT-0 (*m/z* = 1302), and each subsequent peak differed by two more mass units (*m/z* = 1298, 1296, and 1294) suggesting each sequential compound differs by the presence of two hydrogen atoms. This loss of two hydrogen atoms can occur during the formation of either a double bond or a cyclization, suggesting the B12-rSAM enzyme is not a methylase. On the KTX column used to separate the lipids, ring-containing GDGTs typically elute after GDGT-0, in contrast to the earlier elution times seen for these new compounds^118^. Further, we previously demonstrated that double bond containing archaeols elute earlier than their saturated counterparts on this same column^35^. Together, this suggested that the newly observed peaks corresponded to a series of unsaturated GDGTs containing 1-4 double bonds.

**Figure 4.**
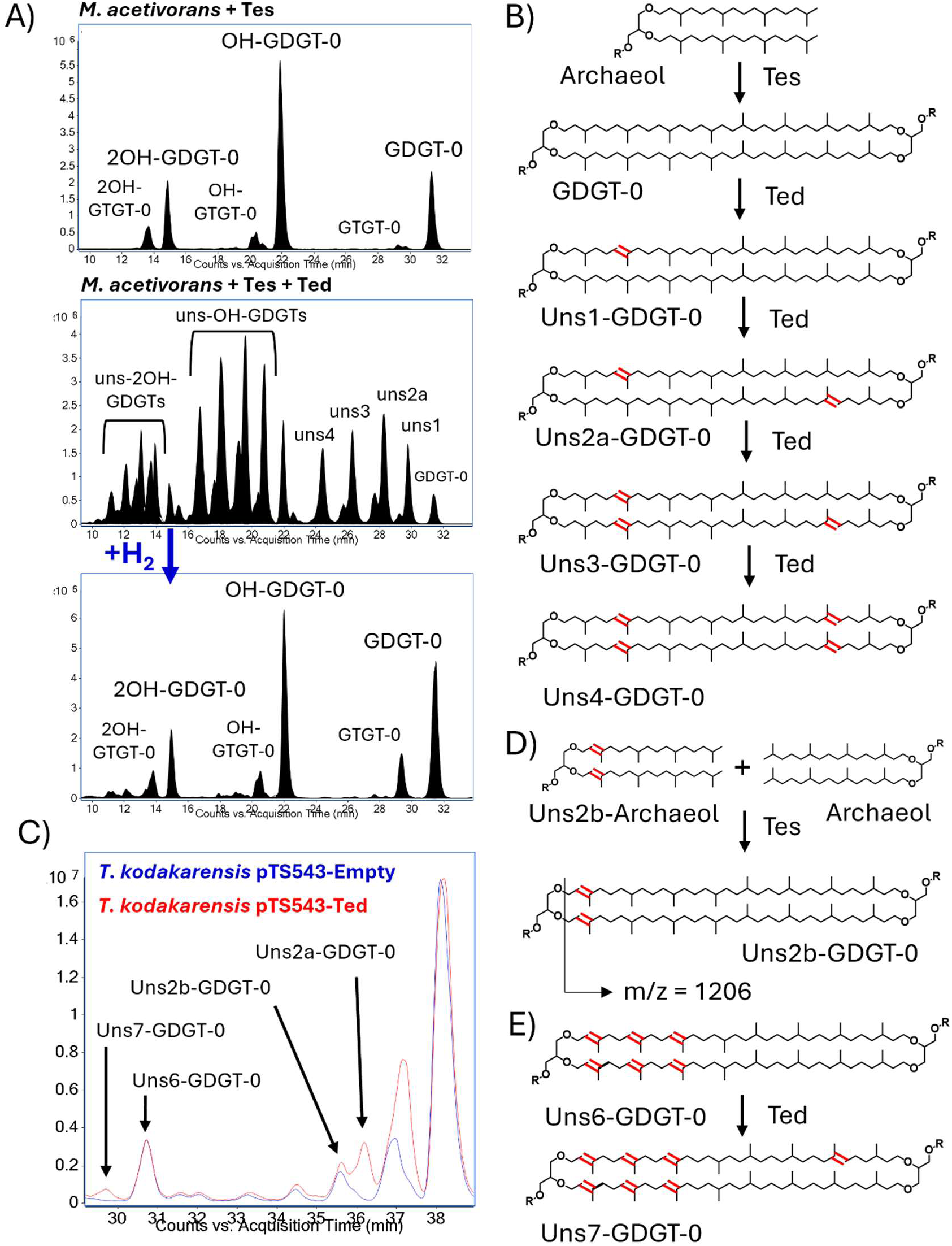
A B12-rSAM enzyme from *Candidatus* Bathyarchaeota B1_G15 encodes a tetraether desaturase (Ted). A) Representative, overlayed EICs of core lipids extracted from base-hydrolyzed biomass of *M. acetivorans* cultures expressing Tes alone or in combination with the B12-rSAM enzyme from *Candidatus* Bathyarchaeota B1_G15. Expression of Tes results in the production of three major GDGT lipids: GDGT-0, OH-GDGT-0, and 2OH-GDGT-0. Co-expression of Tes and the Bathyarchaeal B12-rSAM enzyme results in the production of a series of four new, earlier eluting GDGT peaks with a corresponding homologous series in both the mono- and di-hydroxy GDGTs. These new peaks differ in mass by 2 hydrogen atoms (e.g. *m/z* =1302 for GDGT-0 followed by 1300, 1298, 1296, and 1294 for the four previous peaks, respectively). Hydrogenation of these lipid extracts results in the loss of these new GDGT peaks, indicating that the unsaturations result from the formation of double bonds and not cyclization. B) The *Candidatus* Bathyarchaeota B1_G15 rSAM enzyme is a tetraether desaturase (Ted) which anaerobically forms 1-4 double bonds within the inert hydrocarbon tails of a GDGT. The double bonds are shown arbitrarily at C-7 as their exact positions have not yet been determined. C) Representative, overlayed EICs of core lipids extracted from base-hydrolyzed biomass of a *T. kodakarensis* control culture and a strain expressing a moderately thermophilic Ted homolog (96% identical to the *Candidatus* Bathyarchaeota B1_G15 Ted) grown at 50°C. At this low growth temperature for *T. kodakarensis*, the control strain natively produces a series of unsaturated GDGTs containing 1-6 double bonds which are thought to arise from the dimerization of unsaturated archaeol intermediates (which increase at lower growth temperature) with saturated archaeol. When expressing Ted, we see the production of new unsaturated GDGTs in *T. kodakarensis*: uns2a-GDGT-0 and uns7-GDGT-0 and an increase in uns1-GDGT-0 levels with a corresponding shift in the peak apex position. D) Putative biosynthetic route to unsaturated GDGTs in *T. kodakarensis* involving the dimerization of a saturated archaeol with a partially unsaturated archaeol biosynthetic intermediate. E) Putative biosynthetic route to the production of the uns7-GDGT-0 observed during Ted expression in *T. kodakarensis*.

To confirm this, we hydrogenated the lipids in an atmosphere of H_2_ gas in the presence of a palladium catalyst (Fig. 4A). Under these conditions, double bonds are fully saturated while rings are unreactive. Therefore, if unsaturation is lost upon hydrogenation, this indicates that the lipids possess double bonds and not rings. Hydrogenation of these lipid extracts resulted in the loss of the four earlier-eluting GDGT peaks, indicating that they are double-bond bearing derivatives of GDGTs: uns1-GDGT-0, uns2-GDGT-0, uns3-GDGT-0, and uns4-GDGT-0 (Fig. 4B). This demonstrates that the Bathyarchaeon B12-rSAM enzyme is a tetraether desaturase (Ted) capable of eliminating hydrogen atoms from archaeal lipids without the need for molecular oxygen, a major difference when compared to the oxygen-dependent fatty acid desaturases of bacteria and eukaryotes (Fig. 4A-B). It should be noted that we also detected low levels of hydrogenation-sensitive unsaturated archaeols in the *ted* expression strain (Fig. S18). These uns-archaeols differed from the native unsaturated archaeol intermediates found in the control strain, suggesting Ted may be able to also use bilayer lipids as a substrate but that it prefers GDGTs, similar to the substrate promiscuity of Pgm (Fig. S18).

The location of the GDGT double bonds introduced by Ted was not determined with certainty but was deduced to occur deep within the inert hydrocarbon tails. The double bonds cannot solely take place on the glycerol backbones as they can only accommodate one desaturation each (two total) whereas we observe up to four. Additionally, the presence of four double bonds in OH-GDGTs and 2OH-GDGTs suggests that the double bonds do not occur near the beginning of the tails (between C-2 and C-3 or C-3 and C-4) as the extra hydroxyl group of *Methanosarcina* lipids has been found to occur at the tertiary C-3 carbon^52^. The hydroxylated tertiary carbon thus has no hydrogen atoms to abstract to form a double bond, indicating Ted introduces double bonds elsewhere in the tails. The double bonds could either be between C-1 and C-2 or could be found deeper within the hydrocarbon tails. Introduction of a double bond between C-1 and C-2 would result in the formation of a vinyl ether. Vinyl bonds possess characteristic absorbance maxima around 190nm^119^, however, we did not detect any absorption at this wavelength for any of the unsaturated compounds. This suggests that Ted forms double bonds deep within the inert hydrocarbon tails of a GDGT. This is the first biological example, to our knowledge, of a radical SAM catalyzed desaturation at inert Csp^3^-Csp^3^ hydrocarbon centers.

To further validate this unusual desaturase activity, we sought to corroborate our results from *M. acetivorans* by expressing Ted in another archaeon, *T. kodakarensis*. However, *T. kodakarensis* is a hyperthermophile that grows optimally at 85°C whereas the Bathyarchaeon MAG possessing Ted is from 41°C sediments, suggesting it would likely not function at such high temperatures. As such, we searched the Joint Genome Institute (JGI) for Ted homologs from thermophilic metagenomes. We identified a closely related Ted homolog (96% identical) from a high temperature metagenome of Guaymas basin hydrothermal sediments. To express this gene in *T. kodakarensis*, we placed *ted* under the control of the strong promoter pcsg on plasmid pTS543 and introduced it into *T. kodakarensis* strain AL010.

No desaturase activity was observed when the Guaymas basin Ted homolog was overexpressed in *T. kodakarensis* at 85°C. We subsequently decreased the growth temperature to 50°C – the minimum growth temperature of *T. kodakarensis*. At this “low” growth temperature, we found that the control strain (carrying an empty pTS543 plasmid) produced moderate abundances of a series of putatively unsaturated GDGTs possessing 1-6 double bonds, with a distribution mirroring that of the unsaturation seen in the unsaturated-archaeol intermediates (1, 2, and 6 double bonds being most common) (Fig. S19). This suggests that the native unsaturated GDGTs seen in *T. kodakarensis* are formed by Tes linking a fully saturated archaeol molecule with a partially unsaturated archaeol (Fig. 4D, S19). This also implies that the uns-GDGTs in *T. kodakarensis* are asymmetric, possessing double bonds on one side of the molecule but not the other (Fig. 4D). In support of this, we observed temperature-dependent in-source fragmentation of uns-GDGTs possessing 2 or more double bonds, corresponding to the loss of the glycerol headgroup from the molecule (Fig. S20-21). Such cleavage can readily occur if the uns-GDGTs contained two allylic double bonds (the naturally occurring C2-C3 double bonds found in archaeal lipid biosynthesis intermediates) on the same side of the GDGT molecule but on opposite tails (Fig. S19-21). Such a configuration would occur by the dimerization of saturated and unsaturated archaeol by Tes. Furthermore, this fragmentation pattern is not observed in the unsaturated GDGTs produced by heterologous expression of Ted in *M. acetivorans*. This demonstrates that the double bond introduced by Ted is not located at the C3 position and suggests that we can distinguish the unsaturated GDGTs produced by Ted from those natively occurring in *T. kodakarensis* grown at 50°C.

The heterologous expression of the Guaymas Basin Ted homolog in *T. kodakarensis* at 50°C resulted in the formation of new uns-GDGTs that did not display in-source fragmentation (Fig. 4C, S20). In particular, the uns-GDGTs produced by Ted elute slightly earlier than the native uns-GDGTs in *T. kodakarensis* and are chromatographically aligned with the unsaturated GDGTs produced by the *Candidatus* Bathyarchaeon Ted in *M. acetivorans*. This is most clearly seen for the Ted produced uns2a-GDGT-0 which is chromatographically separable from the uns2b-GDGT-0 natively seen in *T. kodakarensis* (Fig. 4B-D, S20). Uns1-GDGT-0 produced by Ted also appears to elute earlier but is not completely separable from the native uns1-GDGT-0 in *T. kodakarensis*; however, the peak apex of uns1-GDGT-0 is clearly shifted earlier and is more abundant than in the empty plasmid control (Fig. 4C). Furthermore, the expression of Ted in *T. kodakarensis* increased the maximum number of double bonds observed in GDGTs from 6 to 7, suggesting Ted desaturates the natively produced uns6-GDGT-0, yielding uns7-GDGT-0 (Fig 4C,E). This consistent desaturation of GDGTs by Ted homologs in *M. acetivorans* and *T. kodakarensis* supports the classification of Ted as the first known lipid desaturase in archaea.

### Identification of a hydroxy-GDGT synthase belonging to the AttH-like hydratase family

Inspection of the genomic neighborhood of *ted* in *Candidatus* Bathyarchaeota B1_G15 and the Guaymas basin metagenome scaffold revealed that the coding sequence of *ted* overlaps with a conserved gene encoding an AttH-like protein (Fig. 5A). Notably, members of this protein family include 1,2-carotenoid hydratases (CrtC)^120^ and kievitone hydratases^121^, both of which hydrate an isoprene double bond to form a tertiary alcohol. This suggested that the *attH* gene overlapping with *ted* might encode an AttH-like protein that similarly hydrates the double bond(s) of uns-GDGTs formed by Ted to produce hydroxy-GDGTs (OH-GDGTs). To test this, we heterologously expressed the gene encoding the AttH-like protein from the Guaymas basin scaffold alone and in combination with Ted in *T. kodakarensis,* which does not natively produce OH-GDGTs. LC-MS analysis of lipid extracts from the empty plasmid control strain did not detect any OH-GDGTs (Fig. 5B). In contrast, expression of the AttH-like protein alone resulted in the production of OH-GDGT-0 in two isomeric forms but only in trace amounts, suggesting that the preferred substrate of the AttH-like protein is not being produced (Fig. 5B-C). In support of this, co-expression of the AttH-like protein with Ted resulted in the robust formation of both OH-GDGT-0 and 2OH-GDGT-0, demonstrating that this protein is a hydroxy-GDGT synthase (Hgs) that appears to preferentially hydrate the double bonds formed by Ted (Fig. 5B-D, S22). Additionally, we observed the formation OH-uns6-GDGT-0, corroborating that the uns7-GDGT-0 observed in the Ted expression strain is produced by Ted via desaturation of uns6-GDGT-0 and that it can then be hydrated at the introduced double bond to form OH-uns6-GDGT-0 (Fig. S23). The action of a protein working on the downstream uns-GDGT products of Ted further validates the desaturase activity of the Ted enzyme. Additionally, AttH-like hydratases are only known to introduce hydroxyl groups at tertiary carbons, suggesting that the double bond introduced by Ted must also be located at a tertiary carbon atom, either at C-7, C-11, or C-15.

**Figure 5.**
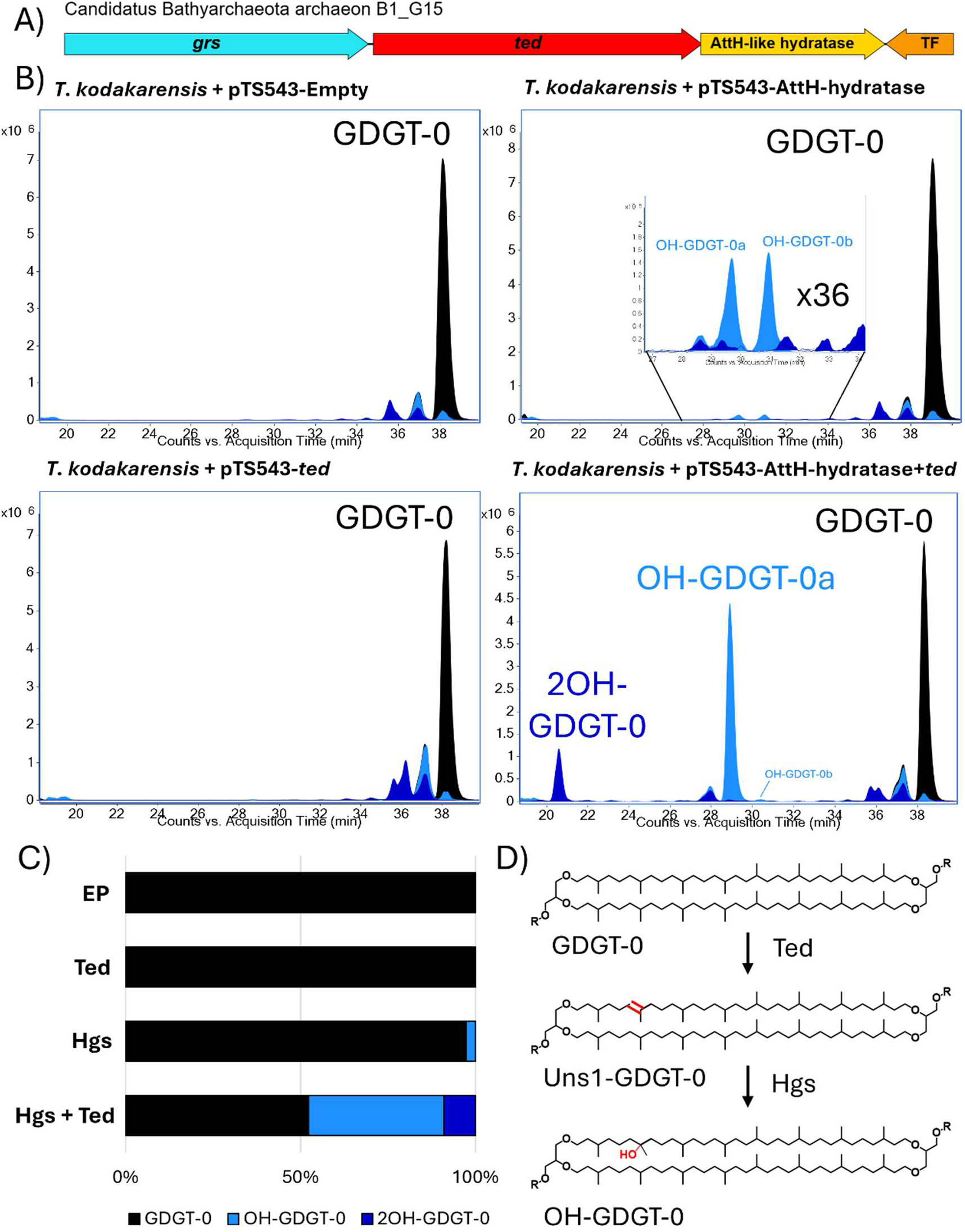
An AttH-like hydratase co-occurs with Ted and encodes a hydroxy-GDGT synthase (Hgs). A) Illustration of the genomic neighborhood of *ted* in *Candidatus* Bathyarchaeota B1_G15 showing that the desaturase gene co-occurs with *grs* and overlaps with an AttH-like hydratase encoding gene. B) Representative, overlayed EICs of core lipids extracted from *T. kodakarensis* carrying an empty pTS543 plasmid (control stain), *T. kodakarensis* expressing Ted alone, and *T. kodakarensis* expressing the AttH-like hydratase alone and in combination with Ted. The empty plasmid control strain produces no hydroxylated lipids. Expression of Ted alone does not lead to the production of hydroxylated lipids. Expression of the AttH-like hydratase alone results in the trace production of OH-GDGTs in two isomeric forms, suggesting that it can form OH-GDGTs but that the native uns-GDGTs in *T. kodakarensis* are not its preferred substrate. Expression of Hgs and Ted together results in the robust production of mono- and di-hydroxy GDGTs, suggesting that Hgs preferentially hydrates the double bonds introduced by Ted to form OH-GDGTs. C) Average relative abundance of core GDGT-0, OH-GDGT-0, and 2OH-GDGT-0 in the four *T. kodakarensis* strains analyzed for OH-GDGT production. Only the strain co-expressing Hgs (the AttH-hydratase) and Ted produces abundant OH-GDGTs. D) The AttH-like hydratase, whose coding sequence overlaps with the *ted* gene, is a hydroxy-GDGT synthase (Hgs) that putatively hydrates the double bonds introduced by Ted to form OH-GDGTs.

BLASTP searches of the NCBI database for Ted (e-value = 1e^-100^, ≥ 500 amino acids) and Hgs (e-value = 1e^-80^, ≥ 300 amino acids), revealed that Ted homologs are distributed amongst Bathyarchaeota (30%), Methanobacteriota such as ANME archaea (20%), Thermoplasmatota such as *Methanomassiliicoccus luminyensis* (19%), and Promethearchaeati/Asgardarchaeota (10%). However, Hgs homologs are more phylogenetically restricted. We identified Hgs homologs almost exclusively in Methanobacteriota (53%) including ANME archaea and the methyl-reducing methanogen *Candidatus* Methanofastidiosum methylothiophilum, and in Bathyarchaeota (25%). This suggests that while Ted is sometimes employed to form uns-GDGTs for hydroxy-GDGT biosynthesis, that the unsaturated GDGTs are not exclusively intermediates to other biosynthetic products. Rather, unsaturated GDGTs may have important functional roles themselves, especially in archaea belonging to Thermoplasmatota and Promethearchaeati/Asgardarchaeota that lack Hgs including the cultured archaeon with a complete genome *M. luminyensis* B10.

Our identification and characterization of a tetraether desaturase and a hydroxy GDGT synthase demonstrates a biosynthetic link between OH-GDGTs and uns-GDGTs in certain archaeal clades. An alternative route to OH-GDGT-0 biosynthesis likely exists in methanogens such as *M. acetivorans* whereby Tes links a “normal” archaeol with an OH-archaeol produced by the phytoene desaturase family protein, hydroxy-archaeol synthase (Has)^51^, that hydrates the unsaturated intermediates in archaeol biosynthesis. This route to OH-GDGTs (and OH-archaeols) requires the synthesis of new lipids as Has works on intermediates in the biosynthetic pathway. In contrast, archaea may employ Ted and Hgs to convert existing GDGT lipids into OH-GDGTs under conditions where new lipid production is unfavorable (e.g. low nutrient conditions), allowing the archaea to respond to environmental challenges quickly even under low nutrient regimes.

## Conclusion

The chemical diversity of lipids produced by life is large, and archaeal membrane lipids are no exception. How archaea generate a diverse lipid pool was largely unknown until the discovery of radical SAM (rSAM) enzymes – tetraether synthase (Tes)^7,8^ and GDGT ring synthase (Grs)^10^ – that were shown to be responsible for the formation and cyclization of archaeal monolayer lipids known as GDGTs, respectively. Since then, rSAM enzymology has been found to underlie a number of archaeal lipid modifications including the covalent crosslinking of GDGT tails to form GMGTs and their subsequent methylation to form Me-GMGTs^11^. In this study, we further highlight the importance of rSAM enzymes in generating diverse archaeal lipids, demonstrating that they are responsible for forming multiple types of Me-GDGTs and for desaturating GDGTs to form unsaturated GDGTs. While bacteria and eukaryotes utilize di-iron, monooxygenase family enzymes to perform lipid desaturation reactions^25^, it is notable that archaea have evolved the tetraether desaturase (Ted) to perform an alternative, oxygen-independent desaturation reaction utilizing organic radical chemistry. Ted is a B12-binding rSAM related to the GDGT methylases (Cgm and Pgm) characterized in this study, highlighting the importance of exploring and characterizing homologous rSAM enzymes in archaea as they may possess diverse activities. Further, we demonstrate the utility of investigating the genomic context of archaeal lipid biosynthesis genes, as this can provide clues to the physiological roles of certain lipid modifications (e.g. the involvement of GDGT methylation in amphiphile stress responses) and reveal other non-rSAM enzymes that are involved in GDGT biosynthesis such as the hydroxy-GDGT synthase (Hgs). Altogether, this study advances our understanding of archaeal lipid biosynthesis and physiology and highlights the nuances that have evolved in lipid biochemistry and resulted in the diversity of membrane structures observed today across all domains of life.

## Methods

### Microbial strains, growth conditions, and media recipes

All strains used for this study are listed in the supplementary information, Table S3. *Sulfolobus acidocaldarius* MW2000 and its derivatives were grown at 75°C and pH = 3.0 in Brock medium^122^ supplemented with 0.1% NZ-amine and 0.2% dextrin at 75°C, unless otherwise stated. The parental and deletion strains are uracil auxotrophs and were supplemented with 10 µg/mL uracil. For growth and lipid analysis experiments, 50 mL liquid cultures were inoculated with actively growing culture to an OD_600_ = 0.01. Overexpression and complementation strains were induced with 0.4% dextrin and harvested on day 3 in late-exponential phase. For amphiphile exposure experiments, powdered Penicillin G or pure liquid hexanoic acid were added to the media after autoclaving, immediately prior to inoculating the cultures. For growth on plates, media was solidified with 0.6% gelrite, 3mM CaCl_2_, and 10mM MgCl_2_.

*Thermococcus aggregans* DSM12819^123^ and *Thermococcus kodakarensis* AL010^35^ and its derivatives were routinely cultured in artificial sea water (ASW) medium prepared in a Coy anaerobic chamber and supplemented with 5 g/L yeast extract, 5 g/L tryptone, 2 g/L sulfur, Wolfe’s trace elements, and KOD vitamins (also known as ASW-YT)^124^. The parental *T. kodakarensis* strain AL010 is an agmatine auxotroph and was supplemented with 1 mM agmatine sulfate. For growth and lipid analysis experiments, 40 mL cultures were inoculated with an actively growing culture to an OD_600_ = 0.01 and were grown under fermenting conditions: ASW-YT lacking sulfur and additionally containing 5 g/L dextrin and 10 µM sodium tungstate dihydrate or 5 g/L sodium pyruvate (basal conditions for pfba-inducible expression). *T. aggregans* was grown at 85°C and *T. kodakarensis* was grown at 85°C, except for Ted and Hgs expression experiments which were carried out at 50°C. For expression experiments at 85°C, cells were harvested after 12-16 hours in late exponential phase. For expression experiments at 50°C, cells were harvested after 7 days of growth. For amphiphile exposure experiments, powdered Penicillin G or pure liquid hexanoic acid were added to the media after autoclaving, inside the anaerobic chamber and immediately prior to inoculating the cultures. For growth on plates, ASW-YT media was solidified with 1% gelrite and supplemented with 2 mL/L of polysulfide solution (10 g Na_2_Sx9H_2_O combined with 3 g of elemental sulfur in 15 mL of DI water) instead of sulfur powder.

*Methanosarcina acetivorans* C2A derivatives were all grown in bicarbonate-buffered high-salt media supplemented with 50mM trimethylamine (HS-TMA) as their growth substrate and 80%:20% N_2_:CO2 gas headspace composition as previously described^125^. For routine cultivation strains were grown in 10ml of HS-TMA in 26ml Balch-type tubes sealed with butyl rubber stoppers. Growth of M. acetivorans cultures was monitored by absorbance at 600 nm in a UV-Vis spectrophotometer (Genesys 50, Thermo Fisher Scientific, Waltham, MA, USA). For lipid analysis *M. acetivorans* cultures were grown in 80ml of HS-TMA media in 150ml serum vials. Tetracycline induction of lipid modifying enzymes was achieved by the addition of an appropriate volume of 10mg/ml anaerobic stock solution (100x) to yield a final concentration of 100µg/ml. Cells were harvested in late exponential phase by centrifugation at 10,000 g for 10 minutes at 4°C (Sorvall Legend XTR,472 Thermo Fisher Scientific, Waltham, MA, USA).

All *Escherichia coli* strains were grown at 37 °C in lysogeny broth (LB) and supplemented with 100 µg/mL carbenicillin, 30 µg/mL kanamycin and/or 20µg/ml Chloramphenicol.

### Molecular cloning and archaeal strain construction

Primers and plasmids used in this study are listed in Tables S4 and S5, respectively. *S. acidocaldarius* plasmids were constructed using overlap-extension PCR and restriction/ligation cloning. To construct *saci_1785* (*cgm*), *saci_1585* (*grsA*), and *saci_0240* (*grsB*) deletion plasmids, we first PCR amplified approximately 700 bp of the upstream (U) and downstream (D) region of each gene using primers 1785F1/R1 and 1785F2/R2 (for *saci_1785*), 1585F1/R1 and 1585 F2/R2 (for *saci_1585*), and 0240F1/R1 and 0240F2/R2 (for *saci_0240*). Next, the U and D region of each gene were fused by overlap-extension PCR, and the joined product was then amplified by addition of the appropriate F1 and R2 primer pair. Finally, the products were inserted into plasmid psva407^126^ at the ApaI and BamHI cut sites to yield plasmids psva407-*saci_1785*UD, psva407-*saci_1585*UD, and psva407-*saci_0240*UD. To construct the *saci_1785* expression plasmid, *saci_1785* was amplified using primers Fwd_BsphHI_S1785 and Rv_S1785_NcoI and inserted into plasmid psva1561 at the NcoI cut site, yielding plasmid psva1561-*saci_1785*. The *S. acidocaldarius* markerless deletion mutants and expression strains were constructed as previously described^127^. Briefly, cells were transformed with methylated plasmid DNA from *E. coli* ER1821 via electroporation, and transformants were selected by growth on plates lacking uracil and with blue-white LacS-based screening. To confirm expression strains, single-colony PCR targeting the psva1561 insert was performed with primers Fwd_psva1561_checking and Rv_psva1561_checking and verified with long-read sequencing at Plasmidsaurus (www.plasmidsaurus.com). For the second step of markerless deletion mutant construction, 5-FOA containing plates were used for second selection, followed by single-colony PCR with primers Fwd/Rv_1785_checking, Fwd/Rv_1585_checking, or Fwd/Rv_0240_checking and verified with long-read sequencing. To obtain and transform the Δ*grsA* deletion mutant, cells were plated at lower temperature (65°C) due to temperature sensitivity of growth.

*T. kodakarensis* pTS543 expression plasmids were synthesized from Twist Biosciences. All genes were codon optimized for expression in *T. kodakarensis* with a codon adaptation index value between 0.7-0.8. In particular, *pgm* from *T. aggregans* TY/DSM12819 (locus tag: *NF865_RS03100*), *ted* from the Guaymas basin metagenome scaffold (gene ID: *GBSed_1000338310*, scaffold: GBSed_10003383, biosample ID: Gb0103921), and *hgs* from the same scaffold (gene ID: *GBSed_1000338311*) were synthesized with either the pfba^86,128^ promoter (for *pgm*) or pcsg^129^ promoter (for *ted* and *hgs*) immediately upstream of the start codon and cloned into pTS543 at the NotI site to yield plasmids pTS543-pfba-*pgm*, pTS543-pcsg-*ted*, and pTS543-pcsg-*hgs*. For co-expression of *pgm* and the native *T. kodakarensis grsA* (TK0167), grsA was synthesized with the pcsg promoter immediately upstream of the start codon and cloned into pTS543 at the BamHI site while *pgm* was introduced as described above, yielding the plasmid pTS543-pcsg-*grsA*-pfba-*pgm*. For co-expression of *ted* and *hgs*, the genes were synthesized together with one pcsg promoter immediately upstream of the start codon of *hgs* which was connected to *ted* via a ribosome binding site (rbs) as follows: tg<u>aGGTGA</u>TATGAatg (rbs underlined, stop and start codons of *hgs* and *ted*, respectively, lower case), yielding plasmid pTS543-pcsg-*hgs*-rbs-*ted*.

Construction of *T. kodakarensis* expression strains was carried out as described previously^11,124^. Briefly, cells were grown to early stationary phase (10-12 hours) and then transformed with approximately 3 µg of plasmid DNA. Transformation was carried out by incubating cells in the presence for 30 minutes on ice, heat shocking for 45-60 seconds at 85°C, and returning to ice for another 30 minutes before plating. Plates were sealed and incubated in a metal anaerobic canister that was kindly given to us by the Santangelo lab at Colorado State University. Transformants were selected by growth on plates lacking agmatine and were confirmed by PCR amplifying the insert region of pTS543 using primers Fwd/Rv_pTS543_checking and verified with long-read sequencing.

*M. acetivorans* expression plasmids were constructed based on the tetracycline induction system developed by Guss et al^117^. Briefly, the strong tetracycline-inducible promotor *PmcrB(tetO1)* drives the expression of Beta-glucuronidase (UidA) in the plasmid pJK027A. To test our various lipid modification enzymes, UidA was removed from pJK027A by digestion with HindIII and NdeI and replaced with M. acetivorans’ native *tes* (MA1486) as well as the *ted* gene from *Candidatus* Bathyarchaeota B1_G15 (locus tag: *DRO69_02625* synthesized from Twist Bioscience). Genes were inserted using Gibson assembly^130^ (see Supplementary Table for primers). The UidA-containing pJK027A served as a negative control plasmid for lipid modification experiments. For the maintenance of these plasmids in *M. acetivorans* pJK027A and derivatives are cointegrated with pAMG40, which contains the pC2A replication machinery, by fosmid retrofitting using Gateway™ BP Clonase™ II (Thermo Fisher). *M. acetivorans* mutants were constructed using liposome-mediated transformation as previously described^131^, and isolated on 1.5% agar plates containing HS-TMA media and 2 µg/mL puromycin as the selective agent. The parental strain for mutant construction was WWM60 (Δ*hpt*::P*mcrB*-*tetR*) which carries the TetR repressor required for tetracycline induction.

### Lipid extraction and analyses

Frozen cell pellets were thawed and resuspended in 2 mL of methanol (MeOH) which was then transferred to 4 mL glass vials and evaporated under N_2_ stream. Samples were then acid or base hydrolyzed by the addition of 2 mL of 1M HCl in MeOH or 1M KOH in MeOH, respectively. Acid hydrolyzed samples were incubated at 85°C for 16 hours, and base hydrolyzed samples were incubated at 75°C for 3 hours. Once the incubation was complete, the reactions were neutralized by the addition of 1 mL of 2M KOH in MeOH or 2M HCl in MeOH for acid and base hydrolyzed samples, respectively. Samples were then transferred to 40 mL glass vials, diluted with 5 mL of deionized water, and extracted three times with 5 mL dichloromethane (DCM). Pooled DCM extracts were evaporated under N_2_ stream. Samples were resuspended in 0.5-1 mL of 9:1 MeOH:DCM or 0.5-1 mL 1:1 isopropanol:hexane for LC-MS analysis on a Kinetex 1.7 µm XB-C18 100 Å LC column or an ACE UltraCore Super C18 column, respectively, and filtered through 0.45 µm polytetrafluoroethylene filters into glass vials.

Samples were analyzed on an Agilent 1260 Infinity II series high-performance liquid chromatography machine connected to an Agilent G6125B single quadrupole mass spectrometer with electrospray ionization in positive mode. The following parameters were kept constant during all analyses: drying gas flow rate (8.0 L/min), nebulizer pressure (35 psi), and capillary voltage (3500 V). The drying gas temperature and fragmentor voltage were adjusted based on the type of lipids being analyzed. For samples containing only inert lipids (e.g. saturated GDGTs, Me-GDGTs, and archaeols) a drying gas temperature of 300°C and a fragmentor voltage of 175 V were used. For more samples containing more labile lipids (e.g. unsaturated GDGTs and OH-GDGTs) a lower drying gas temperature of 225°C and fragmentor voltage of 0V were used to minimize the decomposition of the lipid species. Samples were analyzed in scanning mode between a range of m/z = 600 – 1500.

Separation of lipids was achieved on either 1) an Agilent Poroshell 120 EC-C18 column (1.8 µm, 2.1 × 150 mm) connected in series with an ACE UltraCore Super C18 column (5 μm, 2.1 × 250 mm) with a mobile phase A of 95% MeOH and 5% water with 0.04% formic acid and 0.03% NH_3_ additives and mobile phase B of 50% isopropanol and 50% hexane with 0.04% formic acid and 0.03% NH_3_ additives based on previous methods^132,133^ or 2) a Kinetex 1.7 µm XB-C18 100 Å LC column (150 × 2.1 mm) with a mobile phase A of 100% MeOH with 0.04% formic acid and 0.03% NH_3_ additives and a mobile phase B of 100% isopropanol with 0.04% formic acid and 0.03% NH_3_ additives based on a method from Rattray and Smittenberg^118^. Lipids were identified based upon elution times/order and m/z values in combination with characteristic in-source fragmentation (for certain uns-GDGTs) and dehydration (for OH-GDGTs).

LC-MS data was analyzed using Agilent MassHunter (B.08.00). Unless otherwise stated, the relative abundances of tetraether lipids were calculated based on the peak areas of the protonated and sodiated adducts of the parent ion, [M+H]^+^ and [M+Na]^+^, respectively. For example, the extracted ion chromatograms in Fig.1 were generated by extracting the following ions: m/z = (1294.3, 1316.3; GDGT-4), (1308.3, 1330.3; Me-GDGT-4), (1322.3, 1344.3; 2Me-GDGT-4), (1292.3, 1314.3; GDGT-5), (1306.3, 1328.3; Me-GDGT-5), (1320.3, 1342.3; 2Me-GDGT-5), (1290.3, 1312.3; GDGT-6), (1304.3, 1326.3; Me-GDGT-6), and (1318.3, 1340.3; 2Me-GDGT-6). When a lower drying gas temperature was used for analysis of unsaturated-GDGTs and hydroxylated GDGTs, ammoniated [M+NH_4_]^+^ and di-ammoniated [M+2NH_4_]^2+^ adducts were taken into account as well.

### Bioinformatic and phylogenetic analyses

The *S. acidocaldarius* Cgm (Saci_1785), *Thermococcus guaymasensis* Gmm (WP_062369928.1), *S. acidocaldarius* GrsA (Saci_1585), *T. kodakarensis* GrsA’ (TK0167), *S. acidocaldarius* GrsB (Saci_0240), and *Caldivirga maquilingensis* GrsB’ (Cmaq_1853) were used as queries for a BLASTP search to retrieve sequences for the construction of a phylogenetic tree of GDGT methylases and other B12-binding radical SAM enzymes involved in GDGT biosynthesis. Protein sequences were aligned with MAFFT^134^ on XSEDE (7.505; BLOSUM62 scoring matrix; gap penalty of 1.53). The resultant alignment was used to build a phylogenetic tree using IQtree^135^ (2.2.0) which was visualized with the Interactive Tree of Life (iTOL)^136^.

For analysis of the genomic neighborhood of GDGT methylases, we first generated a sequence similarity network (SSN) of Cgm/Pgm proteins using the Enzyme Function Institute Enzyme Similarity Tool (EFI-ESI). Saci_1785 was used as the query at an e-value of 1e-99. The resultant SSN was then input into the EFI Genome Neighborhood Tool (EFI-GNT) to generate a genome neighborhood network (GNN) of the Cgm-related proteins. The neighborhood region analyzed was set to 10 genes upstream and downstream of *cgm* and a minimal co-occurrence value of 25% was selected as the threshold. The resultant GNN can be accessed here: https://efi.igb.illinois.edu//efi-gnt/view_diagrams.php?gnn-id=38820&key=a11a457a21f580a4aebf1205e32098913938557a (cluster 1).

## Acknowledgments

We would like to thank the Santangelo lab at Colorado State University for the generous gift of the *Thermococcus kodakarensis* AL010 parental strain and cuboid. We would also like to thank the Albers lab at the University of Freiburg for the generous gift of the *Sulfolobus acidocaldarius* MW2000 strain. This work was supported by NSF Grant EAR-1752564 (to P.V.W). Portions of the experiments were performed in the Stanford Geomicrobiology Shared Laboratories Core Facility (RRID:SCR_025000). DDN and GLC would like to acknowledge funding from the Simons Foundation through the Simons Early Career Investigator in Marine Microbial Ecology and Evolution Award.

