## Supplemental Files for "Identification of radical SAM enzymes responsible for the methylation and desaturation of archaeal lipids and an AttH hydratase mediating hydroxy-GDGT biosynthesis"

### Supplementary Information

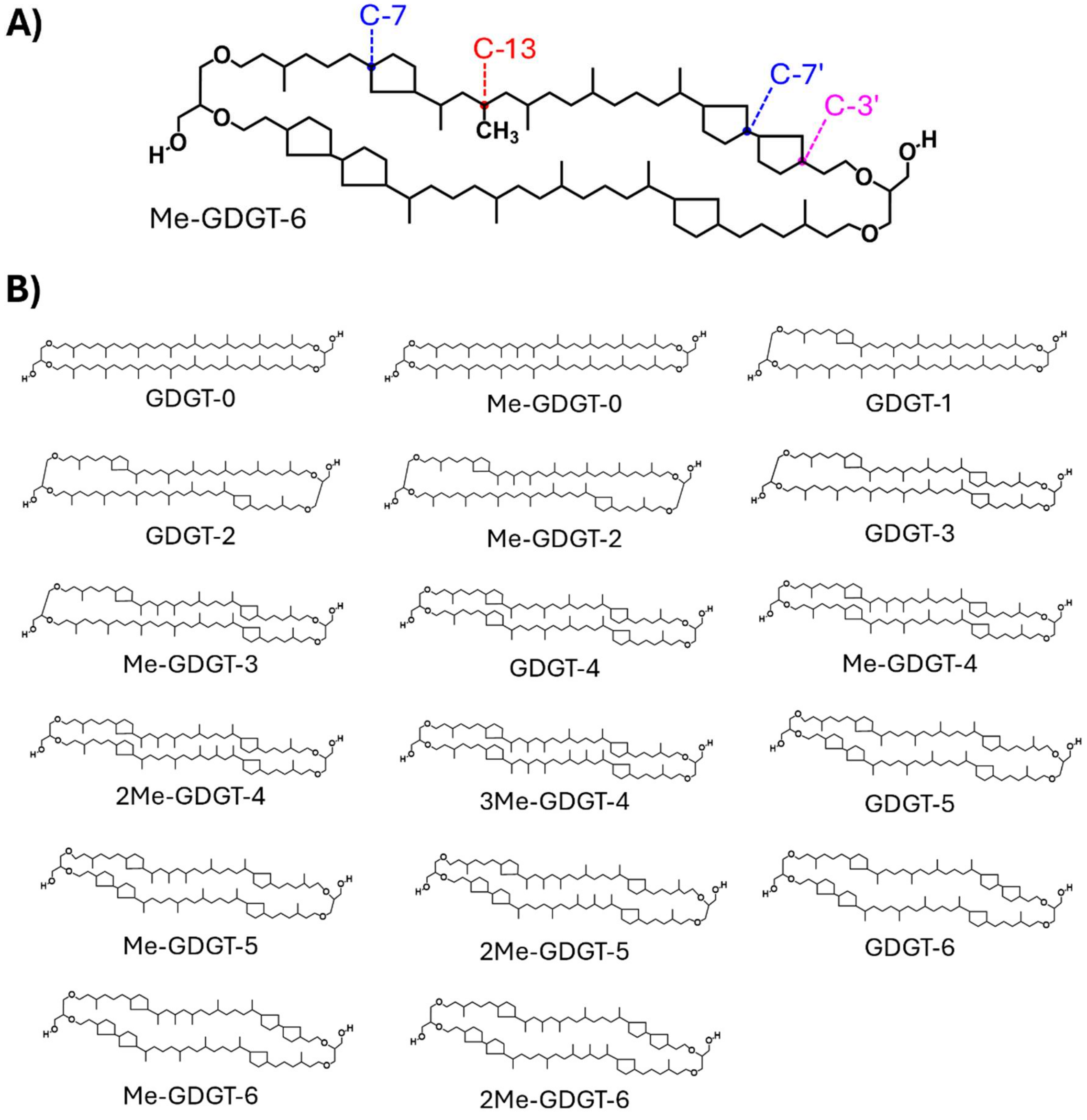


**Fig. S1 *S. acidocaldarius* core lipid structures discussed in text.** A) Tentative structure of Me-GDGT-6. The extra methyl group is located at C-13 (red) based on previous studies[1]. Other sites of GDGT modification are C-7 (blue; cyclization by GrsA) and C-3 (pink; cyclization by GrsB). B) Core lipid structures found in the various *S. acidocaldarius* strains used in this work. One isomer is shown for each lipid species, but multiple isomers of each lipid species are possible due to variation in the orientation of the glycerol backbone (parallel vs. antiparallel), variation in ring location (C3 vs. C7 or on the same vs. opposite tail), and variation in methylation location (same vs. opposite tail).

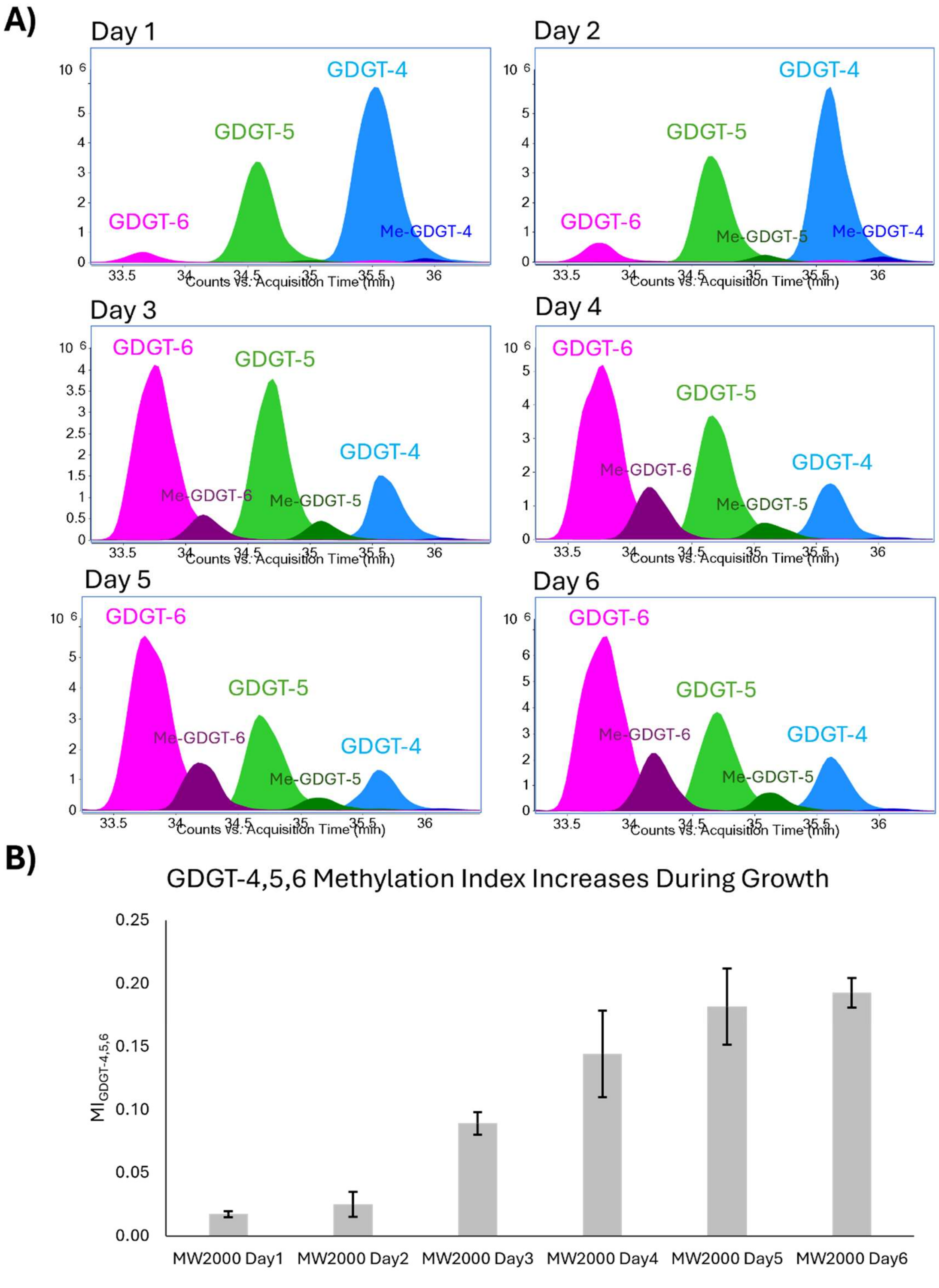


**Fig. S2 GDGT methylation increases throughout the growth phases of *S. acidocaldarius* MW2000.** A) Representative overlayed, extracted ion chromatograms (EICs; *m/z* values below*) of lipids extracted from acid hydrolyzed biomass of *S. acidocaldarius* strain

MW2000 across a 6-day time-course under standard growth conditions. Methylation is lowest on day 1 and increases gradually to day 6. B) Methylation index of GDGTs-4, 5, and 6 ($MI_{GDGT-4,5,6}$; the average number of extra methyl groups per molecule of GDGTs-4, 5, and 6) across 6 days of *S. acidocaldarius* growth under standard conditions.

**m/z* = (1294.3, 1316.3; GDGT-4), (1308.3, 1330.3; Me-GDGT-4), (1292.3, 1314.3; GDGT-5), (1306.3; 1328.3; Me-GDGT-5), (1290.3, 1312.3; GDGT-6), (1304.3, 1326.3; Me-GDGT-6)

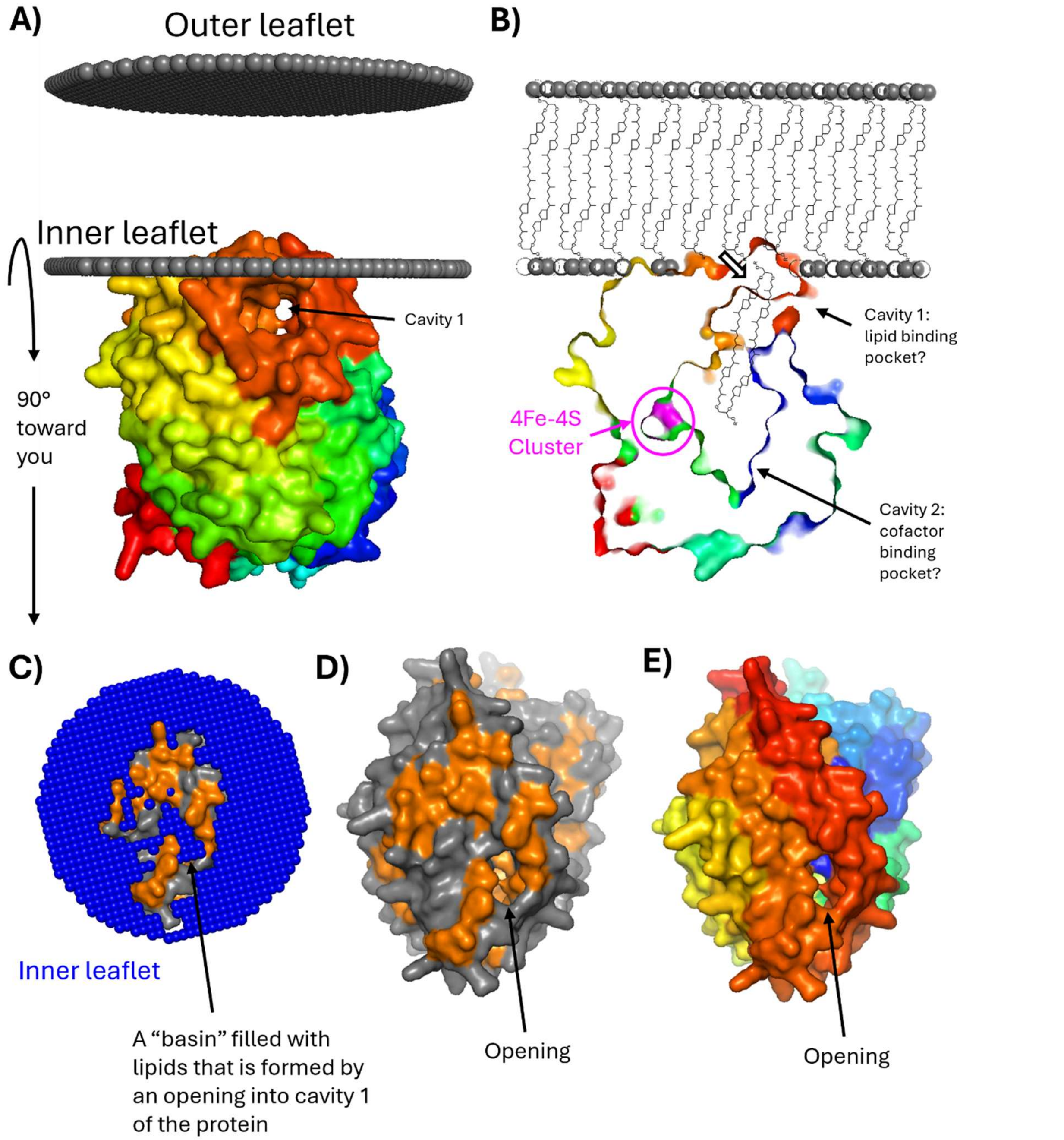


**Fig. S3 Saci_1785 is predicted to be a peripheral membrane protein.** A) The Alphafold[2] model (v.3) of Saci_1785 is positioned peripherally in the cell membrane by the Positioning of Proteins in Membranes (PPM 3.0)[3,4] webserver. The protein is colored blue at the N-terminus and proceeds through the rainbow to red at the c-terminus which is partially embedded in the inner leaflet of the cell membrane. A cavity/binding pocket is modeled directly beneath the membrane-associated portion of the protein. B) A central cross-section of the *saci_1785* encoded protein. The cross section depicts two connected cavities. Cavity 1 is modeled directly beneath the membrane-associated portion of the protein and could perhaps be a lipid binding pocket that retrieves its substrate (e.g. GDGT-6) from the overlying cell membrane. Cavity 2 is modeled to possess the CXXXCXXC motif which binds the 4Fe-4S cluster of radical SAM proteins; thus, cavity 2 may be the cofactor binding pocket and its connection with cavity 1 may facilitate interaction of the substrate with the active site. C) Alphafold model from “A” rotated 90° forward (toward the reader) with the outer leaflet removed to reveal the embedded portion of the protein in the inner leaflet. Hydrophobic residues are shown in orange, and lipid ends/heads are shown in blue. The protein is modeled to possess an opening into cavity one which forms a “basin” that can be filled by the overlying lipids. D) Same as “C” but with the outer leaflet removed to just show the protein structure. Note the opening into cavity 1 that may facilitate uptake of a lipid substrate from the membrane. E) Same as “D” but colored as in “A”.

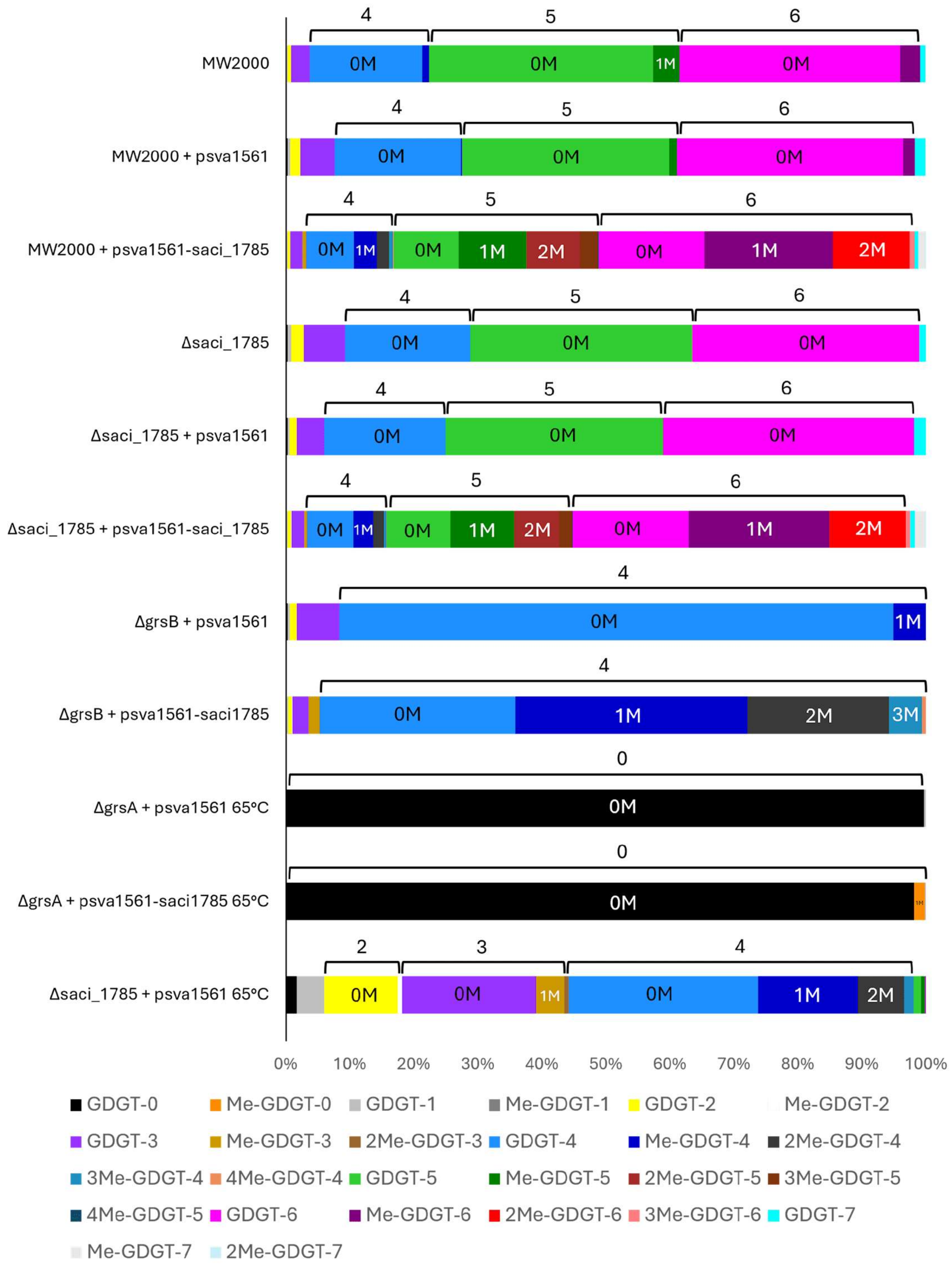


**Fig. S4 Average relative abundance of different core lipid species in the various *S. acidocaldarius* strains used in this study.** The parental strain MW2000 primarily makes the highly cyclized GDGTs-4, 5, and 6 which are predominantly in their unmethylated form

(0M). The parental strain carrying an empty plasmid (MW2000 + psva1561) has a similar lipid profile to MW2000. The parental strain overexpressing *saci_1785* (MW2000 + psva1561-*saci_1785*) converts approximately half of GDGT-4 and a majority of GDGTs-5 and 6 into their mono- (1M) and di-methylated (2M) derivatives (Me-GDGT-4, Me-GDGT-5, and Me-GDGT-6). The *saci_1785* deletion strain (Δ*saci_1785*) lacks all Me-GDGTs as does the *saci_1785* deletion strain carrying an empty plasmid (*Δsaci_1785* + psva1561). Me-GDGT production is restored in the *saci_1785* deletion strain complemented with *saci_1785* (Δ*saci_1785* + psva1561) which produces much higher abundances of Me-GDGTs than MW2000. The *grsB* deletion strain carrying an empty plasmid (Δ*grsB* + psva1561) primarily makes GDGT-4 which is predominantly in its unmethylated form. The *grsB* deletion strain overexpressing *saci_1785* (Δ*grsB* + *saci_1785*) converts a majority of GDGT-4 into its mono-, di-, and tri-methylated (3M) derivatives. The *grsA* deletion strain carrying an empty plasmid (Δ*grsA* + psva1561) and grown at 65°C (rather than the optimal 75°C due to temperature sensitivity) primarily makes GDGT-0 with no detectable methylated derivatives. The *grsA* deletion strain overexpressing saci_1785 (Δ*grsA* + *saci_1785*) and grown at 65°C primarily makes GDGT-0 but now produces a minor amount (~2% of core lipids) of its mono-methylated derivative Me-GDGT-0. The *saci_1785* deletion strain complemented with *saci_1785* (Δ*saci_1785* + psva1561-*saci_1785*) and grown at the same temperature (65°C) primarily makes GDGTs-2, 3, and 4 at this lower temperature; it converts approximately half of GDGT-4 into its mono-, di-, and tri-methylated derivatives (much more methylation than is seen on GDGT-0 when overexpressing *saci_1785* in Δ*grsA*) and converts a smaller percentage of GDGT-3 (~20%) into its mono- and di-methylated derivatives and converts an even smaller percentage (~5%) of GDGT-2 into its mono-methylated derivative.

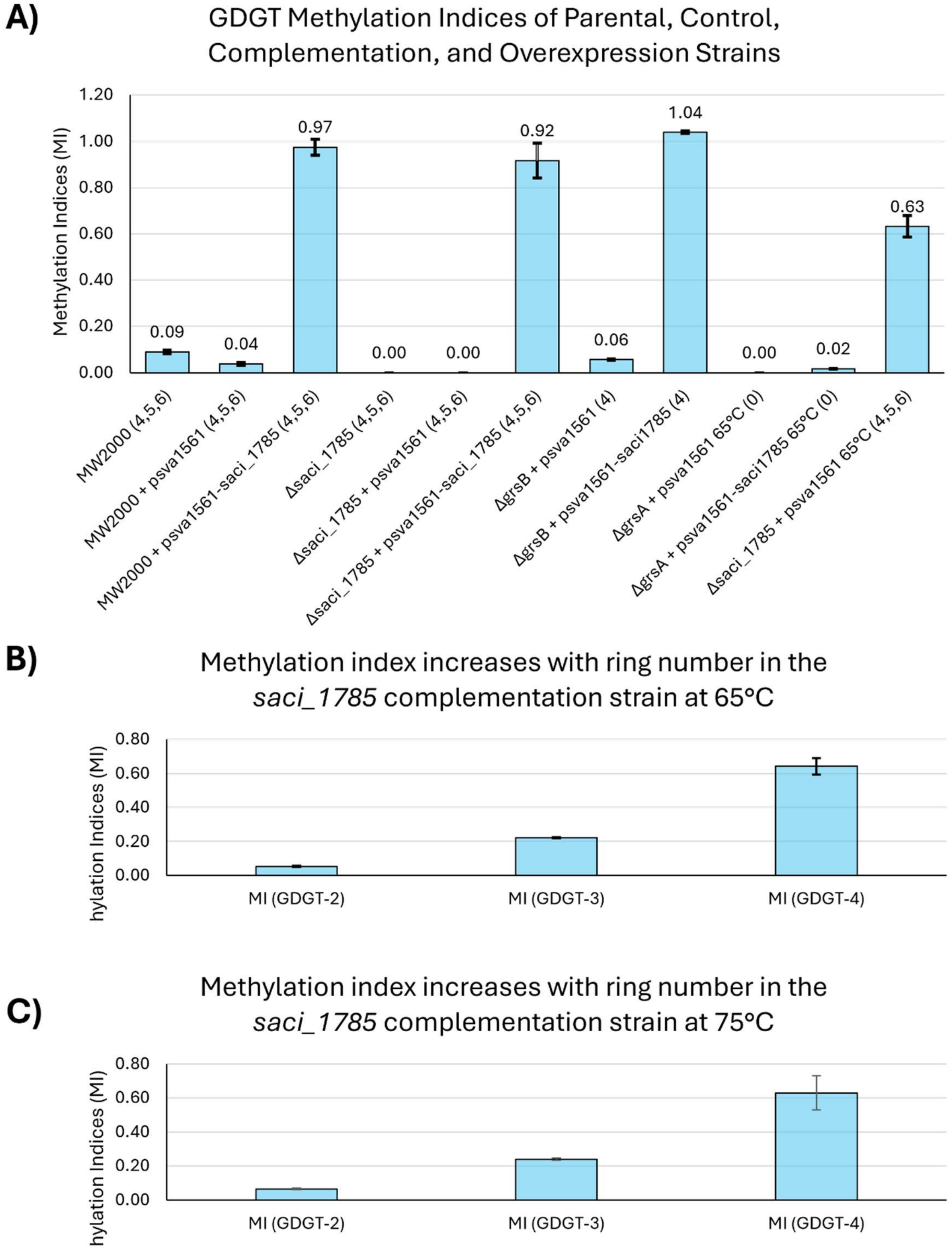


**Fig. S5 GDGT methylation increases during *saci_1785* overexpression and with increasing ring number.** A) Comparison of the GDGT methylation indices of the different *S.*

*acidocaldarius* strains used in this study. The MI is highest in the *saci_1785* complementation and overexpression strains, except for Δ*grsA* + psva1561-*saci_1785* which only methylates the major core lipid GDGT-0 at trace levels. B) Methylation indices of individual GDGTs during expression of *saci_1785* in the complementation strain at 65°C, when the relative abundance of GDGTs-2 and 3 is comparable to GDGT-4. The methylation of GDGTs by Cgm gradually increases with ring number, with no methylation detected on GDGT-0 and GDGT-1, minor methylation on GDGT-2, moderate methylation on GDGT-3, and robust methylation on GDGT-4. C) Methylation indices of the individual GDGTs during expression of *saci_1785* in the same strain but at 75°C, when the relative abundance of GDGTs-2 and 3 is much lower than GDGT-4. Here, the methylation of GDGTs by Cgm increases with ring number, consistent with the results at 65°C.

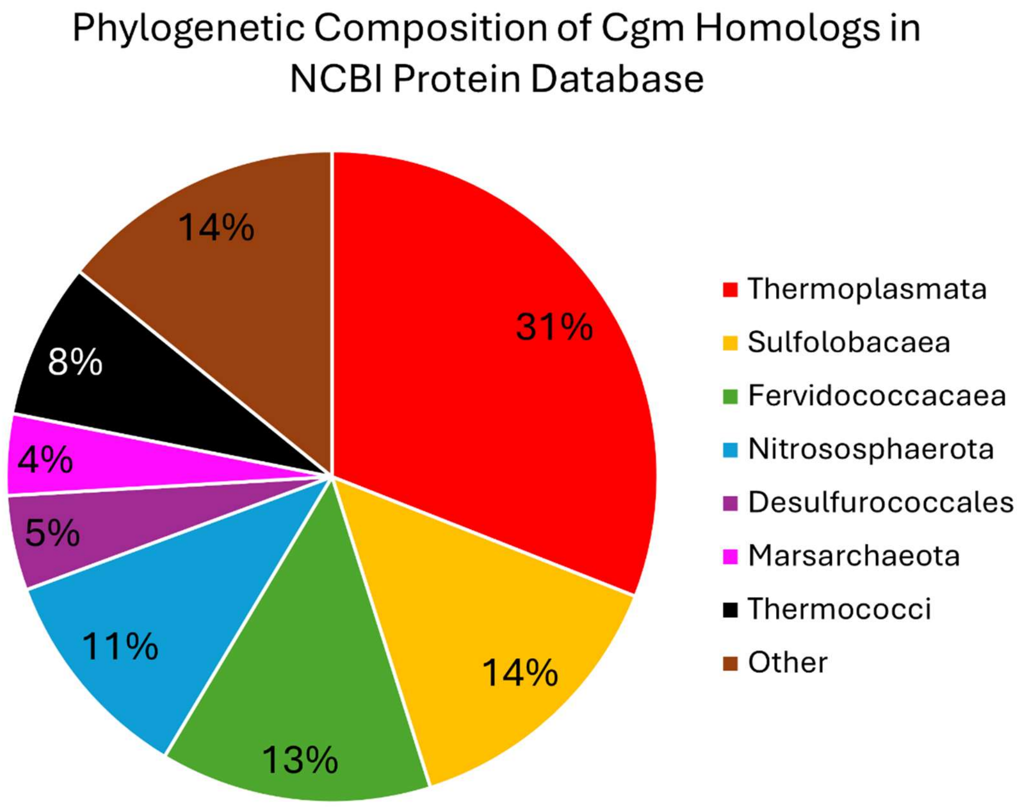


**Fig. S6 Phylogenetic composition of Cgm homologs found in the NCBI protein database.** Cgm homologs are restricted to the TACK and Euryarchaeota and are primarily found in the (thermo)acidophilic Thermoplasmata, Sulfolobacaea, Nitrososphaerota, and Marsarchaeota. They are also found in the more neutrophilic (neutral pH optima) Fervidococcacaea and Thermococci.

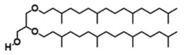
Archaeol

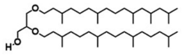
Me-Archaeol

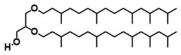
2Me-Archaeol

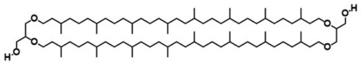
GDGT-0

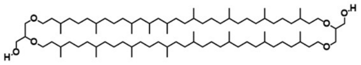
Me-GDGT-0

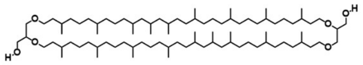
2Me-GDGT-0

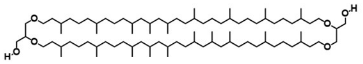
3Me-GDGT-0

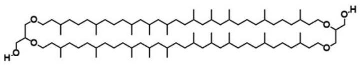
4Me-GDGT-0

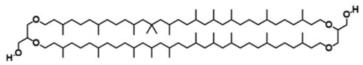
5Me-GDGT-0

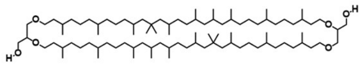
6Me-GDGT-0

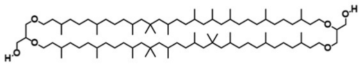
7Me-GDGT-0

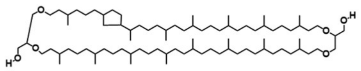
GDGT-1

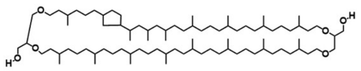
Me-GDGT-1

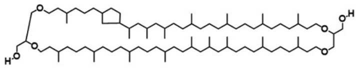
2Me-GDGT-1

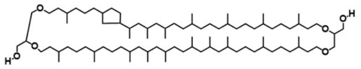
3Me-GDGT-1

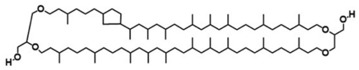
4Me-GDGT-1

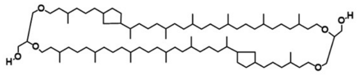
GDGT-2

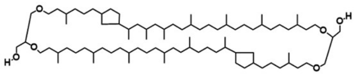
Me-GDGT-2

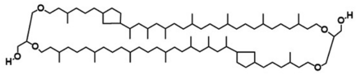
2Me-GDGT-2

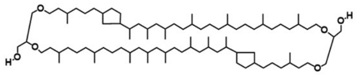
3Me-GDGT-2

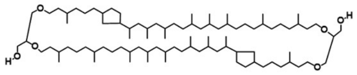
4Me-GDGT-2

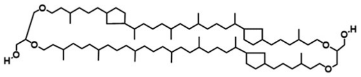
GDGT-3

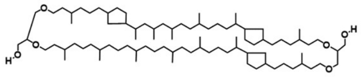
Me-GDGT-3

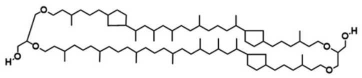
2Me-GDGT-3

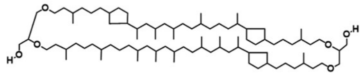
3Me-GDGT-3

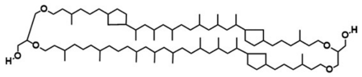
4Me-GDGT-3

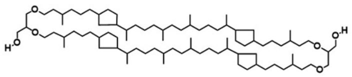
GDGT-4

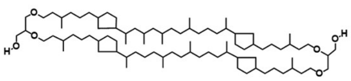
Me-GDGT-4

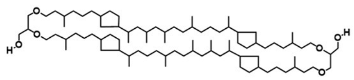
2Me-GDGT-4

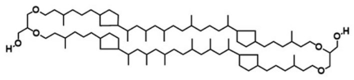
3Me-GDGT-4

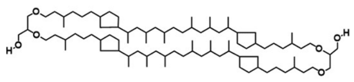
4Me-GDGT-4

**Fig. S7 Core lipid structures detected in the *Thermococcus* strains used in this study.** One potential isomer of each lipid species is shown but multiple regioisomers are possible for each.

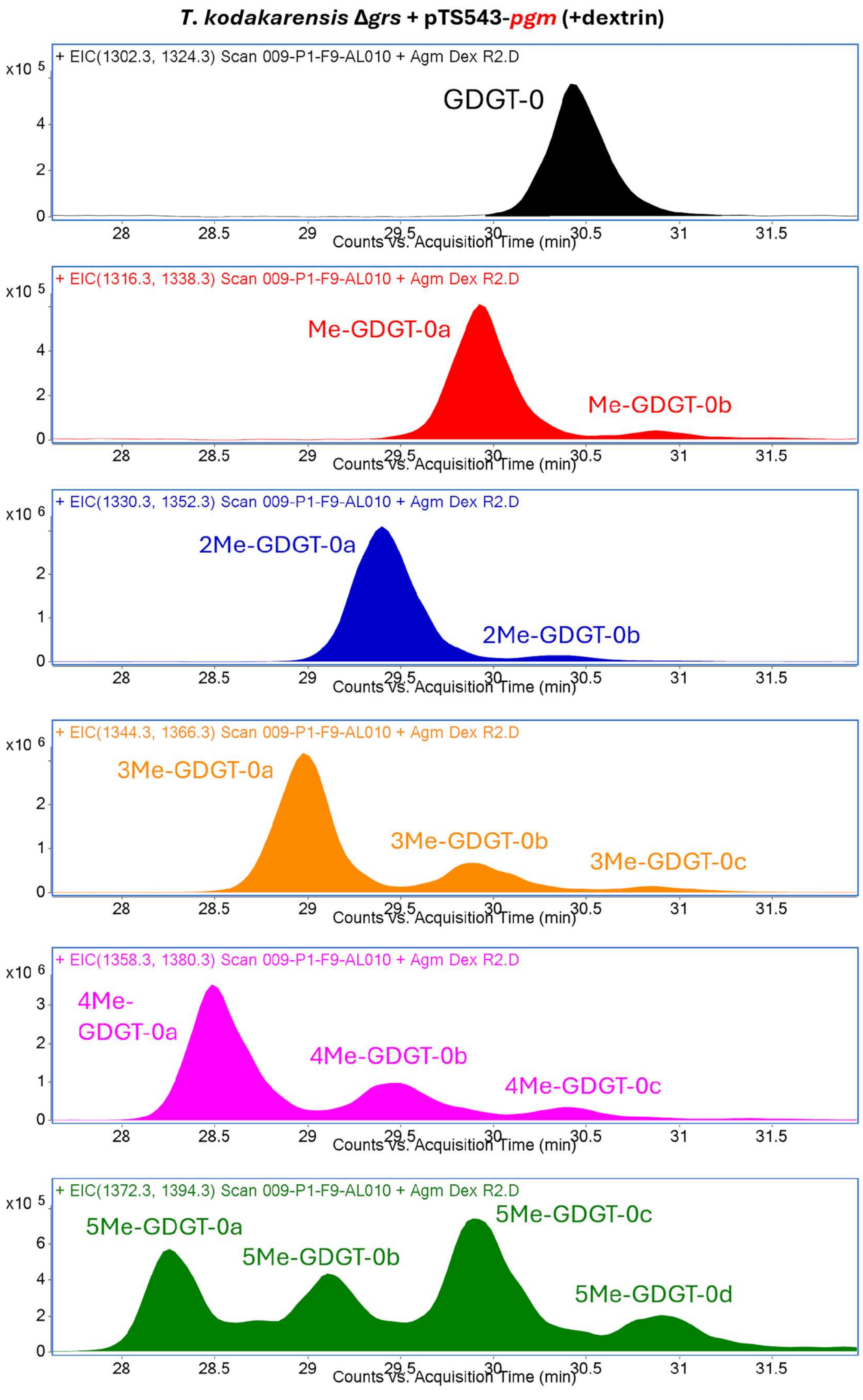
T. kodakarensis Δgrs + pTS543-pgm (+dextrin)
+ EIC(1302.3, 1324.3) Scan 009-P1-F9-AL010 + Agm Dex R2.D
x10 5
GDGT-0
Counts vs. Acquisition Time (min)
+ EIC(1316.3, 1338.3) Scan 009-P1-F9-AL010 + Agm Dex R2.D
x10 5
Me-GDGT-0a
Me-GDGT-0b
Counts vs. Acquisition Time (min)
+ EIC(1330.3, 1352.3) Scan 009-P1-F9-AL010 + Agm Dex R2.D
x10 6
2Me-GDGT-0a
2Me-GDGT-0b
Counts vs. Acquisition Time (min)
+ EIC(1344.3, 1366.3) Scan 009-P1-F9-AL010 + Agm Dex R2.D
x10 6
3Me-GDGT-0a
3Me-GDGT-0b
3Me-GDGT-0c
Counts vs. Acquisition Time (min)
+ EIC(1358.3, 1380.3) Scan 009-P1-F9-AL010 + Agm Dex R2.D
x10 6
4Me-GDGT-0a
4Me-GDGT-0b
4Me-GDGT-0c
Counts vs. Acquisition Time (min)
+ EIC(1372.3, 1394.3) Scan 009-P1-F9-AL010 + Agm Dex R2.D
x10 5
5Me-GDGT-0a
5Me-GDGT-0b
5Me-GDGT-0c
5Me-GDGT-0d
Counts vs. Acquisition Time (min)

**Fig. S8 Pgm produces multiple Me-GDGT isomers.** Me-GDGTs detected in representative extracted ion chromatograms (*m/z* values displayed in upper left-hand corners) of acid hydrolyzed lipid extracts from the induced *pgm* expression strain (*Δgrs* + pTS543-*pgm* + dextrin). Each Me-GDGT is observed to possess at least two chromatographically resolvable isomers. While multiple isomers are present, a single isomer is the major product for each Me-GDGT species except for 5Me-GDGT-0 which has multiple isomers of similar abundance. For mono-, di-, tri-, and tetra-methylated GDGT-0, the major/most abundant isomer is also the earliest eluting isomer.

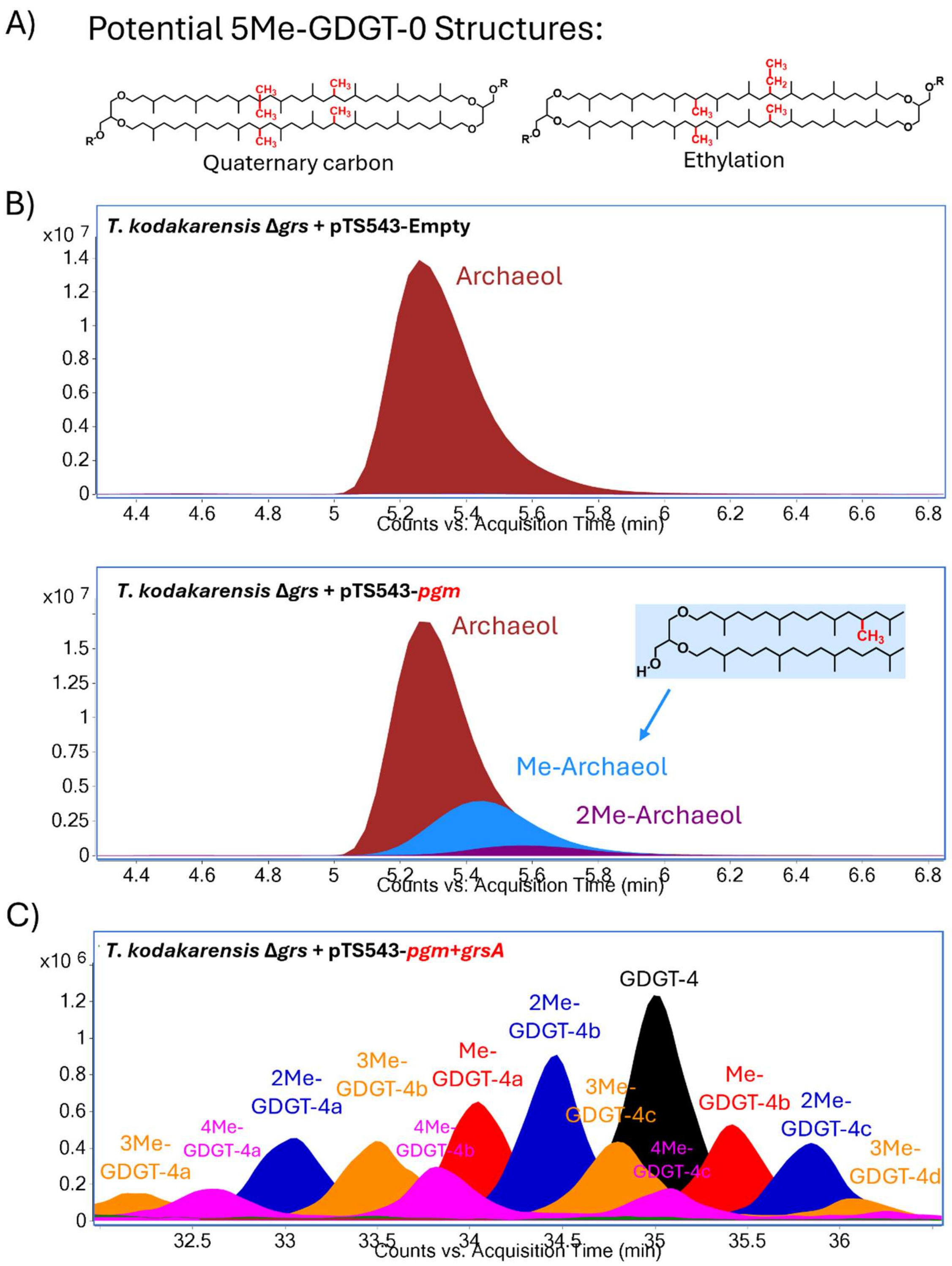


**Fig. S9 Penta-methylated GDGTs, methylated archaeols, and methylated cyclized GDGTs are produced during Pgm expression in *T. kodakarensis*.** A) Diether lipids detected in representative, overlayed, extracted ion chromatograms (EICs; *m/z* values below*) of acid hydrolyzed core lipid extracts from the empty plasmid control strain (*Δgrs* + pTS543-empty) and the *pgm* expression strain (Δ*grs* + pTS543-*pgm*). No methylated derivatives of archaeol are observed in the control strain; in contrast, a minor amount of archaeol is methylated in the *pgm* expression strain to produce mono- and di-methylated archaeol (Me-Archaeol and 2Me-Archaeol, respectively). B) GDGT-4 lipid species detected in a representative, overlayed EIC (*m/z* values below**) of acid hydrolyzed core lipid extracts from the *pgm* + *grsA* co-expression strain (Δ*grs* + pTS543-*pgm*+*grsA)*. A high level of methylation is observed on the GDGT-4 lipids, similar to the level of methylation seen on GDGT-0. The methylated-derivatives of GDGT-4 are present in multiple, chromatographically resolvable isomeric forms of similar abundance. In contrast, methylated GDGT-0 lipids have a single dominant isomer that is much more abundant than the other isomers. Me-GDGT-4 possesses two such isomers, 2Me-GDGT-4 possesses three, and 3Me- and 4Me-GDGTs each possess four. The structural differences between the isomers are currently unknown.

**m/z* = (653.7, 670.8, 675.7; Archaeol), (667.7, 684.8, 689.7; Me-Archaeol), and (681.7, 698.8, 703.7; 2Me-Archaeol)

***m/z* = (1294.3, 1316.3; GDGT-4), (1308.3, 1330.3, Me-GDGT-4), (1322.3, 1344.3; 2Me-GDGT-4), (1336.3, 1358.3; 3Me-GDGT-4), (1350.4, 1372.3; 4Me-GDGT-4), (1364.4, 1386.3; 5Me-GDGT-4)

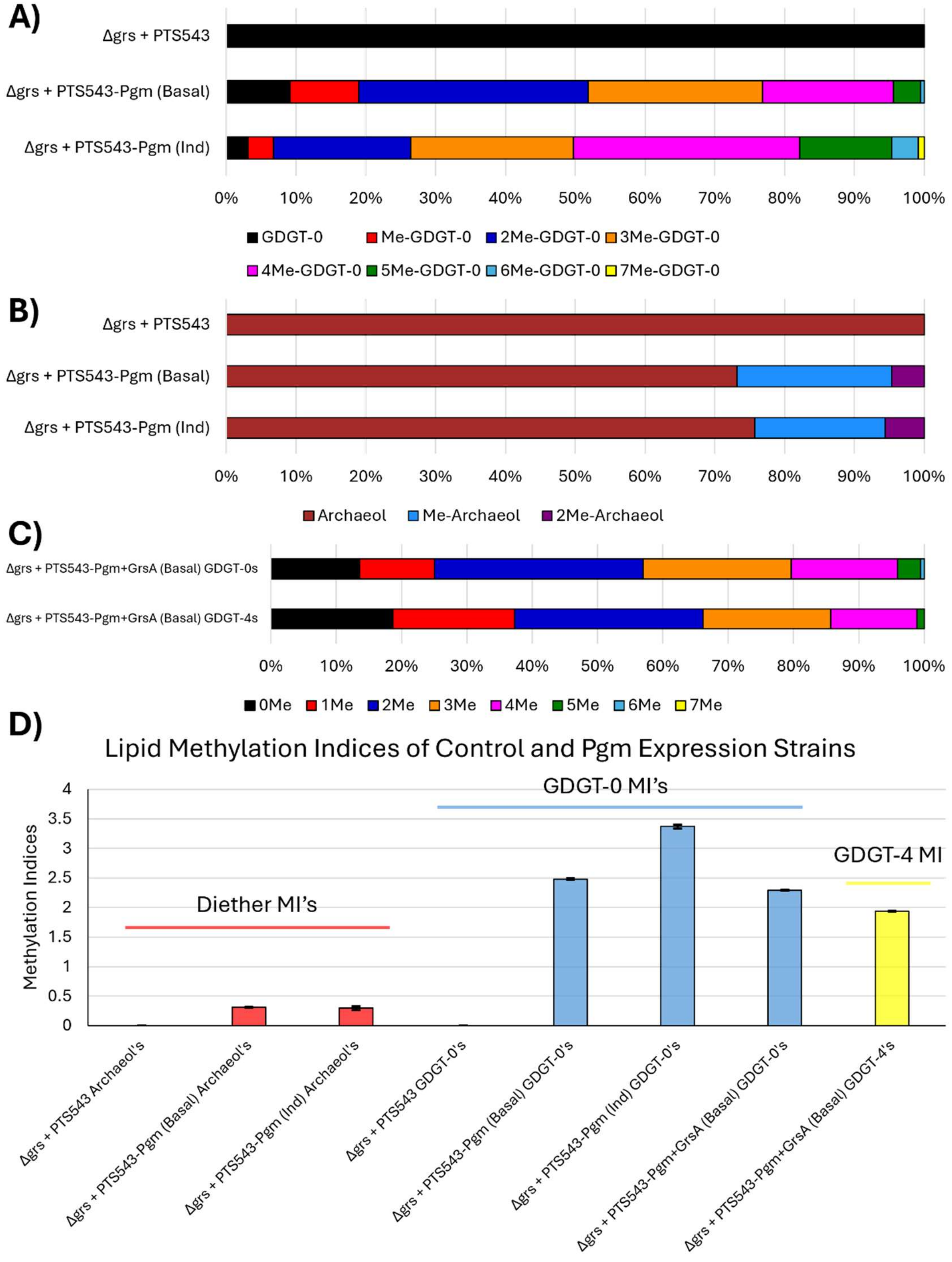


**Fig. S10 Average relative abundance and methylation indices of core lipid species in *T. kodakarensis* control and Pgm expression strains.** A) Average relative abundance of GDGT lipids in the *T. kodakarensis* empty plasmid control strain (Δ*grs* + pTS543), the Pgm expression strain (Δ*grs* + pTS543-*pgm*) under basal conditions (no dextrin), and the Pgm expression strain under inducing (Ind) conditions (+dextrin). The empty plasmid control only makes GDGT-0 and no methylated lipids. In contrast, the vast majority of GDGTs in the Pgm expression strain, under both basal and inducing conditions, are methylated (>90%) and are present in their mono-, di-, tri-, and tetra-methylated forms, as well as in their penta-, hexa-, and hepta-methylated forms, albeit in generally minor amounts (except for higher levels of 5-Me-GDGT-0 during inducing conditions). B) Average relative abundance of diether/archaeol lipids in these same *T. kodakarensis* strains. The empty plasmid control only makes archaeol and does not methylate this lipid. During Pgm expression, however, archaeol is mono- and di-methylated to form Me-archaeol and 2Me-archaeol, albeit to a lesser extent than in the GDGT lipids. This methylation of archaeol demonstrates the potential substrate promiscuity of the enzyme. C) Average relative abundance of the different species of GDGT-0 and GDGT-4 in the Pgm + GrsA co-expression strain. Expression of GrsA with Pgm results in the production of both acyclic and cyclized Me-GDGTs. GDGT-0 and GDGT-4 show similar proportions of methylation and have a similar distribution of mono- to tetra-methylated GDGTs suggesting that Pgm works similarly well on both lipids and/or that GrsA can cyclize methylated GDGT-0, giving rise to cyclized Me-GDGTs as well. D) Lipid methylation indices of the control and *pgm* expression strains. The methylation index (MI) of archaeol and GDGT-0 is zero in the control strain. In the *pgm* expression strain, the GDGT MI is high under both basal and induced conditions. The archaeol MI is low compared to the GDGT-0 MI, demonstrating Pgm prefers a GDGT substrate over bilayer lipids. The MI GDGT-0 and GDGT-4 in the *pgm* + *grsA* co-expression strain is similar, further suggesting Pgm works well on both lipids and/or that GrsA can cyclize the methylated GDGT-0 derivatives, also yielding cyclized Me-GDGTs.

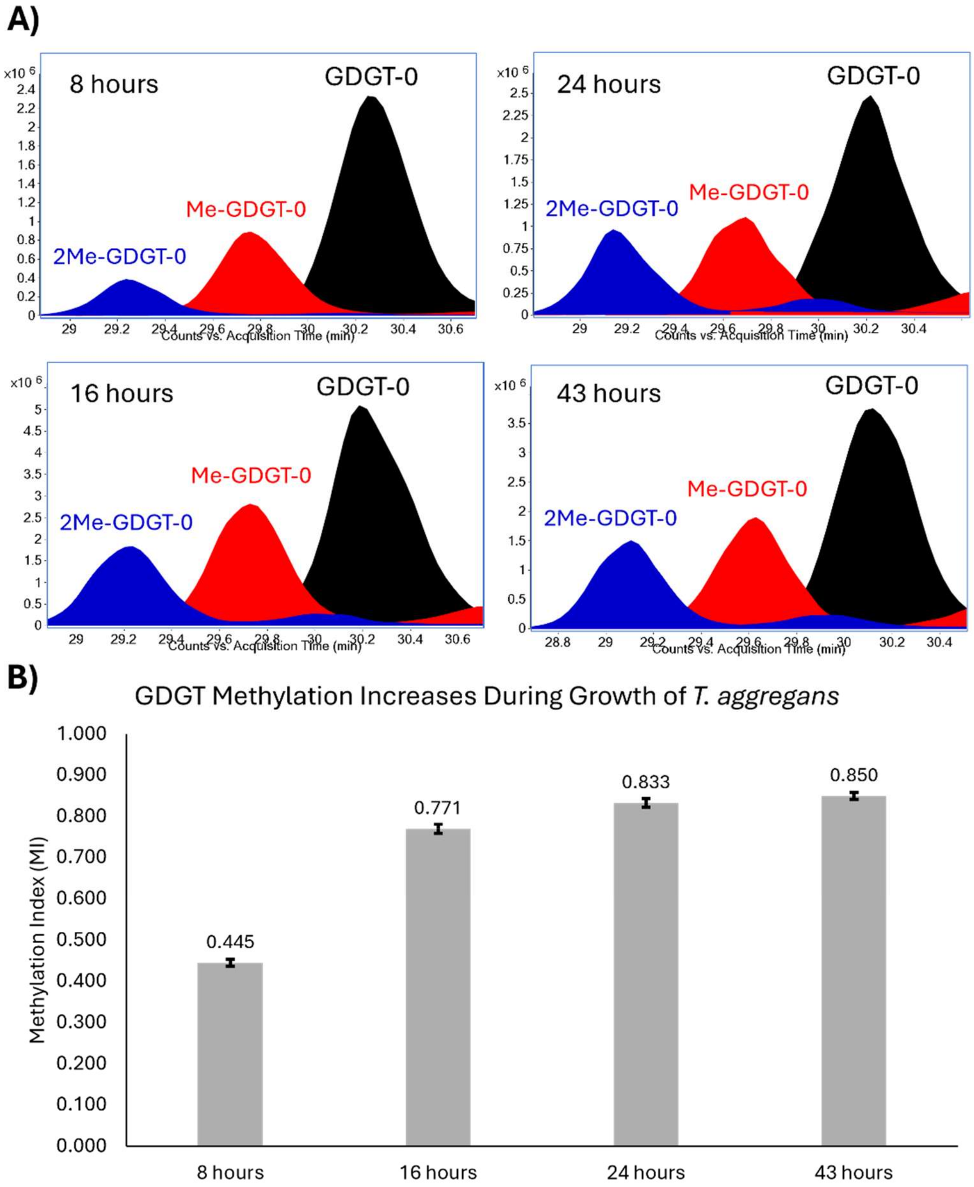


**Fig. S11 *T. aggregans* produces abundant Me-GDGTs, including Me-GDGT-0, which increase throughout the growth phases.** A) Representative, overlayed, extracted ion chromatograms (*m/z* values below*) of acid hydrolyzed biomass from *T. aggregans* showing the production of mono- and di-methylated GDGT-0 during multiple timepoints of growth. Methylated cyclized GDGTs were also detected in similar amounts, but GDGT-0 and its methylated derivatives are shown alone for clarity. B) GDGT methylation index increases throughout the growth of *T. aggregans*, approximately doubling from the first sampling at 8 hours to the last sampling at 43 hours.

* *m/z* = (1302.3, 1324.3; GDGT-0), (1316.3, 1338.3; Me-GDGT-0), (1330.3, 1352.3; 2Me-GDGT-0)

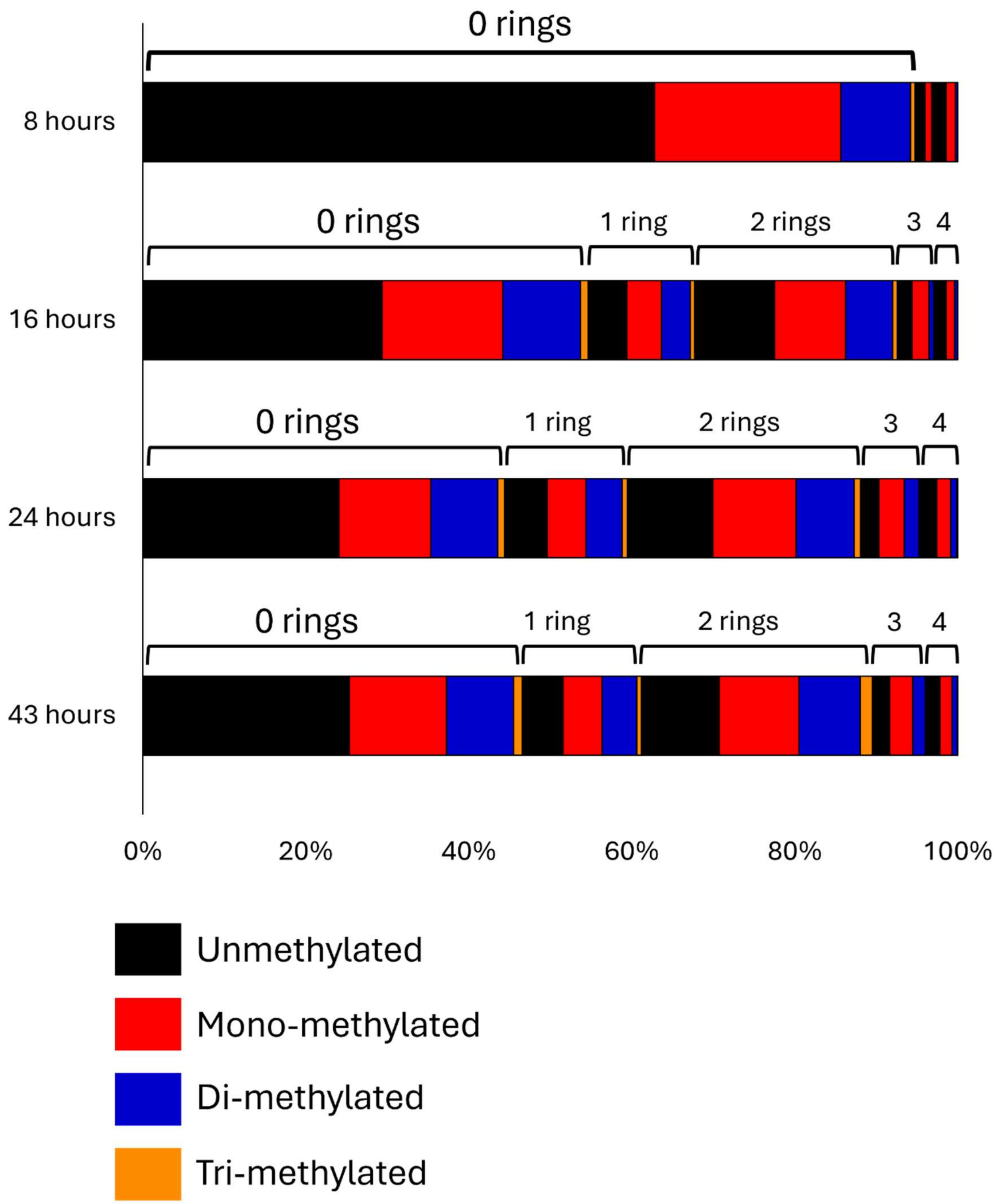


**Fig. S12 Average relative abundance of core lipid species in *T. aggregans* at four timepoints during growth.** Mono-, di-, and tri-methylated derivatives are observed for GDGT-0, 1, 2, 3, and 4. Methylation levels are similar between GDGT-0 and the different cyclized GDGTs, but there is a modest level of enrichment in methylation on the moderately cyclized GDGTs-1 and 2 compared to GDGT-0, GDGT-3, and GDGT-4.

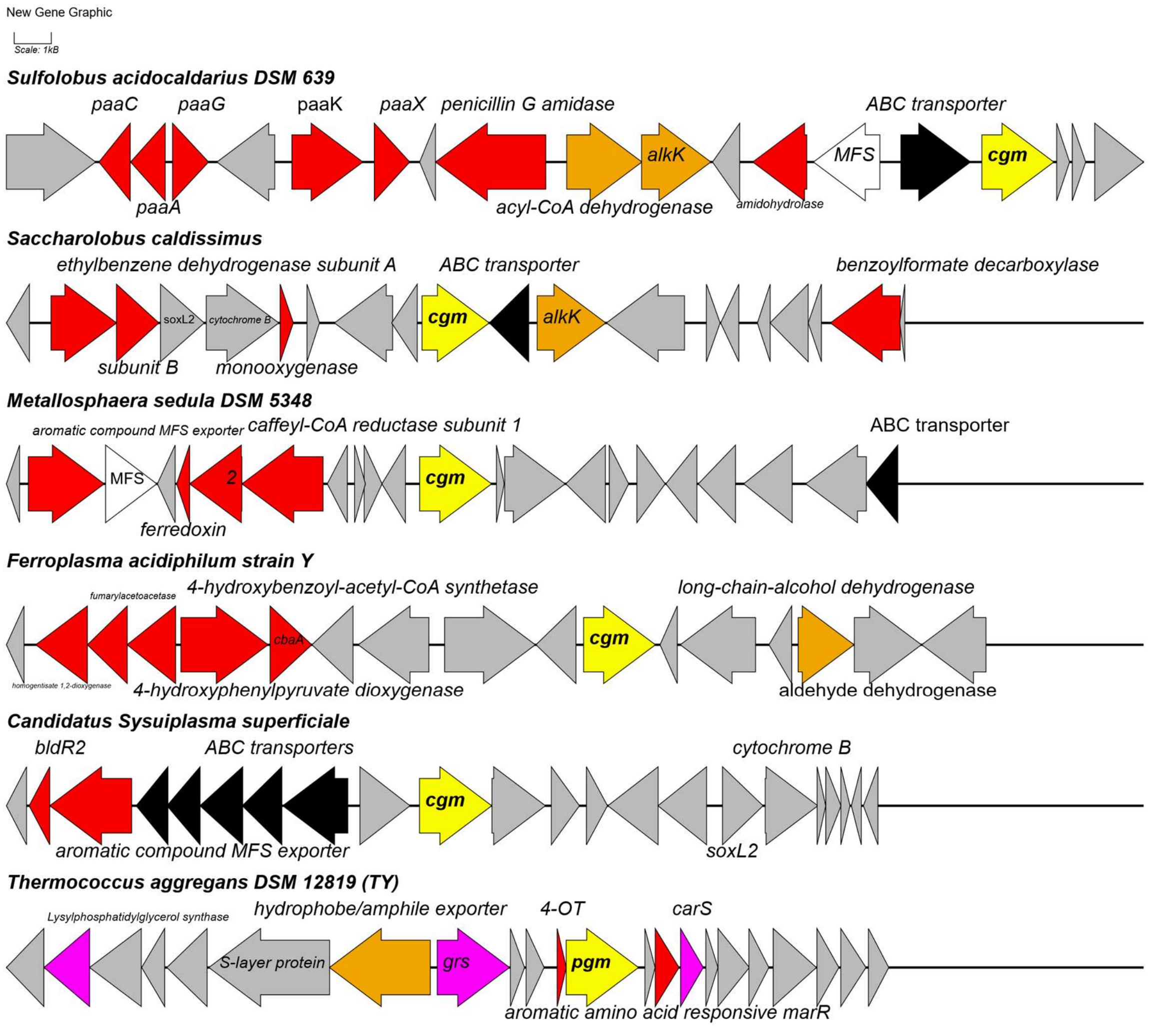


**Fig. S13 GDGT methylase genes of diverse archaea are co-localized with genes involved in the degradation and transport of aromatic compounds and fatty acids.** Representative genome neighborhood diagrams of *cgm* and *pgm* genes (yellow) in archaea, including *S. acidocaldarius* (top) and *T. aggregans* (bottom). Genes involved in aromatic compound degradation, transport, or transcriptional responses are shown in red and genes involved in fatty acid degradation or transport are shown in orange.

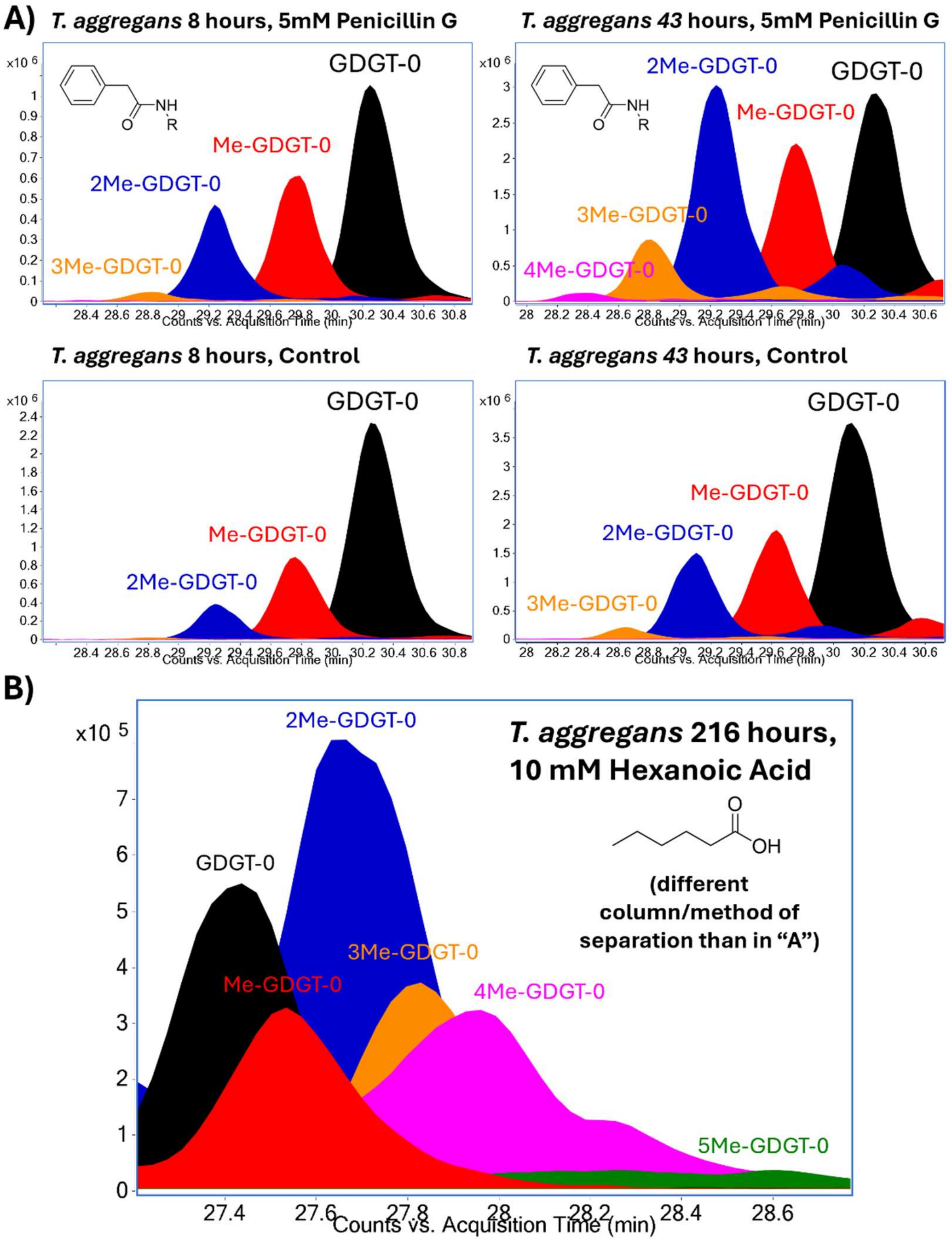


**Fig. S14 GDGT methylation increases in *T. aggregans* in response to exposure to the amphiphiles penicillin G and hexanoic acid.** A) Representative, overlayed, extracted ion chromatograms (EICs; *m/z* values below*) of lipids extracted from acid hydrolyzed biomass of *T. aggregans* grown under standard conditions and in the presence of 5 mM penicillin G. At both 8 hours and 43 hours of growth, GDGT methylation is higher in penicillin G exposed cells as compared to the GDGT methylation during standard growth conditions. GDGT-0 methylation is shown as an example; cyclized GDGTs-1-4 are present in lower individual abundances and possess similar or modestly higher levels of methylation. B) Overlayed EIC (***m/z* values below) of lipids extracted from acid hydrolyzed biomass of *T. aggregans* grown in the presence of 10 mM hexanoic. Lipids were separated on a different column (see Methods) than in "A", resulting in a different elution order of the Me-GDGTs. Growth in the presence of 10 mM hexanoic acid was severely inhibited, and only a single replicate showed growth after 9 days. GDGT methylation was very high in this sample, with robust production of mono-, di-, tri- and tetra-methylated GDGTs. Trace penta-methylated GDGTs (5-Me-GDGT-0) were also detected. GDGT-0 methylation is again shown as an example; cyclized GDGTs and their methylated derivatives are abundant as well.

**m/z* = (1302.3, 1324.3; GDGT-0), (1316.3, 1338.3; Me-GDGT-0), (1330.3, 1352.3; 2Me-GDGT-0), (1344.3, 1366.3; 3Me-GDGT-0), (1358.3, 1370.3; 4Me-GDGT-0)

***m/z* = (1302.3, 1324.3; GDGT-0), (1316.3, 1338.3; Me-GDGT-0), (1330.3, 1352.3; 2Me-GDGT-0), (1344.3, 1366.3; 3Me-GDGT-0), (1358.3, 1380.3; 4Me-GDGT-0) (1372.3, 1394.3; 5Me-GDGT-0)

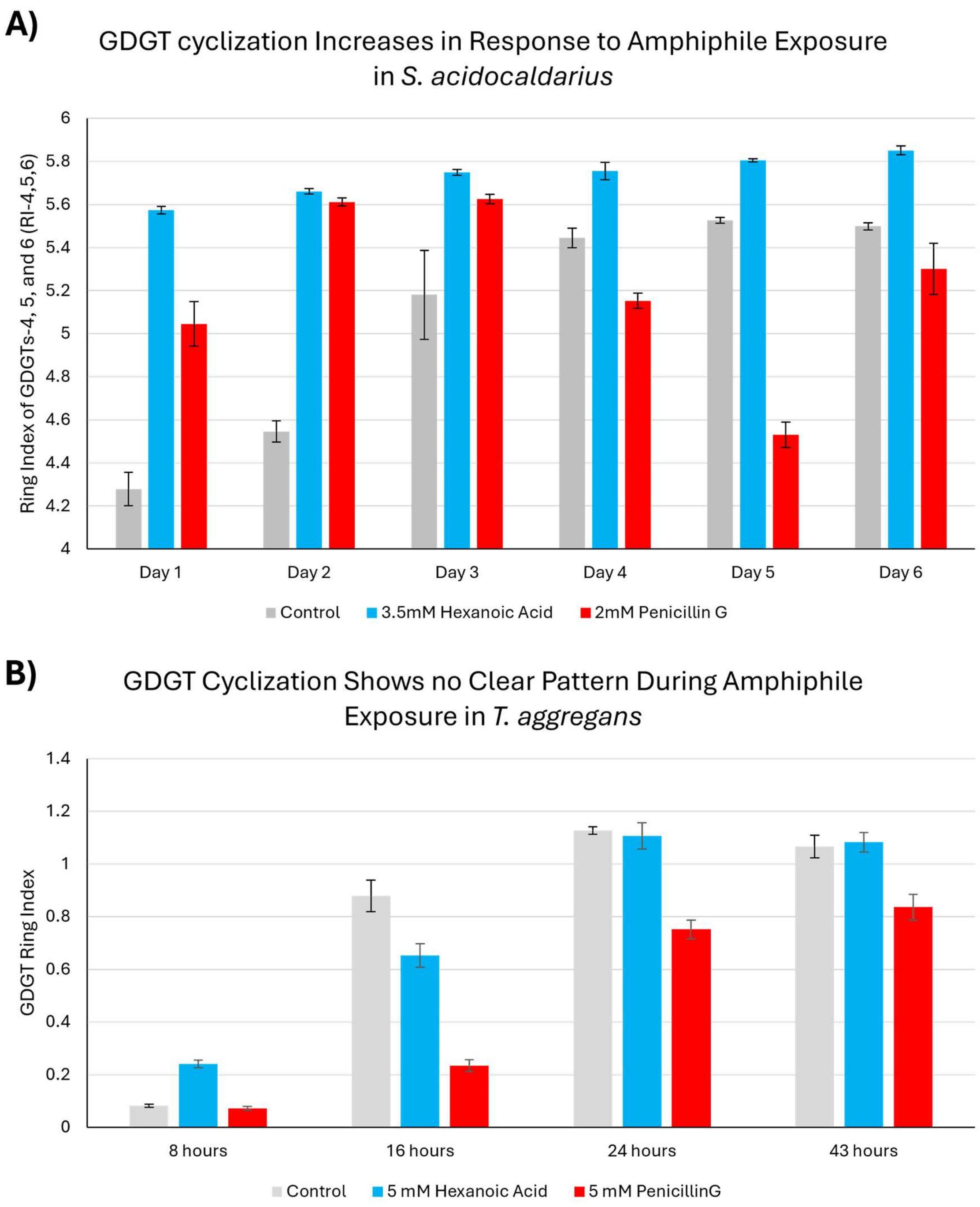


**Fig. S15 Cyclization responses differ during amphiphile exposure in *S. acidocaldarius* and *T. aggregans*.** A) GDGT cyclization increases in response to amphiphile exposure in *S. acidocaldarius*. The degree of cyclization is reported as the ring index of GDGTs-4, 5, 6 which is the weighted average number of rings amongst GDGTs-4, 5, and 6. This measure has a minimum value of 4 and is essentially a measure of highly cyclized GDGT production (GDGT-

5 and 6) by GrsB. This ring index is higher in hexanoic acid exposed cells compared to the control for the full duration of the time course. The ring index is high in penicillin G exposed cells compared to the control as well, but only during days 1-3, similar to the methylation index, and drops significantly in days 4-6 when the growth rate of the cells recovers and penicillin G is presumably degraded. B) GDGT cyclization shows no clear pattern during amphiphile exposure in *T. aggregans*. The degree of cyclization is reported at the ring index of all GDGTs (GDGT-0 to 4) as *T. aggregans* only possesses GrsA and not GrsB and can therefore only synthesize up to four rings. This ring index is only higher in hexanoic acid exposed cells compared to the control during exponential phase. During early stationary phase, late stationary phase, and death phase, the ring index is similar between the control and hexanoic acid exposed cells. The ring index is similar between the penicillin G exposed cells and the control during exponential phase. Alternatively, the ring index is lower in penicillin G exposed cells compared to the control during the other growth stages.

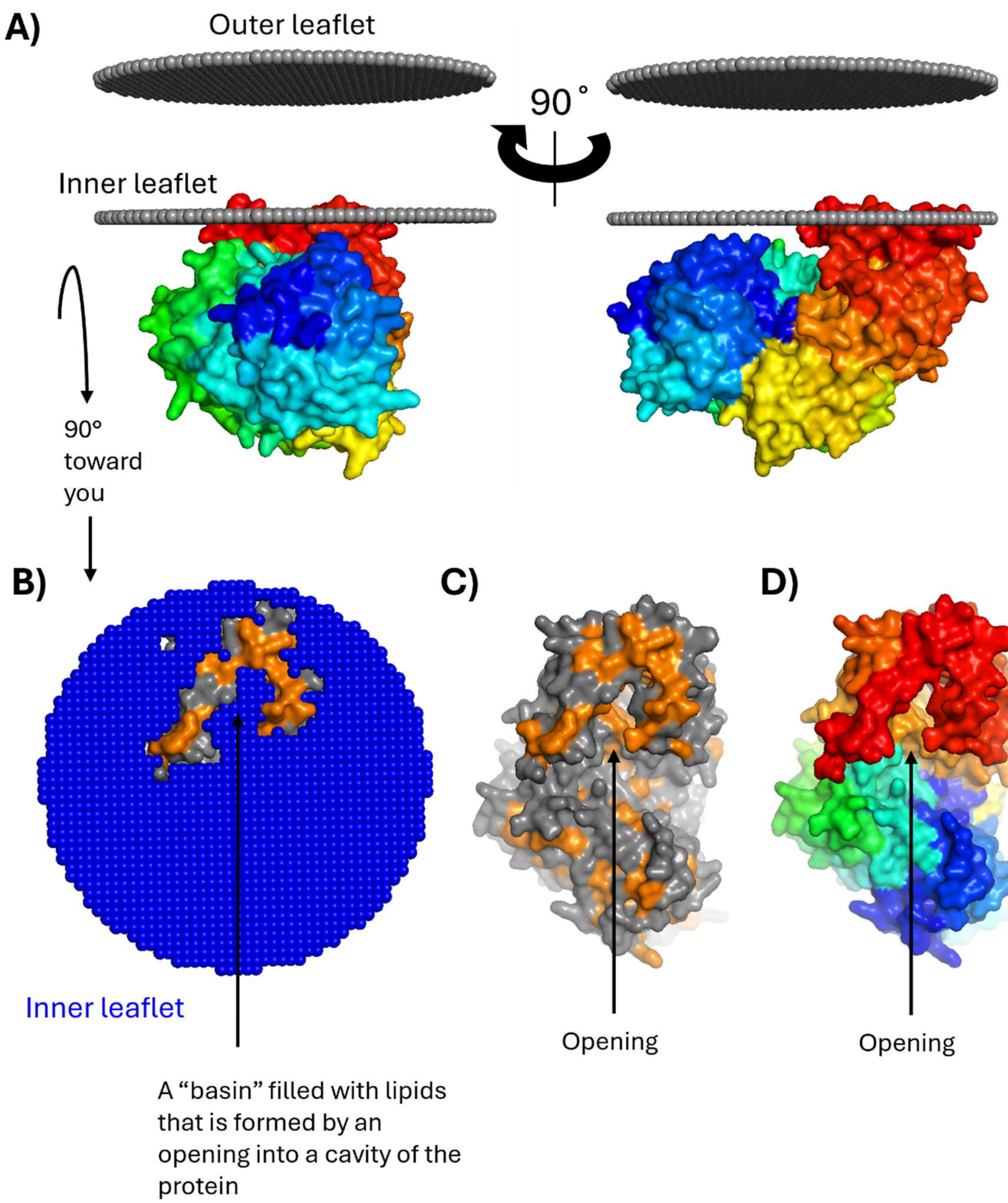


**Fig. S16 A candidate GDGT-modifying rSAM protein from *Candidatus* Bathyarchaeon B1_G15 is predicted to be a peripherally bound membrane protein.** A) An alphafold model (v.3) of the candidate GDGT-modifying rSAM protein from the Bathyarchaeota MAG is positioned peripherally in the cell membrane by the positioning of proteins in membranes (PPM 3.0) webserver. The protein N-terminus is colored blue and proceeds through the rainbow to red at the c-terminus which is partially embedded in the inner leaflet of the cell membrane, similar to Cgm in *S. acidocaldarius*. The protein is modeled to possess a cavity/binding pocket directly beneath the membrane-associated portion of the protein. B) Alphafold model from “A” rotated 90° forward (toward the reader). The outer leaflet is removed to reveal the membrane-embedded portion of the protein. Hydrophobic residues are shown in orange and lipid ends/heads are shown in blue. Much like Cgm in *S. acidocaldarius*, the protein is modeled to possess an opening into a cavity in the protein, forming a “basin” that is filled by the overlying lipids, potentially facilitating substrate binding. C) Same as “B” but the inner leaflet is removed to show the opening into a cavity of the protein. D) Same as “C” but colored as in “A”.

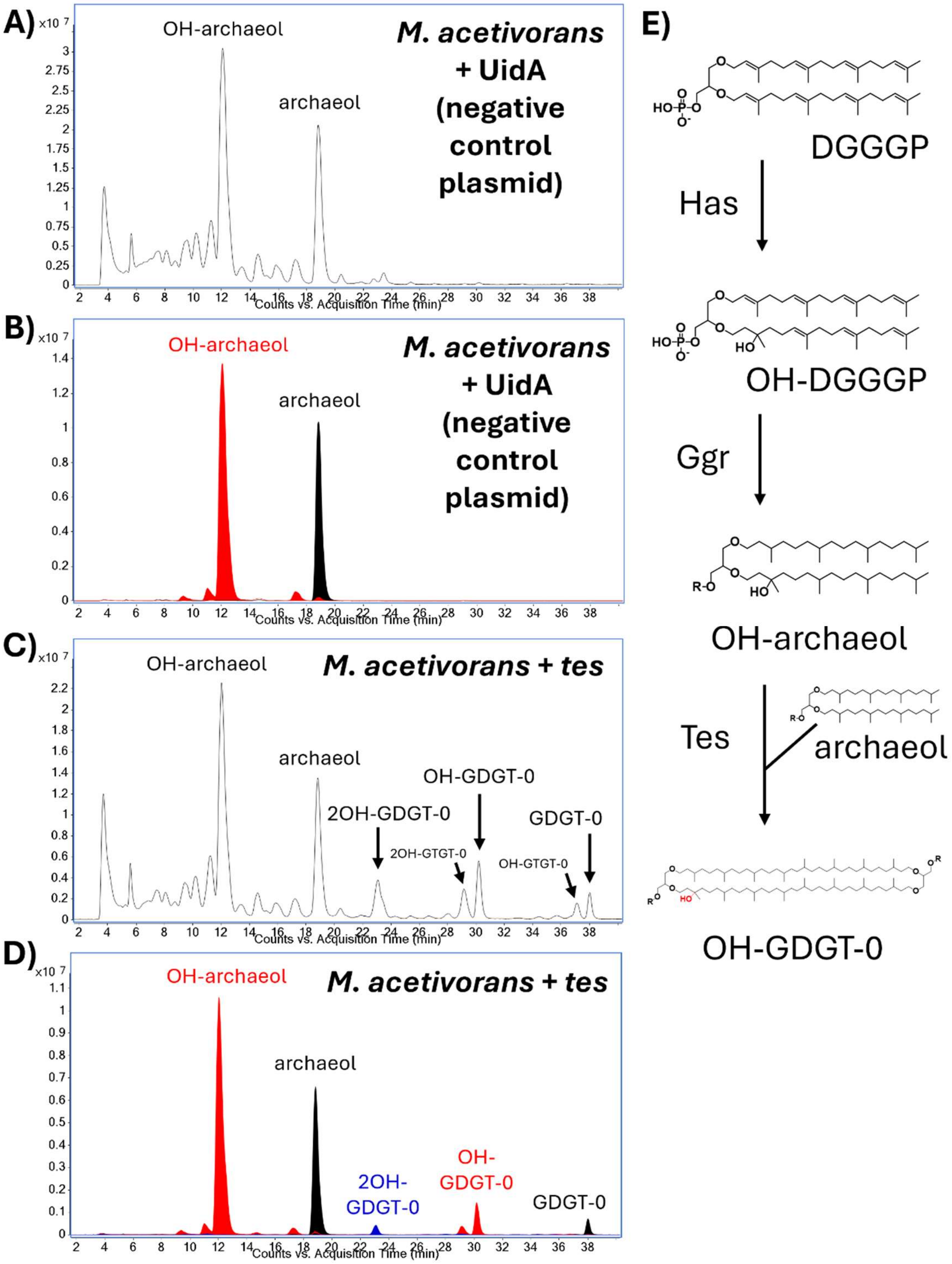


**Fig. S17** ***M. acetivorans*** **natively produces hydroxy-archaeol which is putatively fused with archaeol to form OH-GDGTs during Tes overexpression.** A) Representative total ion chromatogram (TIC) of lipids extracted from base hydrolyzed biomass of the *M. acetivorans*

parental strain carrying a control plasmid . Hydroxy-archaeol (OH-archaeol) and archaeol are the dominant core lipids, as has previously been reported. B) Representative, overlayed, extracted ion chromatogram (EIC; *m/z* values below*) of lipids extracted from base hydrolyzed biomass of the control strain. This EIC highlights the OH-archaeol and archaeol found in the control strain, showing that the peaks formed from extracting the *m/z* values corresponding to their $[M+H]^+$, $[M+NH_4]^+$, $[M+Na]^+$, and $[2M+H]^+$ electrospray ionization adducts correspond to the labeled peaks in the TIC. C) Representative TIC of lipids extracted from base hydrolyzed biomass of *M. acetivorans* overexpressing tetraether synthase (*tes*). During *tes* overexpression we see the production of various tetraether lipid species, including GDGT-0, OH-GDGT-0, and 2OH-GDGT-0. The proportions of OH-GDGTs and GDGTs reflect the proportions of OH-archaeols and archaeols, suggesting the OH-GDGTs are formed when Tes covalently joins OH-archaeol with a non-hydroxylated archaeol, yielding OH-GDGT-0. D) Representative, overlayed, EIC (*m/z* values below*) of lipids extracted from the *tes* overexpression strain, highlighting OH-archaeol, archaeol, 2OH-GDGT-0, OH-GDGT-0, and GDGT-0. E) Putative biosynthesis pathway of hydroxy-GDGTs in *M. acetivorans* and other archaea possessing homologs of the previously characterized *ma0127* encoded phytoene desaturase family hydratase, here termed hydroxy-archaeol synthase (Has). The Has protein hydrates the C2-C3 double bond of the unsaturated intermediates of archaeol biosynthesis such as digeranygeranylglyceryl phosphate (DGGGP) to form OH-DGGGP which is then saturated by the enzyme geranylgeranyl reductase (GGR) to form OH-archaeol. In the presence of Tes, OH-archaeols are dimerized with non-hydroxylated archaeols to form OH-GDGTs or with one another to form 2OH-GDGTs.

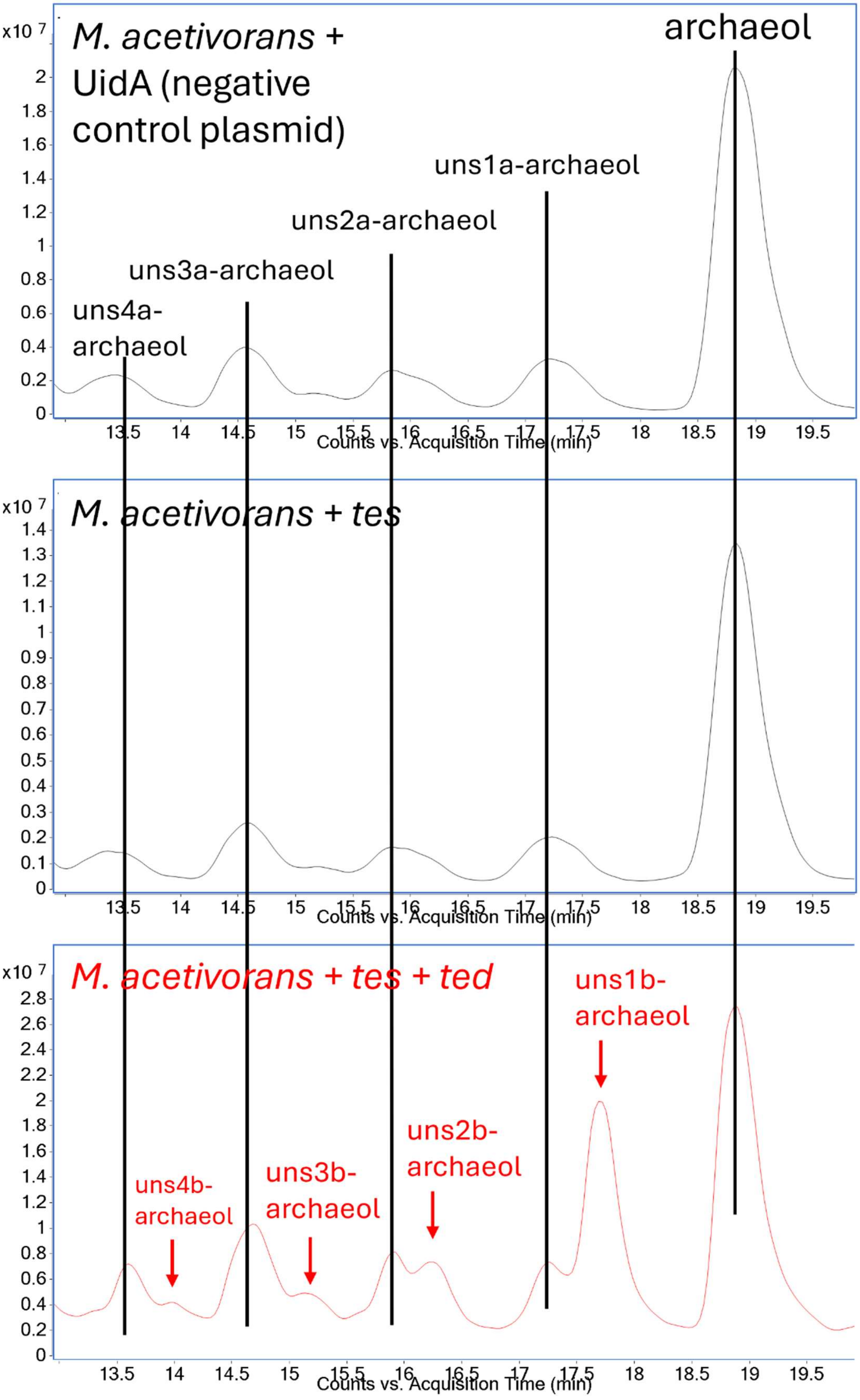
M. acetivorans + UidA (negative control plasmid)
archaeol
uns1a-archaeol
uns2a-archaeol
uns3a-archaeol
uns4a-archaeol
x10 7
Counts vs. Acquisition Time (min)
M. acetivorans + tes
M. acetivorans + tes + ted
uns1b-archaeol
uns2b-archaeol
uns3b-archaeol
uns4b-archaeol

**Fig. S18 Expression of *ted* in *M. acetivorans* results in the production of new unsaturated-archaeol isomers.** Representative total ion chromatograms (TICs) of lipids extracted from base hydrolyzed biomass of *M. acetivorans* with an empty plasmid (control strain), *M. acetivorans* expressing *tes* (*M. acetivorans* + *tes*), and of *M. acetivorans* co-expressing *tes* and *ted* (*M. acetivorans* + *tes* + *ted*). *M. acetivorans* natively possesses low levels of unsaturated archaeol intermediates (e.g. uns1a-archaeol, uns2a-archaeol, uns3a-archaeol, and uns4a-archaeol, with 1, 2, 3, and 4 double bonds, respectively) which are seen in the control strain and the strain expressing *tes*. In the strain co-expressing *tes* and *ted*, we see the production of a new set of unsaturated archaeol isomers (uns1b-, uns2b-, uns3b-, and uns4b-archaeol) in addition to the ones natively observed in *M. acetivorans,* suggesting Ted can desaturate the bilayer lipids as well. However, the GDGTs possess a much higher level of unsaturation than the archaeol lipids, suggesting Ted prefers a tetraether substrate.

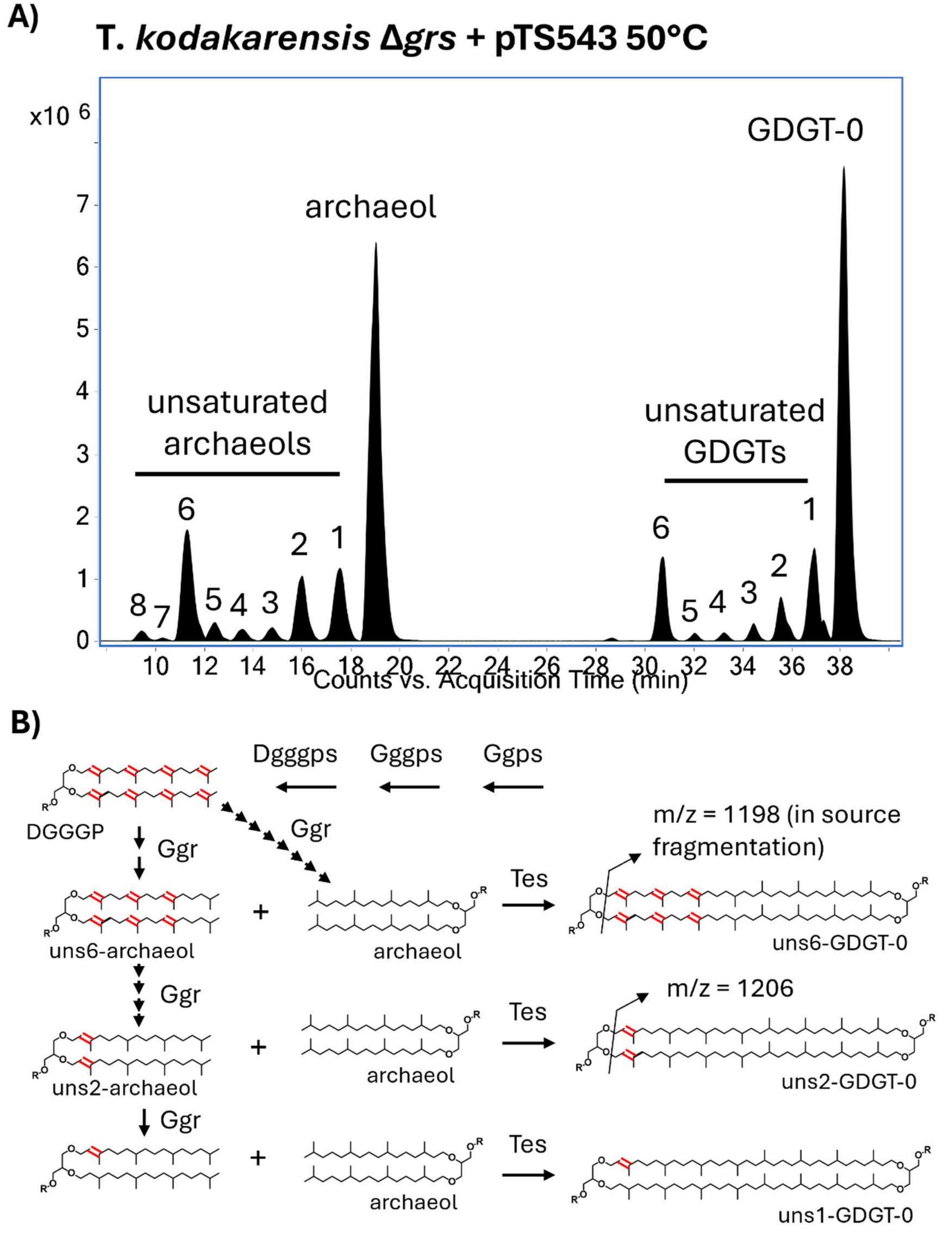


**Fig. S19 *T. kodakarensis* natively possesses unsaturated GDGTs.** A) Representative, overlayed extraction ion chromatograms (EICs) of lipids extracted from base hydrolyzed biomass of the parental *T. kodakarensis* strain (Δ*grs*) carrying an empty plasmid and grown at 50°C. Both unsaturated archaeols and unsaturated GDGTs are seen in the parental *T. kodakarensis* strain. The distribution of the different unsaturated GDGTs is similar to that in the unsaturated archaeols, with both lipid species mainly possessing 1, 2, or 6 double bonds. This suggests the unsaturated GDGTs are formed from the dimerization of an unsaturated archaeol intermediate with a fully saturated archaeol by Tes. B) Putative biosynthetic pathway leading to the production of unsaturated GDGTs in *T. kodakarensis*. DGGGP (uns8-archaeol) is produced by the enzymes Ggps, Gggps, and Dggps which synthesis the C20 isoprene tail and catalyze the formation of the first and second ether bond to glycerol, respectively. DGGGP is subsequently saturated by the enzyme geranylgeranyl reductase (Ggr) producing saturated archaeol as the final product. A variety of unsaturated archaeols possessing 1-7 double bonds are also produced as intermediates by Ggr. These unsaturated intermediates may putatively be covalently joined to a saturated archaeol by tetraether synthase (Tes) to form unsaturated GDGTs. These unsaturated GDGTs exhibit in-source fragmentation during MS analysis due to the aryl nature of the double bonds proximal to the glycerol backbone which facilitate the loss of the glycerol moiety.

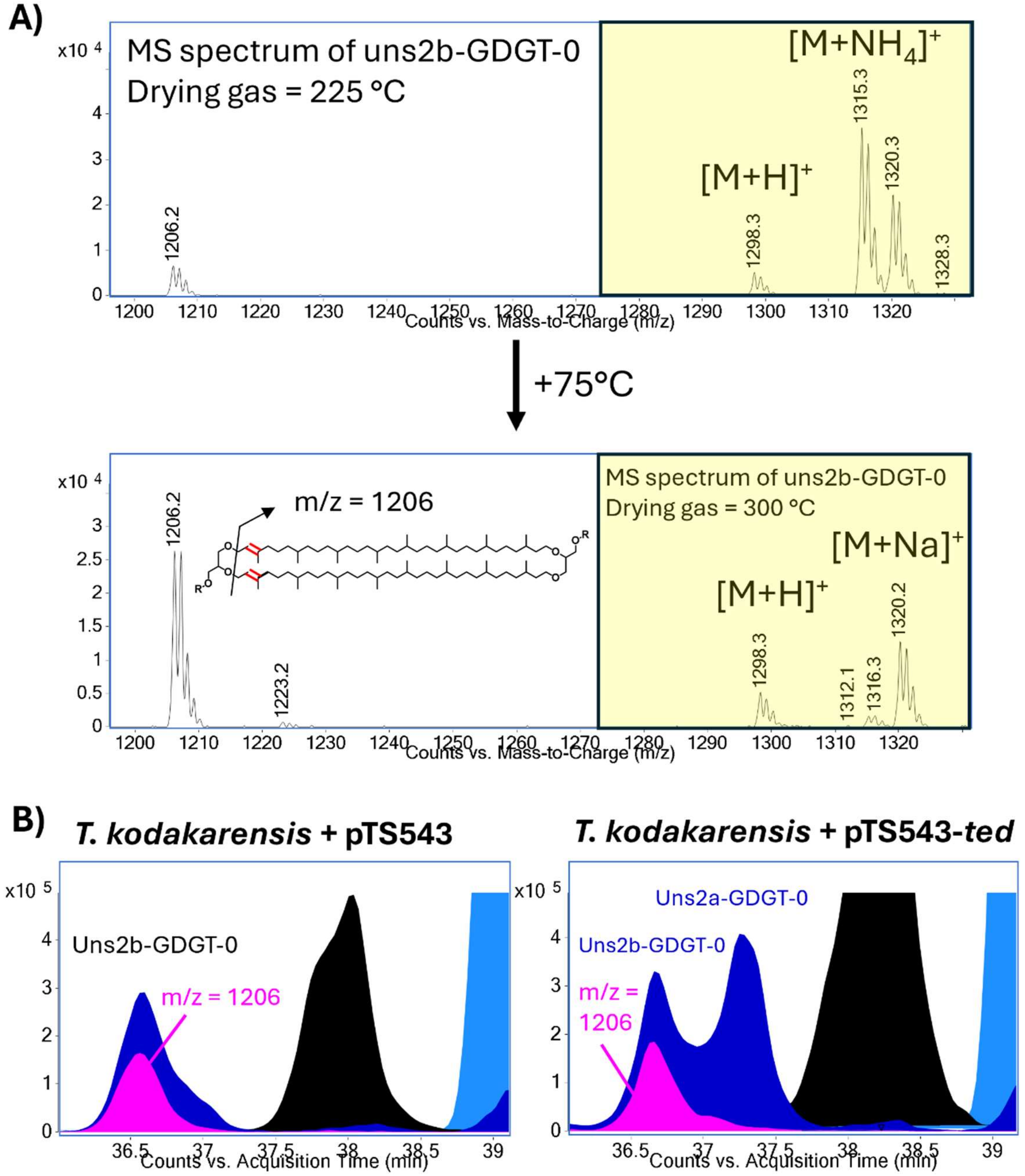


**Fig. S20 The unsaturated GDGTs natively found in *T. kodakarensis* undergo thermally induced in-source fragmentation, resulting in loss of the glycerol backbone.** A) Mass spectrum of the peak corresponding to the native di-unsaturated GDGT found in *T. kodakarensis* (termed uns2b-GDGT-0), shown at two drying gas temperatures. The intact parent ion (M) adducts are shown in the yellow box. At lower drying gas temperatures, uns2b-GDGT-0 is found mainly in its intact form, primarily occurring as ammoniated and sodiated adduct ions at *m/z* = 1315.3 and 1320.3. At higher drying gas temperatures, fragmentation of uns2b-GDGT-0 increases, producing a characteristic ion at *m/z* = 1206.2 which corresponds to a GDGT molecule that has lost a glycerol moiety. This suggests that both double bonds are located on the same end of the GDGT molecule and that they are proximal to the glycerol group, facilitating its cleavage. This *m/z* = 1206 fragment ion is the dominant ion produced by uns2b-GDGT-0 at the higher drying gas temperature of 300 °C, being more abundant than the parent ions. B) Representative, extracted ion chromatograms (EICs) of lipids extracted from base hydrolyzed biomass of the control strain (*T. kodakarensis* + pTS543) and the strain expressing *ted* (*T. kodakarensis* + pTS543-*ted*). Expression of *ted* results in the formation of a new di-unsaturated-GDGT (termed uns2a-GDGT-0) which does not produce fragment ions at m/z =1206, suggesting the location of the double bonds differs between the native uns-GDGTs in *T. kodakarensis* and the uns-GDGTs produced by Ted.

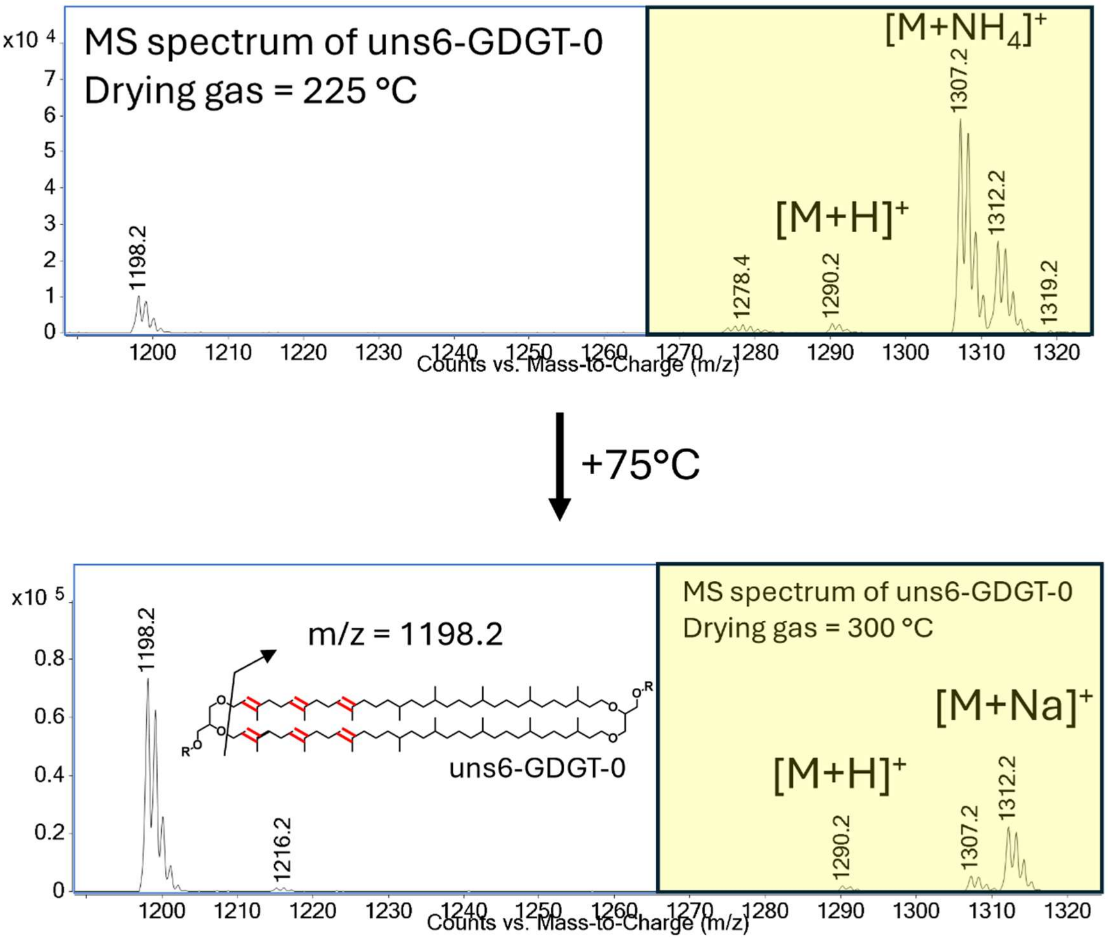


**Fig. S21 The native hexa-unsaturated GDGT-0 (uns6-GDGT-0) found in *T. kodakarensis* undergoes thermally induced in-source fragmentation, resulting in the loss of the glycerol backbone.** This fragmentation produces an ion with an m/z = 1198 and increases with temperature. The intact parent ion (M) adducts are shown in the yellow box.

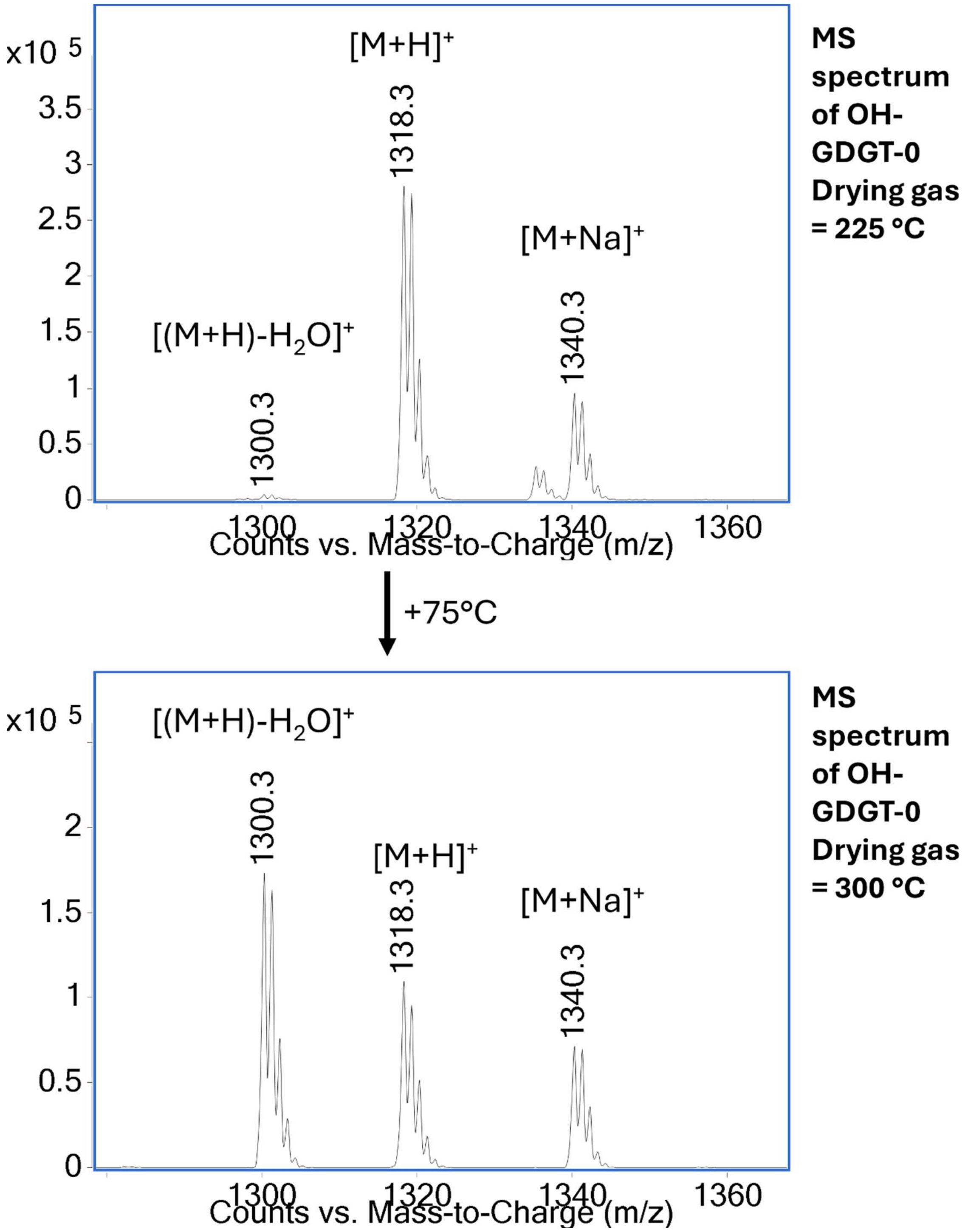


**Fig. S22 The OH-GDGTs produced during *hgs* + *ted* co-expression undergo characteristic, thermally induced, in-source dehydration.** Mass spectrum of the peak corresponding to OH-GDGT-0a in the *ted* + *hgs* co-expression strain. At low drying gas temperatures, OH-GDGT-0 primarily occurs as the protonated and sodiated adducts at m/z = 1318.3 and 1340.4, respectively. When the drying gas temperature is increased by 75°C, this induces the thermal dehydration of OH-GDGT-0, yielding an ion at m/z = 1300.3, 18 mass units less than the parent ion – corresponding to the loss of a water molecule and indicating the compound possesses an additional hydroxyl group compared to “normal” GDGT-0.

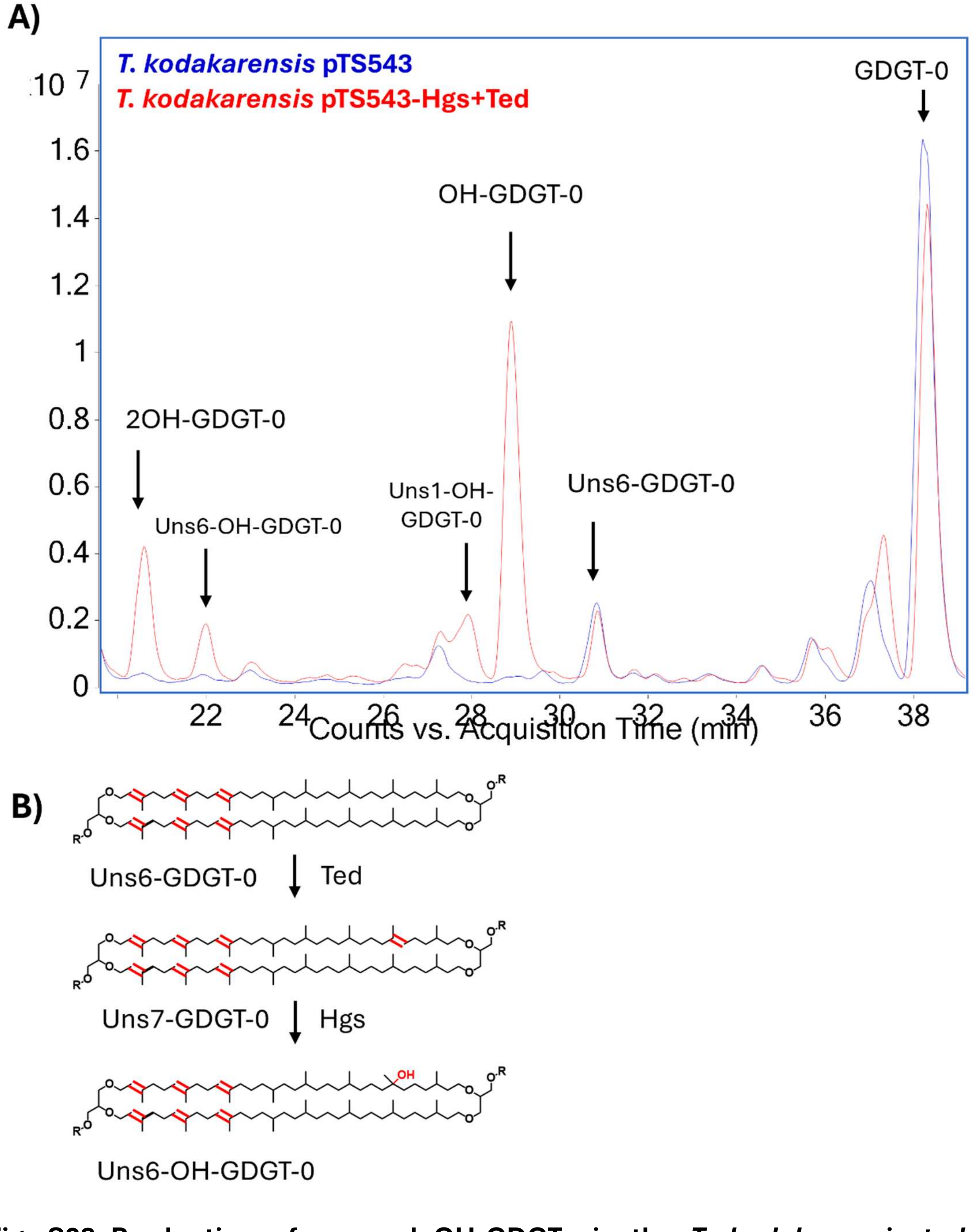


**Fig. S23 Production of unusual OH-GDGTs in the *T. kodakarensis ted* + *hgs* co-expression strain.** A) Representative overlayed total ion chromatograms of lipids extracted from base hydrolyzed biomass of the parental *T. kodakarensis* Δ*grs* strain carrying an empty plasmid and the strain co-expressing *ted* and *hgs*. A series of OH-GDGTs are produced in the co-expression strain including OH-GDGT-0 and 2OH-GDGT-0, putatively resulting from hydration by Hgs of the double bond(s) introduced by Ted. A series of unsaturated mono-hydroxylated OH-GDGTs are also detected, primarily uns1-OH-GDGT-0 and uns6-OH-GDGT-0, likely resulting from the hydration of double bonds that Ted introduced on the native unsaturated GDGTs of *T. kodakarensis*. B) Putative biosynthetic pathway leading to the production of uns6-OH-GDGT-0 in the *ted* + *hgs* co-expression strain. Uns7-GDGT-0 is formed by Ted catalyzed desaturation of the natively produced uns6-GDGT-0. Uns7-GDGT-0 is then hydrated by Hgs at the double bond introduced by Ted, forming uns6-OH-GDGT-0.

**Table S1. Positioning of Proteins in Membranes webserver predicts that Saci_1785 is a peripherally bound membrane protein.** The top 5 Alphafold[2] models (v.3) of Saci_1785 were input into the PPM3.0[4] webserver. All 5 models were predicted to be membrane bound at the C-terminus with the same embedded residues. All models had robust[3] and similar $\Delta G_{transfer}$ values as well.

| Alphafold Model | $\Delta G_{transfer}$ (kcal/mol) | Depth/Hydrophobic Thickness | Embedded Residues |
|---|---|---|---|
| 0 | -14.5 | 4.6 ± 0.9 Å | 424,427,431,442,445-446,448,476,479-481,484,487-488,491-492,495 |
| 1 | -15.2 | 4.9 ± 1.1 Å | 424,427,431,438,442,445-446,448,476,479-481,484,487-488,491-492,495 |
| 2 | -14.2 | 6.0 ± 1.4 Å | 424,427,431,438,441-442,445-446,448,472,476-477,479-485,487-488,491-492,495 |
| 3 | -15.5 | 4.6 ± 1.1 Å | 424,427,431,442,445-446,448,476,479-481,484,487-488,491-492,495 |
| 4 | -14.8 | 4.2 ± 1.0 Å | 424,427,431,442,445-446,448,476,480-481,484,487-488,491-492,495 |

**Table S2. Positioning of Proteins in Membranes webserver predicts that tetraether desaturase (Ted) from *Candidatus Bathyarchaeon* B1_G15 is a peripherally bound membrane protein.** The top 5 Alphafold models (v.3) of Ted were input into the PPM3.0 webserver. All 5 models were predicted to be membrane bound at C-terminus with the same embedded residues. All models had robust[3] and similar $\Delta G_{transfer}$ values as well.

| Alphafold Model | $\Delta G_{transfer}$ (kcal/mol) | Depth/Hydrophobic Thickness | Embedded Residues |
|---|---|---|---|
| 0 | -10.00 | 4.7 ± 0.4 Å | 572-573,576,579,585,588-589,592,607,610-611 |
| 1 | -10.3 | 4.6 ± 0.4 Å | 572-573,576,579,585,588-589,592,607,610-611 |
| 2 | -10.6 | 5.0 ± 1.8 Å | 572-573,576,579,585,588-589,592,607,610-611 |
| 3 | -10.3 | 4.9 ± 2.7 Å | 572-573,576,579,585,588-589,592,607,610-611 |
| 4 | -11.1 | 4.1 ± 0.5 Å | 572-573,576,585,588-589,592,607,610-611 |

**Table S3. Strains used in this study.**

| Strains | | Genotype | Source/Reference |
|---|---|---|---|
| *Escherichia coli* | DH10B | Cloning strain; *F− endA1 recA1 galE15 galK16 nup GrpsL ΔlacX74Φ80lacZΔM15 araD139 Δ(ara,leu)7697 mcrA Δ(mrr-hsdRMS-mcrBC)λ−* | D.K. Newman (Caltech) |
| | ER1821 | Plasmid methylating strain; *F -glnV44 e14- (McrA- ) rfbD1 relA1 endA1 spoT1 thi-1 Δ(mcrC-mrr)114::IS10* | New England Biolabs |
| | WM4489 | *M. acetivorans* plasmid construction; mcrA, Δ(mrr, hsdRMS, mcrBC), Φ80(ΔlacM15), ΔlacX74, endA1, recA1, deoR, Δ(ara, leu)7697, araD139, galU, galK, nupG, rpsL, λattB::pAE12(PrhaB::trfA33, ΔoriR6K-cat::frt5) | Kim et al.[5] |
| | WM7750 | *WM4489/pJK027A* | Guss et al.[8] |
| | WM3357 | *WM1788/pAMG40* | Guss et al.[8] |
| | DN35 | *WM4489/pKES11* | Chadwick et al.[12] |
| | DN616 | *WM4489/pGLC157* | This study |
| | DN677 | *WM4489/pGLC177* | This study |
| *Sulfolobus acidocaldarius* | MW2000 | Δ*pyrEF* (uracil auxotroph parental strain) | Prof. Dr. Sonja-Verena Albers |
| | AG001 | MW2000 Δ*saci_1785*; *cgm* deletion | This study |
| | AG002 | MW2000 Δ*saci_1585*; *grsA* deletion | This study |
| | AG003 | MW2000 *Δsaci_0240*; *grsB* deletion | This study |
| | AG004 | MW2000 + psva1561 | This study |
| | AG005 | MW2000 Δ*saci_1785* + psva1561 | This study |
| | AG006 | MW2000 Δ*saci_1585* + psva1561 | This study |
| | AG007 | MW2000 Δ*saci_0240* + psva1561 | This study |
| | AG008 | MW2000 + psva1561-*saci_1785* | This study |
| | AG009 | MW2000 Δ*saci_1785* + psva1561-*saci_1785*; *cgm* complementation | This study |
| | AG010 | MW2000 Δ*saci_1585* + psva1561-*saci_1785*; *cgm* overexpression in *grsA* deletion | This study |
| | AG011 | MW2000 Δ*saci_0240* + psva1561-*saci_1785*; *cgm* overexpression in *grsB* deletion | This study |
| *Thermococcus kodakarensis* | AL010 | Parental strain; TS559 ΔTK0167* | Liman et al.[6] |
| | AG012 | AL010 + pTS543 | This study |
| | AG013 | AL010 + pTS543-pfba-*pgm*; co-expresses *pgm* from *T. aggregans* (locus tag: *NF865_RS03100*) | This study |
| | AG014 | AL010 + pTS543-pcsg-*grsA*-pfba-*pgm*; expresses *grsA* from *T. kodakarensis* (locus tag: TK0167) and *pgm* from *T. aggregans* | This study |
| | AG015 | AL010 + pTS543-pcsg-*ted*; expresses *ted* from Guaymas basin metagenome (gene ID: *GBSed_1000338310*) | This study |
| | AG016 | AL010 + pTS543 pcsg-*hgs*; expresses *hgs* from Guaymas basin metagenome (gene ID: *GBSed_1000338311*) | This study |
| | AG017 | AL010 + pTS543 pcsg-*hgs*-rbs-*ted*; co-expresses *hgs* and *ted* from Guaymas basin metagenome | This study |

| | | | |
|---|---|---|---|
| *Thermococcus aggregans* | DSM 12819 (TY) | Wildtype | Canganella et al.[7] |
| *Methanosarcina acetivorans* | WWM60 | Δ*hpt*::*PmcrB-tetR* (parental strain) | Guss et al.[8] |
| | DDN351 | WWM60/*pKES11* | This study |
| | DDN398 | WWM60/pGLC157; *tes* (locus tag: MA_1486) overexpression | This study |
| | DDN446 | WWM60/pGLC177; co-expression of *ted* from *Candidatus Bathyarchaeon* B1_G15 (locus tag: DRO69_02625) and *tes* (locus tag: MA_1486)from *M. acetivorans* | This study |

**Table S4. Plasmids used in this study.**

| Plasmids | Description | Source/Reference |
|---|---|---|
| psva407 | *S. acidocaldarius* non-replicative deletion plasmid | Wagner et al.[9] |
| psva1561 | *S. acidocaldarius* replicative expression plasmid | Hoffmann et al.[10] |
| psva407-*saci_1785*UD | *Saci_1785* (*cgm*) deletion plasmid | This study |
| psva407-*saci_1585*UD | *Saci_1585* (*grsA*) deletion plasmid | This study |
| psva407-*saci_0240*UD | *Saci_0240* (*grsB*) deletion plasmid | This study |
| psva1561-*saci_1785* | *Saci_1785* (*cgm*) expression plasmid | This study |
| pTS543 | *T. kodakarensis* replicative expression plasmid | Santangelo et al.[11] |
| pTS543-pfba-*pgm* | *Pgm* (locus tag: *NF865_RS03100*) expression plasmid | This study |
| pTS543-pcsg-*grsA*-pcsg-*pgm* | *GrsA* (locus tag: *TK0167*) and *pgm* co-expression plasmid | This study |
| pTS543-pcsg-*ted* | *Ted* (gene ID: *GBSed_1000338310*) expression plasmid | This study |
| pTS543-pcsg-*hgs* | *Hgs* (gene ID: *GBSed_1000338311* expression plasmid) | This study |
| pTS543-pcsg-*hgs*-rbs-*ted* | *Hgs* and *ted* co-expression plasmid | This study |
| pAMG40 | *Vector for fosmid retrofitting that contains pC2A and λattB* | Guss et al.[8] |
| pJK027A | *Vector with PmcrB(tetO1) promoter fusion to uidA that contains ϕC31-attB and λattP* | Guss et al.[8] |
| pKES11 | Overexpression of UidA (Beta-glucuronidase). Tetracycline inducible. | Chadwick et al.[12] |

| | | |
|---|---|---|
| pGLC157 | T*es* (locus tag: MA_1486) overexpression plasmid. Tetracycline inducible. | This study |
| pGLC177 | Ted (locus tag: *DRO69_02625*) and *tes* (locus tag: MA_1486) expression plasmid. Tetracycline inducible. | This study |

**Table S5. Primers used in this study.**

| Primer Name | Sequence (5' to 3') | Description |
|---|---|---|
| 1785F1 | GTAGGGCCCTGGTATCATTAGTTACTATTGAACTC | *saci_1785* upstream cloning forward and introducing ApaI cut site (underlined) |
| 1785R1 | GATGAAACCAAAATTTGCGTTTTAATCTCTTAATATTAATAC | *saci_1785* upstream cloning reverse and introducing downstream overlapping region (underlined) |
| 1785F2 | GAGATTAAAACGCAAATTTTGGTTTCATCATATCCATGATTTGGG | *saci_1785* downstream cloning forward and introducing upstream overlapping region (underlined) |
| 1785R2 | GTCGGATCCCTCTTTCGAGTTCCACGAGAACTTC | *saci_1785* downstream cloning reverse and introducing BamHI cut site (underlined) |
| Fwd_1785_Checking | GCATTAAAGTTCCTTTTAGCCAG | *saci_1785* deletion check forward |
| Rv_1785_Checking | GACAGGAATAGTTGTTTTGCAG | *saci_1785* deletion check reverse |
| 1585F1 | GTAGGGCCCAGATAACAAGGATATTCTAAGGTG | *saci_1585* upstream cloning forward and introducing ApaI cut site (underlined) |
| 1585R1 | TCTTAAGGTACTGTCAAGAAATCACCTTAACTTAATTG | *saci_1585* upstream cloning reverse and introducing downstream overlapping region (underlined) |
| 1585F2 | TGATTTCTTGACAGTACCTTAAGAAGTAGTATTAGGC | *saci_1585* downstream cloning forward and introducing upstream overlapping region (underlined) |
| 1585R2 | GTCGGATCCGGAGAGGTAATTAAGAGTGTTATAG | *saci_1585* downstream cloning reverse and introducing BamHI cut site (underlined) |
| Fwd_1585_Checking | CATCAATATATGGTGATTTTTGAATG | *saci_1585* deletion check forward |
| Rv_1585_Checking | ACTTTACGCTATTGATACAATAAC | *saci_1585* deletion check reverse |
| 0240F1 | GTAGGGCCCCATGGTTAGTCTCTCTCTATAC | *saci_0240* upstream cloning forward and introducing ApaI cut site (underlined) |
| 0240R1 | TATATTGATTTCCATACATTAAAAAGAGAATCGTATTC | *saci_0240* upstream cloning reverse and introducing downstream overlapping region (underlined) |
| 0240F2 | TTTTAATGTATGGAAATCAATATATTGGCTCATCAG | *saci_0240* downstream cloning forward and introducing upstream overlapping region (underlined) |
| 0240R2 | GTCGGATCCCAAAATAGCATGCGTAGATTTAG | *saci_0240* downstream cloning reverse and introducing BamHI cut site (underlined) |
| Fwd_0240_Checking | CTTCAAATGACGCCTGGTATG | *saci_0240* deletion check forward |
| Rv_0240_Checking | CCGTATGTTAGTTCGAGCTCT | *saci_0240* deletion check reverse |
| Fwd_BspHI_S1785 | CTAGGTATCATGAAAGCCCTATTGGTGAGACCAACAAATCC | *saci_1785* gene cloning forward and introducing BspHI cut site (underlined) |
| Rv_S1785_NcoI | ACGTCAGCCATGGCATAA**TCACCT**CATTCTGTCTTTATCTTGAAG | *saci_1785* gene cloning reverse and introducing NcoI cut site (underlined) and rbs (bold) to connect to *lacS* gene on psva1561 |
| Fwd_psva1561_check | GCGAGTCAGTGAGCGAGGAAG | psva1561 insert check forward |
| Rv_psva1561_check | GGAAGTACAGGTTCTCGGATCCG | psva1561 insert check reverse |

| | | |
|---|---|---|
| Fwd_pTS543_check | GTTGTCCAGCTCATATGCATCACC | pTS543 insert check forward |
| Rv_pTS543 check | GGCGAATTCTGCAGATATCCATCAC | pTS543 insert check reverse |
| GLC362 | CATTATACGAAGTTATCAAGACATATGAATTTCCTCCTTAATGGACTTCAAAAGATTC | Rev primer to amplify MA1486 for cloning into HindIII/NdeI digested pJK027A |
| GLC363 | AATAAATTAAGGAGGAAATTCAATGAAACAGATAAAGTCTG | Fwd primer to amplify MA1486 for cloning into HindIII/NdeI digested pJK027A |
| GLC371 | AGGAGGAAATTCATATGAAACAGATAAAGTCTG | Fwd primer to amplify MA1486 for cloning into HindIII/NdeI digested pJK027A after DRO69_02625 |
| GLC372 | ATAAATTAAGGAGGAAATTCAATGACGTTCAAAAAAATTGTGC | Rev primer to amplify DRO69_02625 for cloning into HindIII/NdeI digested pJK027A before MA1486 |
| GLC373 | TGTTTCATATGAATTTCCTCCTTCAATCACCCATGAATTTATC | Fwd primer to amplify DRO69_02625 for cloning into HindIII/NdeI digested pJK027A before MA1486 |
